# Major change in swine influenza virus diversity in France owing to emergence and widespread dissemination of a newly introduced H1N2 1C genotype in 2020

**DOI:** 10.1101/2024.07.25.605108

**Authors:** Gautier Richard, Séverine Hervé, Amélie Chastagner, Stéphane Quéguiner, Véronique Beven, Edouard Hirchaud, Nicolas Barbier, Stéphane Gorin, Yannick Blanchard, Gaëlle Simon

**Author notes:** Equal contributions (Gautier Richard, Séverine Hervé, Amélie Chastagner).

## Abstract

Swine influenza A viruses (swIAV) are a major cause of respiratory disease in pigs worldwide, presenting significant economic and health risks. These viruses can reassort, creating new strains with varying pathogenicity and cross-species transmissibility. This study aimed to monitor the genetic and antigenic evolution of swIAV in France from 2019 to 2022. Molecular subtyping revealed a marked increase in H1_av_N2 cases from 2020 onwards, altering the previously stable subtypes’ distribution. Whole-genome sequencing and phylogenetic analyses of H1_av_ (1C) strains identified ten circulating genotypes, including five new genotypes, marked by a significant predominance of the H1_av_N2#E genotype. It was characterized by an HA-1C.2.4, an N2-Gent/84, and internal protein-encoding genes belonging to a newly defined genogroup within the Eurasian avian-like (EA) lineage, the EA-DK subclade. H1_av_N2#E emerged in Brittany, the country’s most pig-dense region, and rapidly became the most frequently detected swIAV genotype across France. This drastic change in the swIAV lineages proportions at a national scale was unprecedented, making H1_av_N2#E a unique case for understanding swIAV evolution and spreading patterns. Phylogenetic analyses suggested an introduction of the H1_av_N2#E genotype from a restricted source, likely originating from Denmark. It spread rapidly with low genetic diversity at the start of the epizootic in 2020, showing increasing diversification in 2021 and 2022, and exhibiting reassortments with other enzootic genotypes. Amino acid sequence alignments of H1_av_N2#E antigenic sites revealed major mutations and deletions compared to vaccine 1C strain (HA-1C.2.2) and previously predominant H1_av_N1 strains (HA-1C.2.1). Antigenic cartography confirmed significant antigenic distances between H1_av_N2#E and other 1C strains, suggesting the new genotype escaped from the swine population preexisting immunity. Epidemiologically, the H1_av_N2#E virus exhibited epizootic hallmarks with more severe clinical outcomes compared to H1_av_N1 viruses. These factors likely contributed to the spread of H1_av_N2#E within the pig population. The rapid rise of H1_av_N2#E highlighted the dynamic nature of swIAV genetic and antigenic diversity, underscoring the importance of adapted surveillance programs to support risk assessment in the event of new outbreaks. This also demonstrate the need to strengthen biosecurity measures when receiving pigs in a herd and to limit trading of swIAV-excreting live swine between European countries.

## Introduction

Swine influenza A viruses (swIAV) are responsible for a common respiratory disease in pigs and infections are very frequent in pig herds worldwide (1, 2). The respiratory syndrome is usually acute, characterized by cough, sneezing, nasal discharge, fever and apathy for a period of five to seven days (3). The disease can persist within a farm, contributing to the porcine respiratory disease complex, and cause major animal health issues and economic impact. SwIAV are of a One Health concern (4) considering their zoonotic potential due in part to their ability to bind α(2,6)-linked sialic acids (SA), the major IAV receptors in humans, and the weak immune cross-protection of human population against swIAV (5, 6). SwIAV also represent a risk for some avian species, especially turkeys, quails and pheasants that also carry α(2,6)-linked SA in addition to α(2,3)-linked SA, the receptors preferentially targeted by avian IAV (7–10). Pigs are susceptible to human and avian IAVs (11, 12). As IAVs have a segmented RNA genome that allows genomic reassortment between strains when a host cell is co-infected, pigs have a high capacity to generate reassortant IAVs, leading them to be described as “mixing vessel” hosts (13, 14). Reassortment may occur between different swIAV strains co-circulating in pig populations, sometimes incorporating one or more alternate genes from human or avian IAVs. New reassortant viruses with genomic segments derived from multiple parental lineages may exhibit increased pathogenicity in pigs and/or intra- or inter-species transmission capacity (13).

IAVs classify into subtypes based on antigenic differences in their two major surface glycoproteins, the hemagglutinin (HA) and the neuraminidase (NA). Three main subtypes, i.e. H1N1, H1N2 and H3N2 circulate in pigs, within which several lineages and whole genome constellations are distinguished by geographical regions according to the history of their emergence (1, 15). In 2016, a phylogeny-based global nomenclature classified the H1 viruses in three main clades (15): clade 1A grouping swIAVs with H1 from the so-called “classical swine lineage”, including the 2009 pandemic H1 virus; clade 1B grouping viruses with a H1 from human seasonal lineages (H1_hu_); and clade 1C grouping viruses with HA from the “Eurasian avian-like” (EA) lineage (H1_av_). In Europe, three main lineages were described as enzootic in the European pig population in the early 2000: the “avian-like swine H1N1” (H1_av_N1; HA-1C, N1) lineage, the “human-like reassortant swine H3N2” (H3N2; H3-1970.1, N2-Gent/84) lineage and the “human-like reassortant swine H1N2” (H1_hu_N2; HA-1B.1, N2-Scotland/94) lineage, all with internal protein-encoding genes (IG, which encompass PB2, PB1, PA, NP, M and NS) from the EA lineage. However, over the last 20 years, some major changes in the genetic diversity of European swIAVs have been reported: the emergence in 2003 in Denmark of a H1_av_N2 reassortant, with a Gent-like N2 and EA internal genes (H1_av_N2), which has replaced the H1_hu_N2 lineage in this country (16) and progressively spread in Germany (17), Spain (18, 19), Italy (20) and Bulgaria (21); the gradual decrease in the frequency of H3N2 virus in several countries (6, 22, 23); the introduction of the 2009 pandemic H1N1 virus which has become the “pandemic-like swine H1N1” (H1N1pdm; HA-1A.3.3.2, N1_pdm_) lineage, the fourth enzootic virus at the European level; the emergence of numerous reassortant viruses containing one or more gene(s) from the H1N1pdm, which were sporadically detected or have been established locally, such as H1_pdm_N2 viruses in UK, Germany, Spain, Italy and Belgium (15, 19, 20, 22, 24).

In a previous study which characterized swIAV genetic diversity in France from 2000 to 2018, we also showed an increase in the number of genotypes identified over time, despite very few H3N2 virus detections (25). However, the most dominant viruses characterized annually from 2010 to 2018 were H1_av_N1 (40-78 %) and H1_hu_N2 (13-37 %) lineages, both with EA internal segments, followed by H1N1pdm (1-12 %) (25). An H1_hu_N2 antigenic variant (H1_hu_N2_Δ146-147_), that emerged in 2012 after having fixed mutations in epitopes and deletions in the receptor binding-site on the HA gene, was detected increasingly until 2014 before being less prevalent amongst H1_hu_N2 in following years. Until 2018, reassortant viruses remained sporadic but not rare. For instance, local H1_av_N1 and H1_hu_N2 viruses reassorted into H1_hu_N1 and H1_av_N2 (25), as well as H1_av_N1 which reassorted with H1N1pdm, giving reassortants with various EA and/or pdm internal genes constellations (6). Contrary to other European countries, only two incursions of the Danish H1_av_N2 virus with EA internal genes were reported in France in 2015, whereas another H1_av_N2 of Denmark origin with H1N1pdm internal genes was identified once in late 2018 (25).

Given the ever-increasing genetic and antigenic diversity of swIAV, it is essential to continue to monitor their evolution in order to be able to conduct risk assessments of influenza infections in pigs, both from an animal health and public health perspective. In this study, we characterized the swIAVs detected in France through passive surveillance from 2019 to 2022 with a particular focus on viruses carrying an HA of the 1C clade. In an exceptional way, a marked epizootic took place in 2020 due to an H1_av_N2 genotype that has drastically changed the frequencies of the swIAV subtypes previously established in France for years. This genotype resembled the Danish H1_av_N2 strains previously identified sporadically in 2015. On the basis of genetic and antigenic data we have reconstructed the evolutionary history of this newly dominant H1_av_N2 virus and evaluated the determinants that would have helped its rapid and widespread dissemination in the pig population in France.

## Results

### Molecular subtyping of swIAVs detected in France in 2019-2022

In 2019, 387 independent respiratory outbreaks were investigated in pig herds in France, a number similar to the mean annual number (405±57) of farms investigated from 2015 to 2018 (26). By contrast, this number has risen sharply in 2020, as 661 outbreaks were investigated. The proportions of outbreaks confirmed to be swIAV positive by M-gene RT-qPCR in 2019 and 2020 (47.3% and 52.6%, respectively) were similar to the average annual proportion of 48.2±3.7% calculated in 2015-2018 (26). Among the 531 cases of swIAV infected herds detected in 2019-2020, 375 (70.6%) swIAV strains were fully HA/NA subtyped by specific RT-qPCRs. In 2019, 120 strains were identified as follows: 90 H1_av_N1 (75%), 13 H1_hu_N2 (10.8%) including one H1_hu_N2_Δ146-147_ variant, 8 H1N1pdm (6.7%) and 9 H1_av_N2 (7.5%) (Figure 1). Compared to the distributions reported from 2015 to 2018 (25), which showed quite stable numbers and proportions in the different H1 lineages over time, the proportion of H1_hu_N2 has halved, the H1_av_N2 proportion tripled, whereas other subtype frequencies remained equivalent. In 2020, 255 virus strains were fully subtyped, i.e. twice more than in previous years when 116 to 150 strains were identified per year. Intriguingly, the proportions of the different subtypes were completely shuffled, with 168 H1_av_N2 (65.9%), 64 H1_av_N1 (25.1%), 11 H1_hu_N2 (4.3%), 9 H1N1pdm (3.5%) and 3 H1_pdm_N2 strains (1.2%).

**Figure 1:**
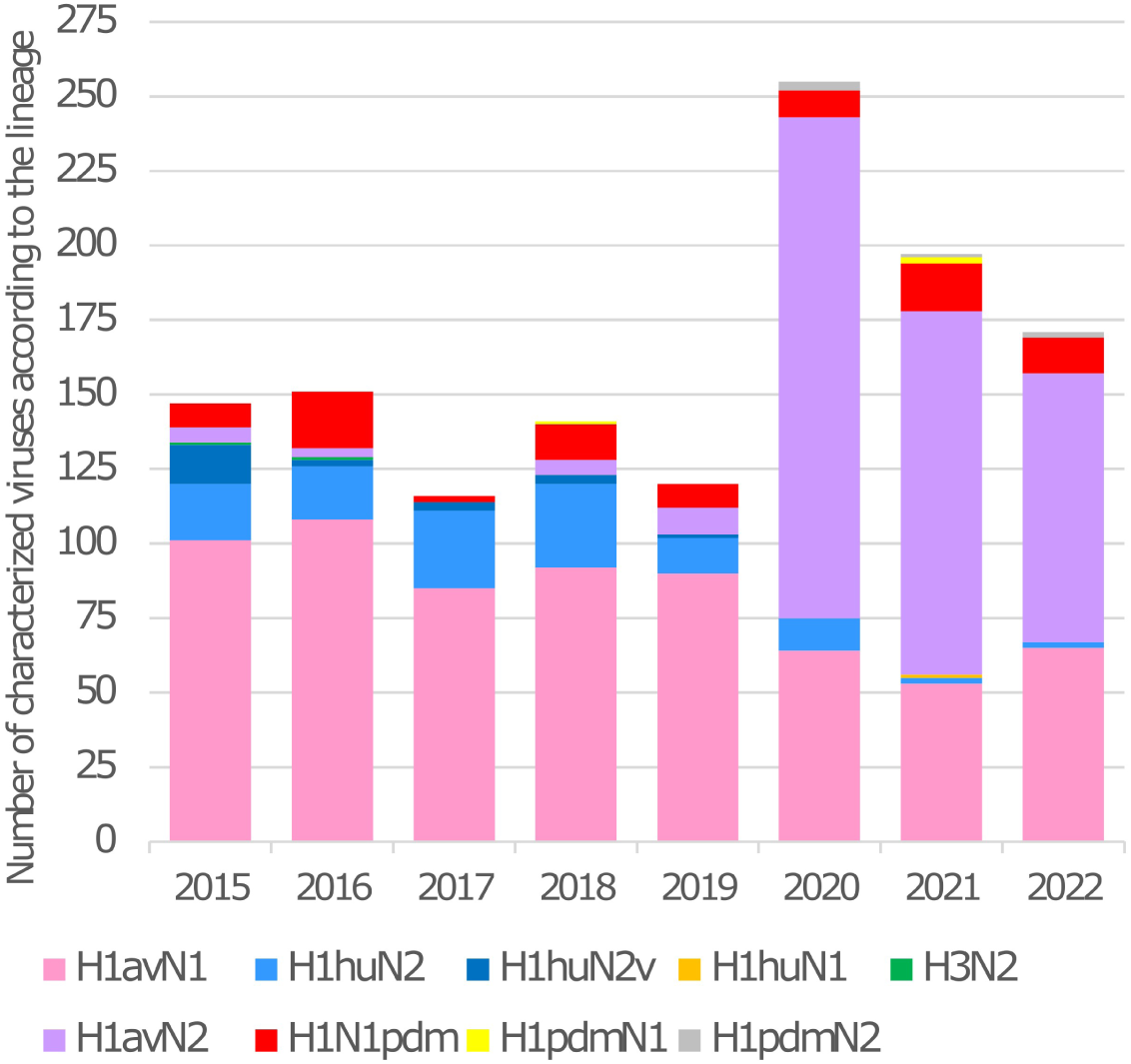
Numbers of swine influenza A virus strains identified in France from 2019 to 2022, per year according to their subtype, and comparison to numbers previously reported from 2015 to 2018 (25).

In 2021, the swIAV passive surveillance reached 727 respiratory outbreaks and 307 (42.2%) positive cases. Among them, 197 swIAV strains were subtyped and distributed as follows: 122 H1_av_N2 (61.9%), 53 H1_av_N1 (26.9%), 16 H1N1pdm (8.1%), 2 H1_hu_N2 (1%), two H1_pdm_N1 (1%), 1 H1_hu_N1 (0.5%) and 1 H1_pdm_N2 (0.5%) (Figure 1). In 2022 the situation was quite similar to 2021 and 2020, with however fewer cases investigated (548 outbreaks, 45.3% of positive cases). The H1_av_N2 remained the most prevalent strain (n=90, 52.6%), followed by 65 H1_av_N1 (37.8%) and 12 H1N1pdm (7%). The H1_hu_N2 stayed low (n=2, 1.2%) and two H1_pdm_N2 (1.2%) were detected.

### Phylogenies, genotyping and spatiotemporal distribution of 2019-2022 1C swIAV

The diversity and phylogeny of these newly emerging and persisting H1_av_N2 viruses were then characterized by monitoring the evolution of swIAV displaying an HA gene of the 1C clade. Swine IAV strains collected from 2019 to 2022 were submitted to whole-genome sequencing. After genotyping, 331 swIAV genome sequences displayed an HA of the 1C clade and were analyzed using Maximum-likelihood phylogenies, when sequence quality permitted, alongside related European sequences that were publicly available at the time of September 2023 (Figure 2). For HA, the clades 1C.2.1, 1C.2.2 or 1C.2.4 were detected, and the N1-EA, N2-Gent/84, N2-Scotland/94 and human seasonal-N2 lineages were detected for the NA segment (Figure 2A). Three lineages were distinguished for all six internal protein-encoding genes (IG): EA, EA-DK and pdm (Figure 2B). The EA-DK lineage was newly established in this study considering its important evolutionary distance compared to the EA lineage for all IG. The name EA-DK was given as sequences from Denmark were predominant in the roots of this new genogroup (Supplementary Figure 1).

**Figure 2:**
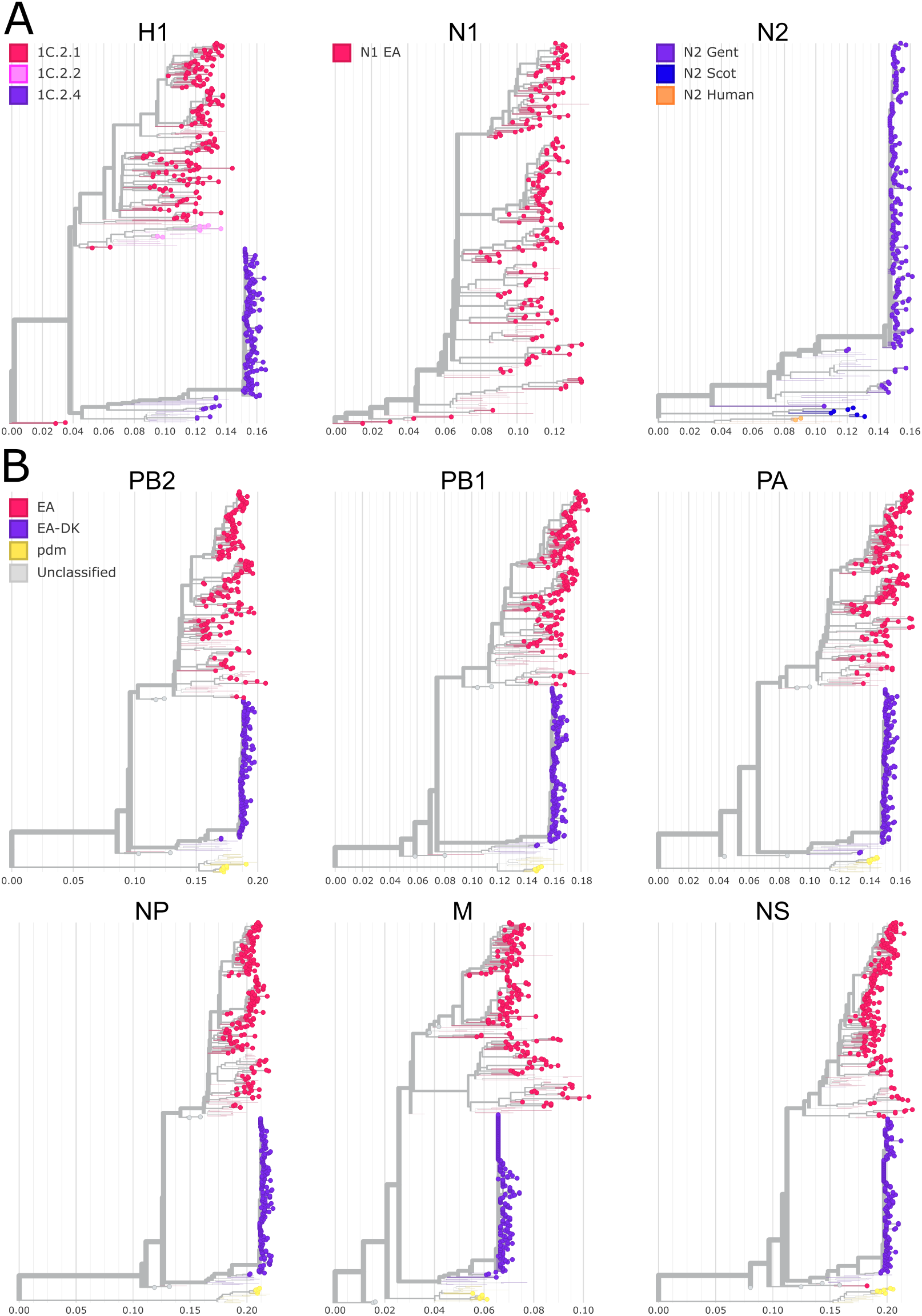
Maximum-likelihood divergence phylogenies of the eight segments of swine influenza A viruses collected and sequenced in France up to 2022 having an HA of the 1C (Eurasian-avian-like) clade, and the related publicly available European sequences. **A**. Surface protein-encoding genes phylogenies (H1, N1 and N2). **B**. Internal protein-encoding gene phylogenies (PB2, PB1, PA, NP, M and NS). Sequences are represented by dots colored according to their individual segment clade: Eurasian avian-like (EA) in pink, Eurasian avian-like Denmark (EA-DK) in purple, Pandemic09 in yellow (pdm). Publicly available European sequences as of September 2023 are displayed as thin branches without dots. Interactive versions of the trees can be found at https://nextstrain.org/community/gtrichard/swIAV-H1av-France-2019-2022

Overall, 10 full-genome genotypes were identified among 1C strains collected between 2019 and 2022 in France (Figure 3A). Regarding H1_av_N1 subtype, four genotypes were distinguished: genotype H1_av_N1#A (HA-1C.2.1, N1-EA, IG-EA) continued to be predominant in 2019 (Figure 3A and 3B) and across the entire country (Figure 3C). Both genotypes H1_av_N1#B (HA-1C.2.1, N1-EA, IG-EA-DK) and H1_av_N1#D (HA-1C.2.1, N1-EA, IG-EA with M-EA-DK), which had never been detected in France before, were sporadic and found only in the western part of France throughout the 2020-2022 period (Figure 3B and 3C). Genotype H1_av_N1#C (HA-1C.2.2, N1-EA, IG-EA) was also sporadically reported at the end of 2019 in the *Hauts-de-France* region and throughout 2022 in *Hauts-de-France* and *Auvergne-Rhône-Alpes* regions (Figure 3B and 3C).

**Figure 3:**
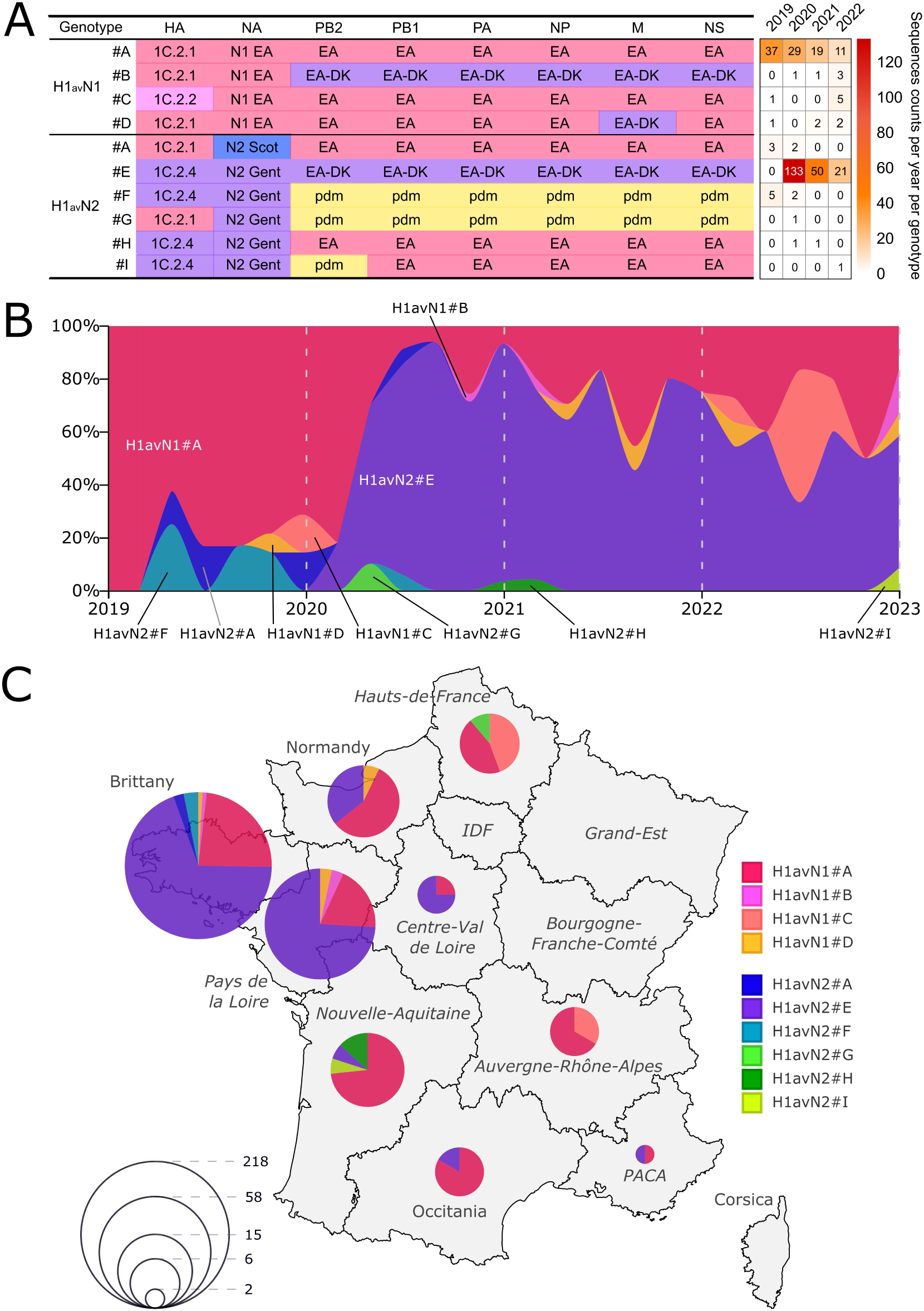
Spatiotemporal distribution of France 1C swIAV genotypes identified in France from 2019 to 2022. **A.** Detail of the gene constellation of the 10 swIAV HA-1C genotypes. The counts of full-genome sequences collected each year is displayed as a heatmap, for each genotype, at the right of the table. **B.** Normalized bimestrial country-wide proportion of the swIAV genotypes. **C.** Region-resolved map of France. Each pie-chart represents the proportions of swIAV genotypes identified in the associated region between 2019 and 2022. The size of the pie-chart is proportional to the log2 number of sequences collected. IDF: *Ile-de-France*, PACA: *Provence-Alpes-Côte-d’Azur*.

Within the H1_av_N2 subtype, six genotypes were identified (Figure 3A). The H1_av_N2#A (HA-1C.2.1, N2-Scotland/94, IG-EA) used to be sporadically detected almost every year since 2008 following a reassortment between local enzootic H1_av_N1#A and H1_hu_N2 viruses (25). However, only five H1_av_N2#A strains were detected in Brittany in 2019 and 2020. The vast majority of the sequenced H1_av_N2 strains belonged to the H1_av_N2#E (HA-1C.2.4, N2-Gent/84, IG-EA-DK) (Figure 3A). This genotype started to be detected sporadically at the beginning of 2020 to rapidly rise as the most detected genotype, especially in Brittany, the French region with the highest density of pig farming. It was then detected in seven out of the nine monitored administrative regions of the country and maintained as the most detected strain for the rest of the study period, thus replacing H1_av_N1#A as the predominant genotype (Figure 3B). The other H1_av_N2 genotypes were sporadic just like the H1_av_N2#A. Five strains belonged to the H1_av_N2#F genotype (HA-1C.2.4, N2-Gent/84, IG-pdm) (Figure 3A) and were reported in Brittany in 2019 and 2020. It was first detected in November 2018 in Brittany (25) and was suggested to be introduced *in toto* from abroad, likely from Denmark considering the HA/NA phylogeny of the newly sequenced H1_av_N2#F strains (Supplementary Figure 2A). Three H1_av_N2 genotypes were detected for the first time during the study period: H1_av_N2#G (HA-1C.2.1, N2-Gent/84, IG-pdm) which corresponded to a single strain detected in March 2020 in the *Hauts-de-France* region; H1_av_N2#H (HA-1C.2.4, N2-Gent/84, IG-EA) which was detected only twice in the *Nouvelle-Aquitaine* region in December 2020 and January 2021; and a single strain similar to H1_av_N2#H but with a PB2 from the pdm lineage, detected in the same region at the end of 2022 and got designated as belonging to the H1_av_N2#I genotype (Figure 3).

### Origins of the newly detected sporadic swIAV genotypes

Throughout this study, five swIAV genotypes were detected for the first time in France and their potential origins were checked through maximum-likelihood HA/NA tanglegram (Figure 4) and whole-genome phylogenies. The H1_av_N1#B and H1_av_N1#D strains, considering that their HA and NA phylogenies were confounded within the H1_av_N1#A phylogenies, were most likely reassortants between this genotype and H1avN2#E from which they acquired one or more EA-DK IG (Figure 4).

**Figure 4:**
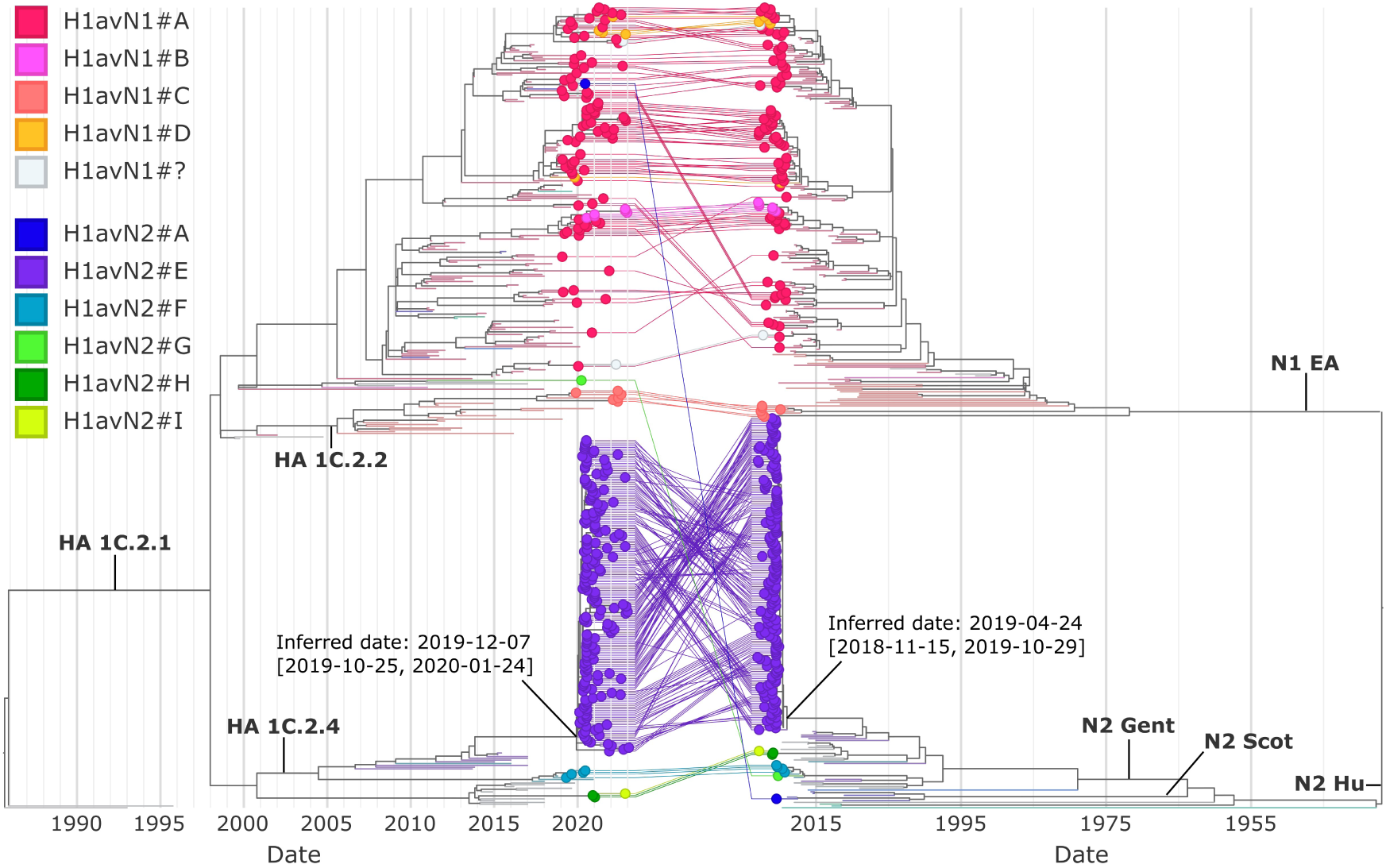
Time-resolved maximum-likelihood phylogenies of the surface protein-encoding genes (left: HA, right: NA) colored by genotypes as presented in Figure 3. Dots represent swIAV sequences collected in France between 2019 and 2022. Branches without dots represent publicly available European 1C swIAV strains as of September 2023, including previous strain sequences collected in France. Central lines connect the HA and NA sequences of the same strains. H1_av_N1 strains with missing internal genes sequences are indicated as H1_av_N1#?. Inferred date and confidence interval is displayed for the H1_av_N2#E 2020-2022 clade common ancestor in both HA and NA trees. An interactive version of the tree can be found at: https://nextstrain.org/community/gtrichard/swIAV-H1av-France-2019-2022/HA:community/gtrichard/swIAV-H1av-France-2019-2022/NA?c=genotype&f_country=France

The H1_av_N2#G strain displayed HA and NA genes closer to Danish strains than French strains (Supplementary Figure 2B) and full-genome alignments showed a proximity to H1_av_N2#F strains from Denmark (Supplementary Figure 3A). This suggested that the H1_av_N2#G strain was likely an H1_av_N2#F reassortant on the HA gene (HA-1C.2.4 was replaced by HA-1C.2.1) introduced *in toto* from abroad, likely from Denmark.

The H1_av_N2#H and H1_av_N2#I strains were not related to any already sequenced strain from France but clustered with sequences found in Denmark and Italy at both the HA/NA and full-genome levels (Supplementary Figures 2C and 3B). This suggested an introduction *in toto* from abroad of the H1_av_N2#H strain. Considering the HA/NA phylogeny, the #I genotype was most likely a reassortant of the #H genotype on the PB2 segment (EA to pdm). The reassortment might have happened in France considering how close the two strains are on the HA and NA genes as well as considering that they were detected sequentially in the same region of the country.

### Origins of the newly predominant H1_av_N2#E swIAV genotype

Genotyping of the 1C viruses revealed that the H1_av_N2#E genotype was solely responsible for the rise of the H1_av_N2 detections during the 2020-2022 period. Such a contrasted and fast switch of the mainly detected swIAV genotypes at the scale of France was unprecedented. This raised the question of the potential origins of these H1_av_N2#E strains.

The H1_av_N2#E genotype formed a single phylogenetic cluster, whose inferred date of the last common ancestor was estimated to be set between October 2019 and January 2020 according to Nextstrain analyses of the HA gene, while the analyses on the NA gene had confidence interval spanning over a year between November 2018 and October 2019 (Figure 4). Before 2020, only two H1_av_N2#E isolates were characterized in the South-Western part of France in 2015 (25). The H1_av_N2#E genotype was thus not detected from 2016 to 2019, until its identification in Brittany in February 2020. HA/NA phylogenies indicated that the H1_av_N2#E 2020 strains detected in France were related to 2017-2018 sequences detected in Europe (Figure 4). Full genome phylogeny of all H1_av_N2#E strains revealed that the strains detected in France in 2020 were closer to strains from Denmark identified from 2015 to 2017 than to the French 2015 strains (Figure 5A). Amino-acid conservation analyses confirmed that most IG of a representative H1_av_N2#E 2020 strain were closer to 2015-2017 strains from Denmark than to the A/swine/France/65-150242/2015 strain, while HA and NA were equally similar (Figure 5B).

**Figure 5:**
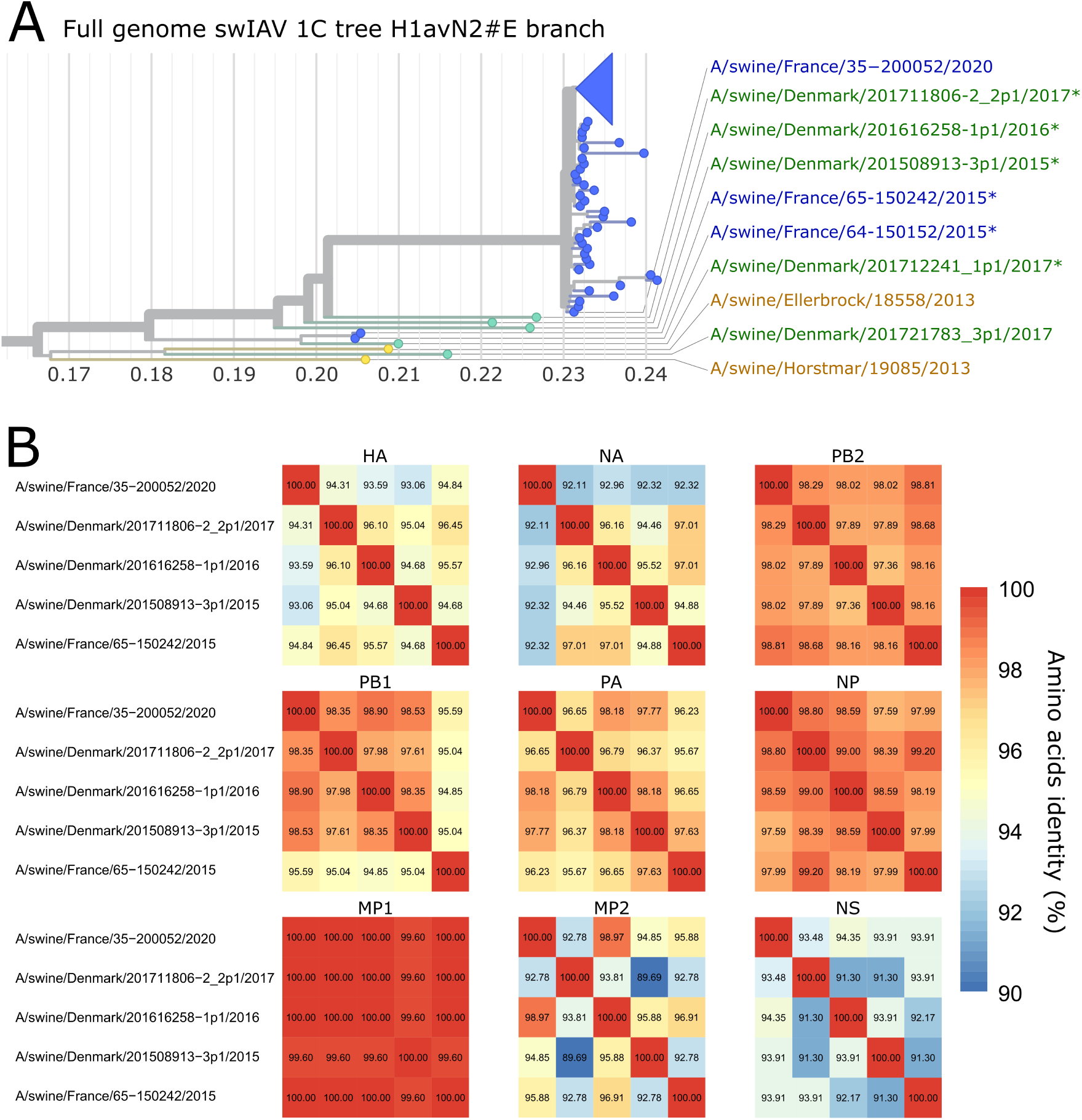
Probable origins of the H1_av_N2#E viruses detected in France from 2020 to 2022. **A.** Left: H1_av_N2#E viruses branch of the full genome maximum-likelihood divergence tree of the 1C swIAV strains in France and Europe. Right: name of the publicly available European sequences, as of July 2024, that are the most closely related to the H1_av_N2#E viruses detected in France. The A/swine/France/35-200052/2020 corresponds to the H1_av_N2#E detected in France that is the closest to pre-2020 sequences. Sequences annotated by stars represent the H1_av_N2#E publicly available sequences closest to the H1_av_N2#E collected in France during the 2020-2022 period. **B.** Amino acid identity comparison of the sequences of nine proteins of the A/swine/France/35-200052/2020 strain and the closest pre-2020 European H1_av_N2#E strains.

Further analyses of the HA segment phylogeny branch length showed that H1_av_N2#E HA diversity was quite low compared to long-established clades like H1_av_N1#A (Figure 6A). However the divergence among the H1_av_N2#E strains significantly increased on the HA gene between 2020 and 2021, as well as between 2021 and 2022, while HA divergence within H1_av_N1#A remained stable over the same period (Figure 6B). The estimation of the mutation rate was also higher for the H1_av_N2#E HA compared to the H1_av_N1#A HA, with 4.18e-3 and 3.97e-3 substitutions per site per year, respectively (Figure 6C).

**Figure 6:**
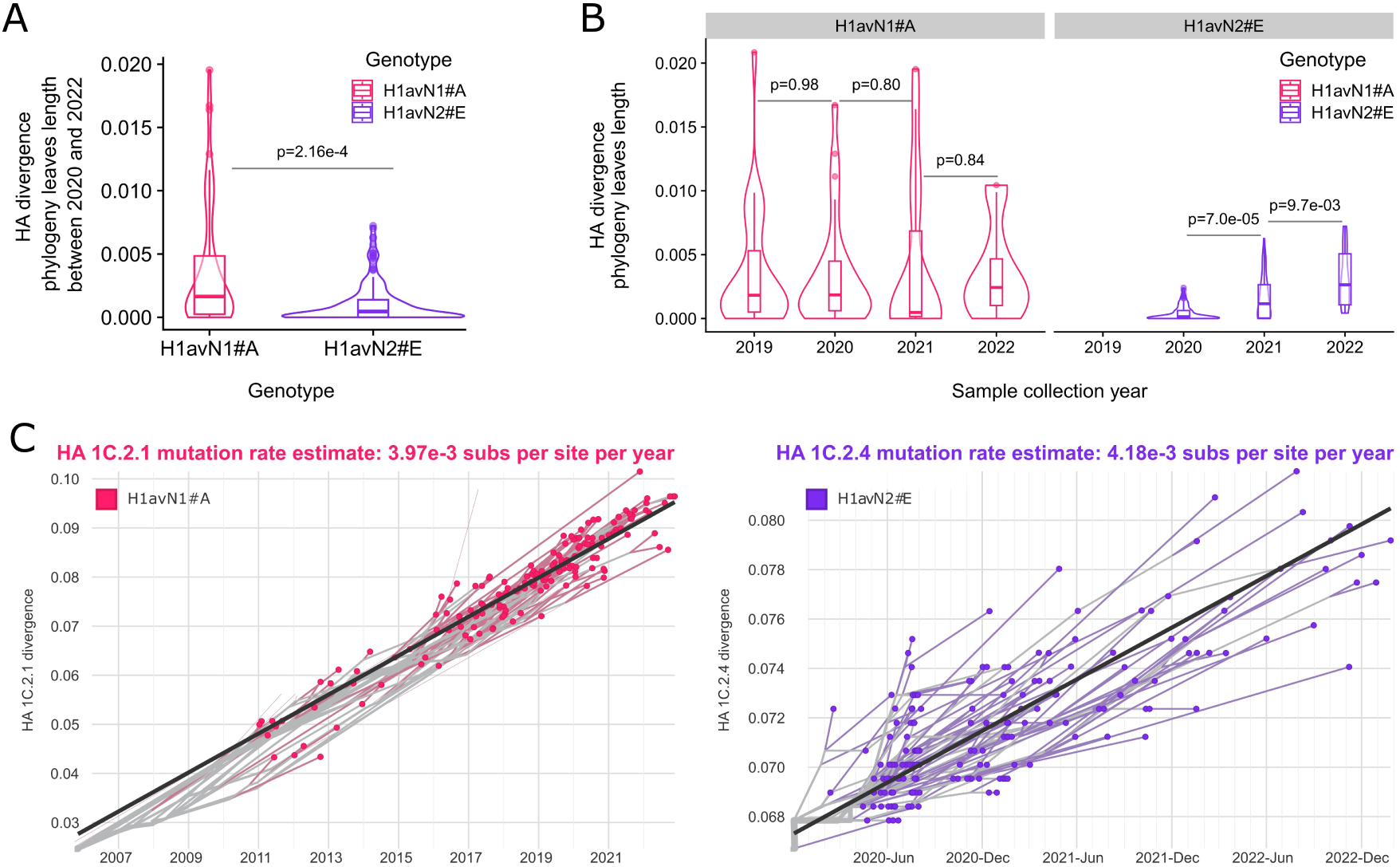
Evolution rates of the newly spreading H1_av_N2#E (purple) viruses compared to the long established H1_av_N1#A (pink) viruses in France. **A.** Distribution of the HA divergence phylogeny leaves length for H1_av_N1#A and H1_av_N2#E viruses collected between 2020 and 2022 in France, displayed as superposed boxplots and density plots. Wilcoxon Rank-sum Test p-values compare the distributions between the virus genotypes. **B.** Distribution of the HA divergence phylogeny leaves length for H1_av_N1#A and H1_av_N2#E viruses collected between 2019 and 2022 in France, split by year. Wilcoxon Rank-sum Test p-values compare the distributions within one genotype between two consecutive years. **C.** H1_av_N1#A and H1_av_N2#E France phylogenies clock view (time compared to genetic divergence) and mutation rate estimates as reported by Nextstrain. Selecting strains by their collection dates to 2019-2022 did not change the mutation rate estimate of the H1_av_N1#A viruses.

Taken together, this suggested that H1_av_N2#E strains were introduced *in toto* from abroad, possibly from Denmark, from a limited number of entry points considering the low diversity of the H1_av_N2#E at the start of the epizootic in 2020, as well as the single group structure of the H1_av_N2#E phylogeny. The evolution rates and branch length suggested that the HA-1C.2.4 of the H1_av_N2#E strains detected in France tend to diversify overtime and at a faster rate than the HA-1C.2.1 of long-established genotypes such as H1_av_N1#A.

### Antigenic characterization of H1_av_N2#E strains

In order to better understand how the H1_av_N2#E strains spread so rapidly over the country, their antigenic properties were compared to those of long-established H1_av_N1#A strains, as well as vaccine related strains (HA-1C or N2 antigens).

HA and NA antigenic sites as well as HA Receptor-Binding Site (RBS) of the H1_av_N2#E collected in France from 2020 to 2022 (n = 177, called H1_av_N2#E≥2020) were aligned (Figure 7A) with i) H1_av_N2#E strains found in Denmark and France up to 2017 (n = 6, called H1_av_N2#E≤2017, Figure 5 star-annotated sequences), ii) the HA-1C.2.2 and N2 sequences of strains included in the RespiPorc® Flu3 vaccine (Ceva Santé Animale, France), the only vaccine currently available in Europe with a 1C antigen.valence, and iii) with H1_av_N1#A strains collected during the study period (n = 84). This analysis revealed a low diversity of the antigenic sites and RBS among the H1_av_N2#E≥2020, as only 2 positions (HA 239 E or K, NA 334 N or K) displaying more than one amino acid with the alternative forms represented over 10% among the 177 analyzed sequences (Figure 7A). This suggested stable antigenic properties of the H1_av_N2#E strains collected over the study period. By contrast, compared to the other analyzed sequences, the H1_av_N2#E≥2020 strains displayed many amino acid mutations involving chemical properties change in the HA and NA antigenic sites as well as RBS (Figure 7A, orange-annotated positions). This included a deletion at position 146 in the RBS and the overlapping Sa HA antigenic-site, thus forming a 146-147 double deletion compared to the H1_av_N1#A strains and 1C.2.2 vaccine strain. Taken together, these mutations would predict differences in antigenic properties of the H1_av_N2#E strains compared to the 1C vaccine strain as well as to the previously predominant H1_av_N1#A strains.

**Figure 7:**
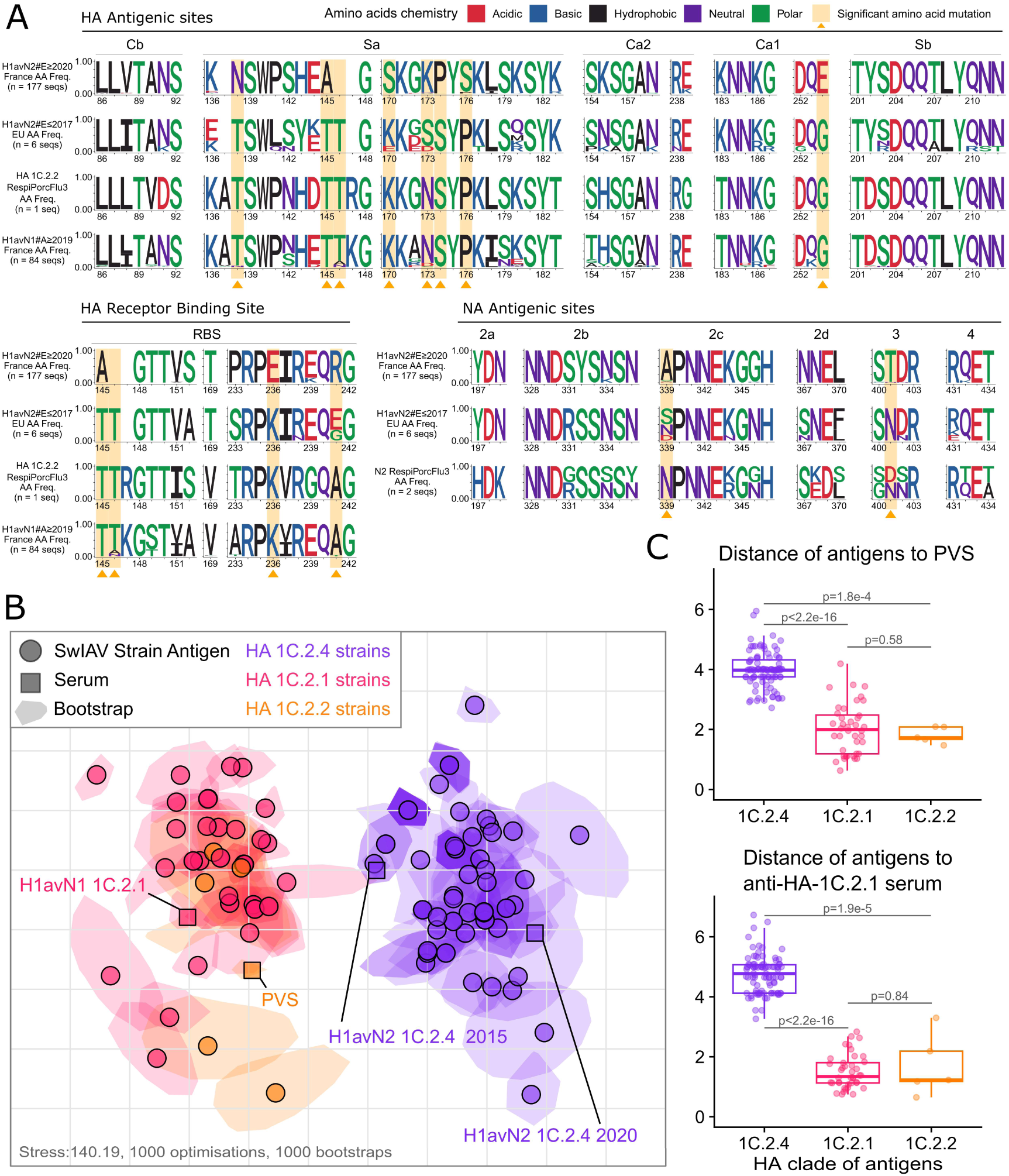
Antigenic properties of the H1avN2#E viruses detected in France between 2020 and 2022. **A.** Amino acid motifs of the HA/NA antigenic sites and HA receptor binding sites (RBS). Frequency of amino acids found for each position was calculated across the 177 HA or NA H1avN2#E sequences (top line), the 6 HA or NA H1avN2#E sequences detected in France and Denmark before 2020 (second line, stars annotated sequences in Figure 5), for the HA of the 1C.2.2 antigen contained in the RespiPorcFlu3 vaccine (third line), or for the H1avN1#A HA-1C.2.1 sequences collected in France between 2019 and 2022 (fourth line). Mutations between the top line and the other lines leading to changes in chemical properties are highlighted by an orange background and arrow. **B.** Antigenic map based on HI-test results. Virus strains detected in France from 2019 to 2022 are displayed as circles: HA-1C.2.1 (H1avN1#A) in pink, HA-1C.2.2 (H1avN1#C) in orange and HA-1C.2.4 (H1avN2#E) in purple. Four sera are displayed as squares: anti-HA-1C.2.4 towards 2015 and 2020 viruses in purple, anti-HA-1C.2.1 towards 2009 virus in pink and one post-vaccination serum (PVS, RespiPorcFlu3® trivalent vaccine) in orange. Vertical and horizontal axes are expressed in Antigenic Units (AU, Racmacs after 1000 optimisations). Variations across 1000 bootstraps are displayed as shadowed area around each dot. **C.** Antigenic distances distributions between strains and antisera of each HA clade, measured from the antigenic map and displayed as boxplots with the median as a bold line. Wilcoxon Rank-test sum p-values are represented above the compared distributions. Antigens to antigens distances distributions are represented in Supplementary Figure 5.

Antigenic distances between H1_av_N2#E strains, H1_av_N1 strains and vaccine antigens were measured in hemagglutination inhibition (HI) tests using two anti-HA-1C.2.1 hyperimmune sera, one anti-2015-HA-1C.2.4 serum, one anti-2020-HA-1C.2.4 serum and one post-vaccination serum (PVS) containing anti-HA-1C.2.2 antigen (Table 1, Supplementary Figure 4). The H1_av_N2#E 2020 reference antigen and H1_av_N2#E strains collected in 2020-2022 reacted properly with the anti-2020-HA-1C.2.4 serum (HI titers of 1280 and 584 in average, respectively), while the reaction was much lower with the anti-2015-HA-1C.2.4 serum (HI titers of 160 and 107 in average, respectively) and little to no cross-reaction was detected with the anti-HA-1C.2.1 H1_av_N1 serum (HI titers of 20 and 41 in average, respectively) and with the PVS (HI titers of 10 and 20 in average, respectively). The HA-1C.2.1 reference antigens as well as the HA-1C.2.1 and HA-1C.2.2 strains detected in the study period (H1_av_N1#A and #C, respectively) did not cross-react with the anti-2020-HA-1C.2.4 serum, and only slightly with the anti-2015-HA-1C.2.4 serum, while they reacted well with the PVS (Table 1). The differences in HI test results evidenced between H1_av_N1 and H1_av_N2#E strains were significant for all antisera (Supplementary Figure 4), which highlighted the inability of anti-HA-1C.2.1 sera and PVS to recognize H1_av_N2#E≥2020 strains antigens.

**Table 1.**
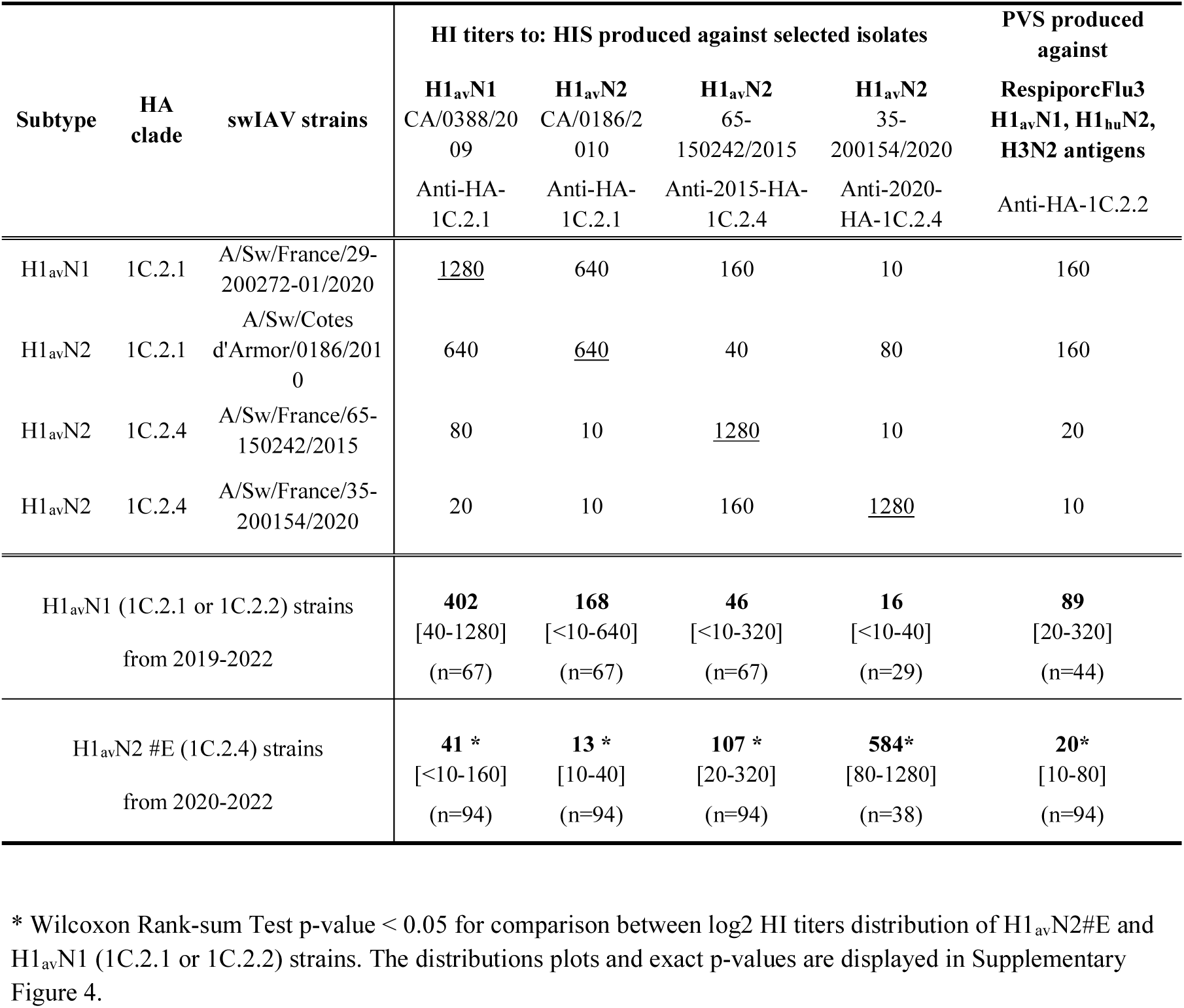
Cross-hemagglutination inhibition (HI) data obtained for H1_av_N1 and H1_av_N2#E viruses against hyperimmune sera (HIS) and post-vaccine serum (PVS) produced in SPF pigs. The table reports the HI titers for reference antigens (homologous HI titers are underlined). Then, the geometric mean HI of each group of tested strains is indicated in bold, significant difference (T-test p-value <0.05) between the two groups is pointed by an asterisk, the range of HI antibody titers obtained within each group is reported in square brackets and the number of tested strains in brackets.

An antigenic cartography was built using HI titers of HA-1C.2.4, 1C.2.1 and 1C.2.2 strains against anti-2015-HA-1C.2.4, anti-2020-HA-1C.2.4, anti-HA-1C.2.1 sera and the PVS (Figure 7B). HA-1C.2.4 antigens displayed consistent antigenic distances between them, with a median of 1.36, but a median antigenic distance of 4.15 and 4.56 units when compared to HA-1C.2.1 and HA-1C.2.2 antigens, respectively (Supplementary Figure 5). These differences were found signifcant (Wilcoxon rank-sum test p-value < 1.13e-10). The same was observed when measuring antigen to serum distances (Figure 7C): HA-1C.2.4 strains displayed significant antigens-serum distances over 4 (p-values < 2.1e-3, Wilcoxon Rank-sum Test) for both anti-HA-1C.2.1 serum and PVS when compared to HA-1C.2.1 and HA-1C.2.2 strains, which displayed non-significant median distances below 2 antigenic units (p-values > 0.20, Wilcoxon Rank-sum Test) (Figure 7C). Taken together, the newly predominant H1_av_N2#E strains displayed significant (27–29) antigenic distances with HA-1C.2.1 and HA-1C.2.2 strains, as well as with the PVS.

### Epidemiology of the H1_av_N2#E outbreaks

Owing to the detection rate of swIAV strains belonging to the emerging H1_av_N2#E genotype in 2020, we aimed to rapidly discriminate them within H1_av_ (clade 1C.2) strains following the initial HA/NA molecular subtyping step. Thus, we developed a Taqman RT-qPCR assay targeting a specific region of the HA-1C.2.4 segment of the H1_av_N2#E strains, an assay which displayed 100% analytical and diagnostic specificity and 96.55% diagnostic sensitivity (see Materials and Methods section).

This new HA-1C.2.4-gene RT-qPCR led to the identification of 352 cases of infection by an H1_av_N2#E virus from 2020 to 2022. Epidemiological data collected at the time of pig sampling in those affected herds were compared to those collected in the 272 herds that were found infected by H1_av_N1 strains (HA-1C.2.1) during two periods of time, i.e.2019-2020 and 2021-2022. When comparing the outbreaks caused by H1_av_N2#E and H1_av_N1 viruses, no difference was found concerning their repartition within the different types of farms (Table 2). During the early period of the epizootic (2019–2020), H1_av_N2#E cases were more often associated to sporadic acute infection (“classical form”, RR=1.93 [1.53-2.43], 77.4% versus 45.7%, Chisq p=3.5e-8), and to clinical signs more severe than the norm (RR=1.50 [1.15-1.95], 50% versus 30.9%, Chisq p=1.6e-3) compared to H1_av_N1 cases which displayed recurrent infection patterns with moderate clinical signs (Table 2). No differences between both groups were significant when comparing the form and the clinical intensity of the outbreaks that occurred in 2021-2022. H1_av_N2#E infections were reported in pigs of all physiological stages just like H1_av_N1 viruses. However, during the early period of epizootic, the proportion of piglets infected before 10 weeks of age was weaker in H1_av_N2#E than in H1_av_N1 outbreaks (RR= 0.66 [0.51 - 0.85], Chisq p=7.4e-4), while the proportions of growing pigs (≥10 weeks) were similar in both groups. The proportion of breeding animals infected by H1_av_N2#E was higher than by H1_av_N1 (RR=2.99 [1.58 - 5.64], 24.3% versus 5.5%, Chisq p=7.2e-6). 75.4% of H1_av_N2#E infected gilts and sows were hosted in farms that applied mass or batch-to-batch vaccination programs with the Respiporc® Flu3 trivalent vaccine. When only considering infected breeders, more H1_av_N2#E outbreaks were reported in vaccinated farms than H1_av_N1 in 2021-2022 (RR=3.53 [1.63-7.69], Chisq p=6.7e-4). However no significant differences were found in distribution of outbreaks in vaccinated farms based on the detection of infected animals of all physiological stages.

**Table 2.** Distribution of swIAV-positive cases belonging either to H1_av_N2#E (HA-1C.2.4) or to H1_av_N1 (HA-1C.2.1) subtype according to epidemiological data recorded in infected farms. ^a^ The ‘Classical’ form is defined by a sporadic acute infection and the ‘Recurrent’ form is related to an endemic pattern of infection at the herd level, affecting each successive batch of pigs at a given physiological stage (25). ^b^ The ‘Moderate’ intensity of the disease is defined by clinical signs lasting no more than 2–3 days at the individual level and without mortality while the ‘High’ intensity form lasts longer than usual and/or can be associated to some mortality. ^c^ Vaccination against swine influenza was only applied in breeding animals. Results of Chi-squared test p-value: <0.05.

|  | 2019 - 2020 |  | 2021 - 2022 |  |
| --- | --- | --- | --- | --- |
|  | H1<sub>av</sub>N1 | H1<sub>av</sub>N2 | H1<sub>av</sub>N1 | H1<sub>av</sub>N2 |
| **Type of production farm** |  |  |  |  |
| Farrowing-to/post-weaning (N, NPS) | 7 | 13 | 9 | 24 |
| Post weaning (PS) | 1 | 1 | 0 | 1 |
| Farrowing-to-finishing (NE) | 104 | 113 | 87 | 135 |
| Post weaning-finishing (PSE) | 31 | 17 | 17 | 29 |
| Finishing (E) | 8 | 7 | 2 | 7 |
| **Epidemiological pattern of infection<sup>a</sup>** |  |  |  |  |
| Classical form | 64 | 113* | 71 | 112 |
| Recurrent form | 76 | 33 | 38 | 69 |
|  | p=3.5e-8, RR=1.93 [1.53 - 2.43] |  |  |  |
| **Severity of clinical signs<sup>b</sup>** |  |  |  |  |
| Moderate | 94 | 63 | 63 | 108 |
| High | 42 | 63* | 25 | 36 |
|  | p=1.6e-3, RR=1.50 [1.15 - 1.95] |  |  |  |
| **Physiological stage of pigs** |  |  |  |  |
| piglets (<10 weeks) | 96 | 67* | 61 | 108 |
|  | p=7.4e-4, RR=0.66 [0.51 - 0.85] |  |  |  |
| growing (≥10 weeks) | 41 | 42 | 35 | 54 |
| breeding | 8 | 35* | 17 | 30 |
|  | p=7.2e-6, RR=2.99 [1.58 - 5.64] |  |  |  |
| **Vaccination program<sup>c</sup>** |  |  |  |  |
| *All physiological stages of infected animals* |  |  |  |  |
| Not vaccinated farm | 70 | 61 | 62 | 88 |
| Vaccinated farm | 76 | 85 | 48 | 107 |
| *Only considering infected breeders* |  |  |  |  |
| Not vaccinated farm | 2 | 11 | 11 | 4 |
| Vaccinated farm | 6 | 23 | 6 | 23* |
|  |  |  | p=6.7e-4, RR=3.54 [1.63-7.69] |  |

## Discussion

The surveillance of swIAVs organized in France since the 2009 pandemic and the in-depth characterization of the strains, notably thanks to the Résavip surveillance network, allowed to regularly monitor their genetic and antigenic evolution. Subtyping, sequencing, phylogenetics and the compilation of related epidemiological data helped answering questions concerning the etiology of cases of acute respiratory syndromes and have led to the development of new diagnostic tools to adapt to swIAV genetic diversification. From 2019 to 2022, when compared to the previous 2000 to 2018 period (25), new H1_av_N2 genotypes, i.e. H1_av_N2#G, #H, #I and the reassortants H1_av_N1#B and #D, have been characterized while remaining sporadic. More importantly, a new predominant H1_av_N2 genotype was identified, the H1_av_N2#E, which was detected only sporadically in 2015 in the South-West of France but reappeared in the North-West in February 2020.

In 2020, the number of swIAV herd infection cases detected through the passive surveillance has doubled compared to previous years, coinciding with the emergence of H1_av_N2#E viruses which widely spread in the country. H1_av_N2#E quickly became the most frequently detected genotype, especially in Brittany, which accounted for 56.3% of the French pig population in 2020 in only 5% of the country’s territory (30). The H1_av_N2#E genotype was not related to viruses that were previously circulating in France and was found to be more similar to the enzootic Danish H1_av_N2 viruses (16). The H1_av_N2#E genotype exhibits an HA gene classified by OFFLU within HA-clade 1C.2.4 among the 1C Eurasian avian lineage (https://www.offlu.org/wp-content/uploads/2021/03/OFFLU-VCM-SWINE-FINAL3_to_be_Uploaded.pdf), a clade shared by other H1N2 viruses found in Denmark, Spain, Italy and Bulgaria (21). Meticulous phylogenic analyses revealed that H1_av_N2#E internal protein-encoding genes could be classified as a new EA sub-clade that we named EA-DK considering the prevalence of Danish strains at the root of this clade. Three H1_av_N2 strains from Denmark were closer to the H1_av_N2#E strains when considering the entire IAV genome. Meanwhile, these Danish HA sequences shared 94.3% identity to the earliest detected H1_av_N2#E strain in 2020 in France, which was on par with the 94.8% HA identity of the 2015 H1_av_N2#E strains detected in south-western France. This led to hypothesize an introduction *in toto* from Denmark between November 2018 and January 2020, but considering the sparseness of the swIAV sampling in Europe, the virus could have transited to France from another European country. It is however important to note that France imported live pigs for rearing / breeding mainly from Denmark from 2019 to 2021 (Supplementary Figure 6).

In addition to this newly predominant H1_av_N2#E virus, four sporadic H1_av_N2 genotypes were detected during the study period. Three of them seemed to have been introduced *in toto* from foreign countries just like the H1_av_N2#E, i.e. H1_av_N2#F, H1_av_N2#G and H1_av_N2#H. The introduction from abroad of all these genotypes recalled the importance of testing animals for swIAV positivity before exportation to another country, as well as quarantining animals in specific rooms with adapted biosecurity measures when entering in intensive confined farms, regardless of their origin. Such adequate biosecurity measures should be adopted and applied at the European level to control swIAV spreading as efficiently as possible. Moreover, the circulation of additional viral genotypes in herds increases the risks of simultaneous infections by different influenza A viruses, which can themselves lead to the emergence of new reassortant viruses (31). This threat was here illustrated by the detection of three new reassortant genotypes, i.e. H1_av_N1#B, H1_av_N1#D, and H1_av_N2#I, which likely formed on the territory from genotypes introduced from abroad. The impact of such reassortants on the evolution of swIAV strains in France cannot be predicted and this poses new challenges and threats considering their zoonotic potential and ability to infect other animal species.

Since the emergence of the H1_hu_N2 virus in 1994, which contributed to the disappearance of the H3N2 virus in North-Western France (22, 32, 33), the H1_av_N2#E virus is the first to drastically disrupt the relative proportions of swIAV lineages that circulated in France for 30 years. Such a diversity switch in the swIAV population at a national scale was unprecedented, and the collected sequencing data constitute a unique opportunity to follow the evolution of a newly spreading swIAV. Interestingly, H1_av_N2#E displayed low diversity at the start of the epizootic in 2020 on their HA gene, suggesting a restricted source of introduction in France, and as time passed, their diversity increased. The becoming of such a virus remains unknown and only future sequences analyses will allow us to understand this virus evolution pattern. Comparatively, in 2009, despite the pressure of the H1N1pdm virus causing a pandemic in humans and a panzootic disease in pigs worldwide, French pig farms were weakly affected. The H1N1pdm virus failed to prevail against the enzootic H1_av_N1 and H1_hu_N2 viruses in north-western regions where they were well established (34). This context arose the question of the factors allowing a new swIAV genotype to emerge, become epizootic, and latter, enzootic.

Several biotic and abiotic factors drive the becoming of an airborne virus, like influenza, to become epizootic after its emergence: i) the density of sensitive host, i.e. the proportion of host population without preexisting immunity (35, 36) and ii) the fitness/virulence of the virus which defines its ability to infect the host cells or prevail in the case of viral competition (37).

Concerning the proportion of a host population without preexisting immunity, we showed in the present study that H1_av_N2#E strains were significantly distant antigenically (above 4 AU) from HA-1C.2.1 strains, as well as from PVS Respiporc® Flu 3 antigens. These results are consistent with those we reported in a previous study where H1_av_N2#E was efficiently neutralized by homologous anti-serum, but not by anti-HA-1C.2.1 serum or PVS (38). In this previous study, we also reported that RespiporcFlu3 vaccinated piglets were not fully protected from H1_av_N2#E infection whereas they were against H1_av_N1#A under our experimental conditions (38). Contrastingly, H1N1pdm strains were shown to cross-react with anti-HA-1C.2.1 sera, and the antigenic distance between antigens was in that case below 3 (29, 39). Overall, this suggested that H1_av_N2#E viruses likely escaped the swine population pre-existing immunity, either to the previously predominant H1_av_N1#A swIAV or to the Respiporc® Flu 3 vaccine and related strains, thus contributing to the rapid spread of the H1_av_N2#E virus over the country, unlike H1N1pdm virus in 2009. The introduction of the H1_av_N2#E virus in a high density pig farming region such as Brittany also most likely contributed to the successful H1_av_N2#E spread. This raises apprehension considering the intensification of the pig farming, with a stable decrease of the number of holdings, but an increase or stabilization of the number of pigs since 2010 (30).

Concerning the virus fitness and ability to infect new hosts, this is often associated to the induced symptoms and viral shedding. The beginning of the H1_av_N2#E epizootic was associated with a higher proportion of high severity clinical signs in HA-1C.2.4 swIAV infected animals compared to H1_av_N1 infected animals. This is in accordance with our experimental results as we also showed that H1_av_N2#E infected animals displayed stronger and prolonged clinical signs compared to H1_av_N1 infected ones (38). The H1_av_N2#E strains thus appeared to be more virulent than H1_av_N1 strains, at least before significant genetic evolution, which most likely contributed to their rapid spread and to the increase in the total number of swIAV outbreaks investigated in 2020 through the passive surveillance programs. Interestingly, the virulence of the H1_av_N2#E outbreaks during the 2021-2022 period was not significantly different from that of the H1_av_N1 outbreaks. This suggests that a certain level of population immunity may have developed during the H1_av_N2#E enzootic period.

To conclude, H1_av_N2#E was detected as the new predominant swIAV genotype in France and while some elements concerning their antigenic and virulence properties were studied, their airborne transmissibility and resistance in the environment to temperature or pH remain to be characterized and compared to the previously predominant H1_av_N1 viruses (40). The pathway of introduction H1_av_N2#E swIAVs to France from abroad as well as the spreading dynamics in France also remain to be elucidated. Phylodynamic and phylogeographic analyses at the scale of France, and eventually Europe, linking precise swIAV samples geographic and sequence data with animal movements and epidemiological data would help to characterize swIAV transmission pathways. Moreover, H1_av_N2#E spread in France raises the question of potential spillovers of the genotype to other countries importing live pigs from France. To prevent new epizootics and the establishment of new enzootic viruses, it is crucial to prevent the introduction of new swIAV genotypes in herds by strengthening quarantine measures and encouraging the monitoring of swIAV infections in live swine traded, especially for breeding, both between and within European countries, as is done for other diseases (41), even though swine influenza is not a regulated disease.

## Material and Methods

### Samples, swIAV detection and HA/NA molecular subtyping

Nasal swabs (MW950Sent2mL Virocult®, Kitvia, Labarthe-Inard, France) were collected from January 2019 to December 2022 from pigs with acute respiratory disease in France thanks to the passive surveillance program implemented by Résavip (https://www.plateforme-esa.fr/fr/virus-influenza-porcins) (26), diagnostic offers made by Ceva Santé Animale (Libourne, France) or requested by veterinarians, or epidemiological investigations conducted by the French Agency for Food, Environmental and Occupational Health & Safety (ANSES, Ploufragan, France). The Résavip network alone covered 207 different farms in 2019, 272 in 2020, 238 in 2021 and 200 in 2022, spanning 9 out of the 13 administrative regions of metropolitan France. In parallel with the sampling, the veterinarians collected outbreak-related information about the sampling date, the herd type, the vaccination program, the age and/or physiological stage of the sampled animals, the influenza-like illness intensity, and the epidemiological pattern of the outbreak.

Nasal swab supernatants were first screened in local veterinary labs for the presence of swIAV genome using M gene RT-qPCR from two commercial kits validated by the French National veterinary Reference Laboratory (NRL) for Swine Influenza (ANSES, Ploufragan, France) (42). Then, swIAV were subtyped by the NRL or LABOCEA22 (Ploufragan, France) using RT-qPCR assays specific to HA and NA genes of swIAV lineages known to circulate in France in 2018 (H1_av_, H1_hu_, H1_pdm_, H3, N1, N1_pdm_ and N2), as previously described (43). A new RT-qPCR assay was developed to specifically target HA-1C.2.4 of H1_av_N2#E strains (see at the end of the Material and Methods).

### swIAV sequencing

Viral RNA was extracted using the NucleoSpin® RNA Kit (Macherey-Nagel, Düren, Germany) directly from nasal swab supernatants, or after virus propagation on Madin-Darby Canine Kidney (MDCK) cells according to standard procedures (25). All genomic segments of swIAV were amplified simultaneously before sequencing with the SuperScript III one-step RT-PCR system (Thermo Fisher Scientific, Waltham, MA, USA) using the universal IAV primers (44). The nucleotide sequences were obtained by next-generation sequencing (NGS) on Thermo Fisher Scientific’s Ion Proton instrument (Thermo Fisher, Carlsbad, CA, USA). The reads were cleaned with Trimmomatic v0.36, assembled into contigs using Spades, which were subsequently compared to an NCBI IAV database using megablast to find reference sequences for each segment, on which the sequenced reads where mapped using BWA v0.7.15-r1140. The resulting BAM files were used to call consensus sequences using Influenza Sequences Toolbox v0.4 callConsensus tool (45), which is notably based on seqkit (46) and samtools consensus (47) with the following parameters “-t :hiseq --het-scale 0.42 --low-MQ 9 --scale-MQ 1.15”. The NGS raw data obtained in this study have been made available in NCBI Bioproject <u>PRJNA623701</u>. Only consensus sequences of complete genome were deposited in Genbank and identification numbers are reported in Supplementary Table 2. If the consensus sequence did not cover the entire coding sequence, the segment was considered as partially sequenced and the strain genome as incomplete. Sequences from incomplete genomes were used for genotyping when they presented enough informative loci and are available upon request.

### swIAV genotyping

In addition to molecular sub-typing, the genetic lineage of each genomic segment was determined by using a BLAST strategy implemented in the Influenza Sequences Toolbox v0.4 genotypeFasta tool (45). This tool used BLAST to compare the obtained strains sequences to an in-house hand-curated database which is continuously incremented with data from swIAV strains identified in France since 2000, which is available upon specific request. For each gene, the best BLAST hit genotyping result was compared to the following 7 best hits to verify the best hit genotype consistency. The full genome genotype and final FASTA files were then called for each strain using Influenza Sequences Toolbox v0.4 annotateFasta tool, based on an in-house curated genotypes database and samples metadata database. The attributed HA genotypes were further confirmed by using the BV-BRC webserver HA subspecies classification tool using the Orthomyxoviridae – Swine Influenza H1 global classification (https://www.bv-brc.org/app/SubspeciesClassification). The consistency of the attributed genotypes was assessed for all segments using maximum-likelihood phylogenies (see below).

### Phylogenetic analyses

All publicly available swIAV sequences were collected from GISAID and NCBI and name duplicates were removed. All sequences were genotyped using the aforementioned genotyping pipeline and only the strains displaying an HA from the 1C clade were kept for further analyses. Alignments and phylogenies were performed to determine the European strains that were closely related to the strains collected from France since 1990 by removing the poor quality sequences (displaying consecutive stretches of more than 10N) and using MAFFT (48), IQ-Tree (49) and FigTree (https://github.com/rambaut/figtree/releases). European strains located in the phylogeny in branches that displayed French strains were kept for further analyses. Phylogenies were performed using Nextstrain CLI v8.0.1 with default parameters by combining the filtered publicly available 1C European strains, the publicly available French 1C strains, as well as all 1C 2019-2022 French strains, for which a metadata table was built to store the following information: strains collection date, full genome genotype, per gene clade, country and location in France. In more details, the sequences were filtered compared to the sequences metadata table using augur filter. Sequences were then aligned using augur align and poor quality sequences were then manually inspected and removed. Maximum-likelihood divergence trees were built using augur tree, based on IQ-Tree ultrafast bootstraps, with the following parameters ’--override-default-args --substitution-model GTR+G4+F+I -- tree-builder-args="-bb 1000 -nt 8 -redo"’. The substitution model was determined as being adapted to the analyzed datasets by using the IQ-Tree ModelFinder. Time-based phylogenies were subsequently built using augur refine, based on the metadata table and resulting divergence trees with the ’--timetree --date-confidence --max-iter 30’ option, based on the TreeTime algorithm (50). The joint N1 N2 phylogenies root node date has been manually edited to reach the oldest N1 or N2 common ancestor date for visualization purposes of time-resolved trees. The resulting phylogenies and associated metadata were finally stored in a json file using augur export v2. The resulting auspice Nextstrain instances were then made available online: https://nextstrain.org/community/gtrichard/swIAV-H1av-France-2019-2022/. The final phylogeny plots were made by extracting vector graphics from the Nextstrain instances and by further modifying them using Inkscape.

### swIAV genotypes distribution over time and space

The distribution of the genotyped viruses over the regions of France from 2019 to 2022 displayed as pie charts over the France map was generated using the sf, ggplot2 and dplyr R packages as well as the France regions GeoJSON (https://france-geojson.gregoiredavid.fr/repo/regions.geojson). Time-resolved metadata frequency plots representing genotypes proportions over time, were generated by extracting exact dates of sampling (already known or inferred by the phylogeny) from the time-resolved HA phylogeny by using custom a jq command-line. The distribution of the genotyped viruses over time was then assessed in two-month windows from 2019 to 2022 using streamgraph (https://github.com/hrbrmstr/streamgraph) with monotone interpolation, lubridate and dplyr R packages.

### Analysis of predicted amino acid sequences of the HA and NA encoding genes

Predicted amino acid sequences of HA and NA from studied strains were obtained using AliView with standard genetic code and were aligned to the sequences described in the relevant figures, notably in order to compare the residues present on receptor-binding site and antigenic sites previously described in literature. Amino acid frequencies and chemical properties were then represented using the ggseqlogo R package (51).

### Antigenic characterization

Swine antisera against swIAV strains representative of European enzootic lineages were produced in specific pathogen-free (SPF) pigs at ANSES facilities, as previously described (34). Four reference strains were used: A/swine/Cotes d’Armor/0388/09 (H1_av_N1, HA-1C.2.1), A/swine/Cotes d’Armor/00186/10 (H1_av_N2, HA-1C.2.1), A/swine/France/65-150242/2015 (H1_av_N2, HA-1C.2.4) and A/swine/France/35-200154/2020 (H1_av_N2, HA-1C.2.4). In addition, an antiserum against vaccine strains was produced by collecting blood of a vaccinated SPF sow one week after farrowing (52).The gilt was primo-vaccinated 6 and 3 weeks before insemination followed by three boosters 6, 3 and 2 weeks before farrowing with a 2 mL of intramuscular injection of the inactivate trivalent vaccine Respiporc® Flu3 (IDT BIOLOGIKA GmBH, Dessau-Rosslau, Germany) comprising the following strains: A/swine/Haseluenne/IDT2617/2003 (H1_av_N1, HA clade -1C.2.2, N1-EA, IG-EA), A/swine/Bakum/IDT1769/2003 (H3N2, N2-Gent/84, IG-EA) and A/swine/Bakum/1832/2000 (H1_hu_N2, HA-1B.2.1, N2-Scot/94, IG-EA).

Hemagglutination inhibition (HI) assays were performed according to standard procedures (53). Briefly, four hemagglutinating units (HAU) of MDCK-propagated virus were incubated with two-fold dilutions (starting at dilution 1:10) of the selection of swine antisera and tested against 0.5% chicken red blood cells. HI antibody titers were expressed as the reciprocal of the highest dilution inhibiting 4 HAU of virus. An antigenic cartography was done using the Racmacs package to visualize the antigenic relationships between the swIAV strains isolated in France in 2019-2022, as well as to quantify the antigenic distances between antigens and sera. Briefly, the HI table was loaded in Rstudio and a first map was made using the make.acmap function with 1000 optimisations. Unstable antigens were then removed and a new antigenic map was built with 1000 optimisations using the dimension annealing option. Antigens and sera groups information were added using the agFill and srFill functions. The bootstrapMap function was then used with 1,000 repeats with the Bayesian method and no reoptimisation and bootstrapBlobs were computed. The final antigenic map and bootstrap variations were then represented. Distances from antigens to sera were extracted with the mapDistances function, while the antigens to antigens distance matrix was extracted from the antigenic map and converted to a 3-column table. These distribution of the distances expressed in Antigenic Units were then represented as boxplots using ggplot2.

### Design of H1_av_#E RT-qPCR assay specific to H1_av_N2#E genotype

Nucleotide sequence alignments of HA-1C.2 genes from representative strains of H1_av_ lineage revealed that H1_av_N2#E strains from 2020 fixed deletions of three and six nucleotides, giving rise to the loss of three amino acids at positions 137, 146 and 147 after the methionine, respectively, as compared to H1_av_ sequences from ancestral enzootic clade 1C.2.1, which made them unique compared to other H1_av_ sequences identified in France, including those from H1_av_N2#E strains of 2015 and from the H1_av_N2#F strains (Supplementary Figure 7). A pair of primers and a TaqMan probe have thus been designed based on these HA-gene sequence alignment analyses : Fw tttcccaaagaactcatggc, at position 399-418 from the first nucleotide of the HA coding sequence MW048844; Rv gaataaggttttcccttgcttg (497–518) ; probe FAM-cgggaacaacagtttcatgctccaa-BHQ1 (431–455).

The RT-qPCR assays were run with MX3005P instrument (Stratagene, Agilent Technologies, La Jolla, CA, USA). Each PCR mixture contained 5 μL of RNA extract and 20 μL of master mix composed by 2X GoTaq Probe qPCR Master Mix, 1X GoScript RT Mix for 1-step RT-qPCR (Promega, Fitchburg, WI, USA) as well as 800 nM of each primer and 300 nM of probe. The cycling conditions used were 15 min of reverse transcription at 45°C, 2 min of denaturation at 95°C and 42 cycles of 15 sec at 95°C and 1 min at 58°C.

Several panels incorporating virus strains and/or clinical samples were constituted to evaluate the performance of the developed RT-qPCR (Supplementary Table 1). Analyses showed 100% analytical specificity. Evaluation of analytical sensitivity indicated that the assay would be able to detect equivalent to around 35 Cq-values based on in-house M gene RT-qPCR. Analyses of viral RNA from nasal swab supernatants previously identified by HA/NA RT-qPCR and sequencing showed 100% [CI95: 87.66-100 %] of diagnostic specificity, and 96.55% [CI95: 82.24-99.91 %] of diagnostic sensitivity with only one false negative sample, probably due to the low amount of genome target.

### Statistical Analyses

Statistics were performed with R 3.4.0. For epidemiological data, the difference of proportion between groups was tested using the following method: based on H1_av_N1 and H1_av_N2 positive samples, considering one epidemiological category (for instance physiological stage of pigs), H1_av_N2#E infected animals numbers were compared to H1_av_N1 infected animals numbers for a given epidemiological category (breeding versus the rest) by assembling a 2×2 contingency table and by using the epitools R package riskratio function (54). This returned for each comparison the chi square test p-value as well as the risk ratio estimate and 95% confidence intervals upper and lower bounds of one epidemiological category for H1_av_N2 versus H1_av_N1 infected animals. For antigenic data comparison, the means of HI antibody titers obtained by groups of strains were log2-transformed and compared with a Wilcoxon Rank-sum Test using R 3.4.0 (wilcox.test).

## Acknowledgments

The authors are indebted to all farmers and veterinarians for their participation to influenza virus monitoring. They gratefully acknowledge all members of Résavip, the French national public-private network for surveillance of influenza A viruses in pigs, especially regional coordinators as well as national partners involved in the dedicated swine influenza virus working group from the national platform for animal health epidemiosurveillance (https://www.plateforme-esa.fr/page/thematique-virus-influenza-chez-le-porc). They also thank Agnès Jardin et al., Ceva Santé Animale (Libourne, France) for their contribution to pig sampling and first intention analyses. Thanks also go to Nicolas Rose et al., from the Swine Epidemiological and Welfare Unit, ANSES (Ploufragan, France) for visiting pig herds and contribution to swIAV surveillance, as well as to Frédéric Paboeuf and colleagues from the SPF Pig Production and Experimentation Unit, ANSES (Ploufragan, France) for antiserum productions. Gautier Richard, Séverine Hervé, Stéphane Gorin, Stéphane Quéguiner, Nicolas Barbier and Gaëlle Simon are members of the French research network on influenza viruses (ResaFlu; GDR2073) financed by the CNRS. Gautier Richard, Séverine Hervé and Gaëlle Simon are members of the European Swine Influenza Network (ESFLU; COST action CA 21132).

## Data availability

The NGS raw data obtained in this study have been made available in NCBI Bioproject <u>PRJNA623701</u>. Only consensus sequences of complete genome were deposited in Genbank and identification numbers are reported in Supplementary Table 2.

## Author contributions

Conceptualization, GS, GR, SH, AC; investigation, SQ, NB, SG, EH, VB; formal analysis, GR, SH, AC; data curation and methods, GR, SH; visualization, GR; writing - original draft preparation, GR, SH, AC, GS; writing - review and editing, GR, SH, AC, YB, GS; project administration, GS, SH. All authors have read and agreed to the published version of the manuscript.

## Supplementary Information

**Supplementary Table 1.** Analytical specificity and sensitivity results and diagnostic ability of the developed HA-gene RT-PCR for discrimination of H1_av_N2#E genotype French strains.

**Supplementary Table 2.** Genbank accession numbers for swIAV sequences isolated in France and used in this study

**Supplementary Figure 1:**
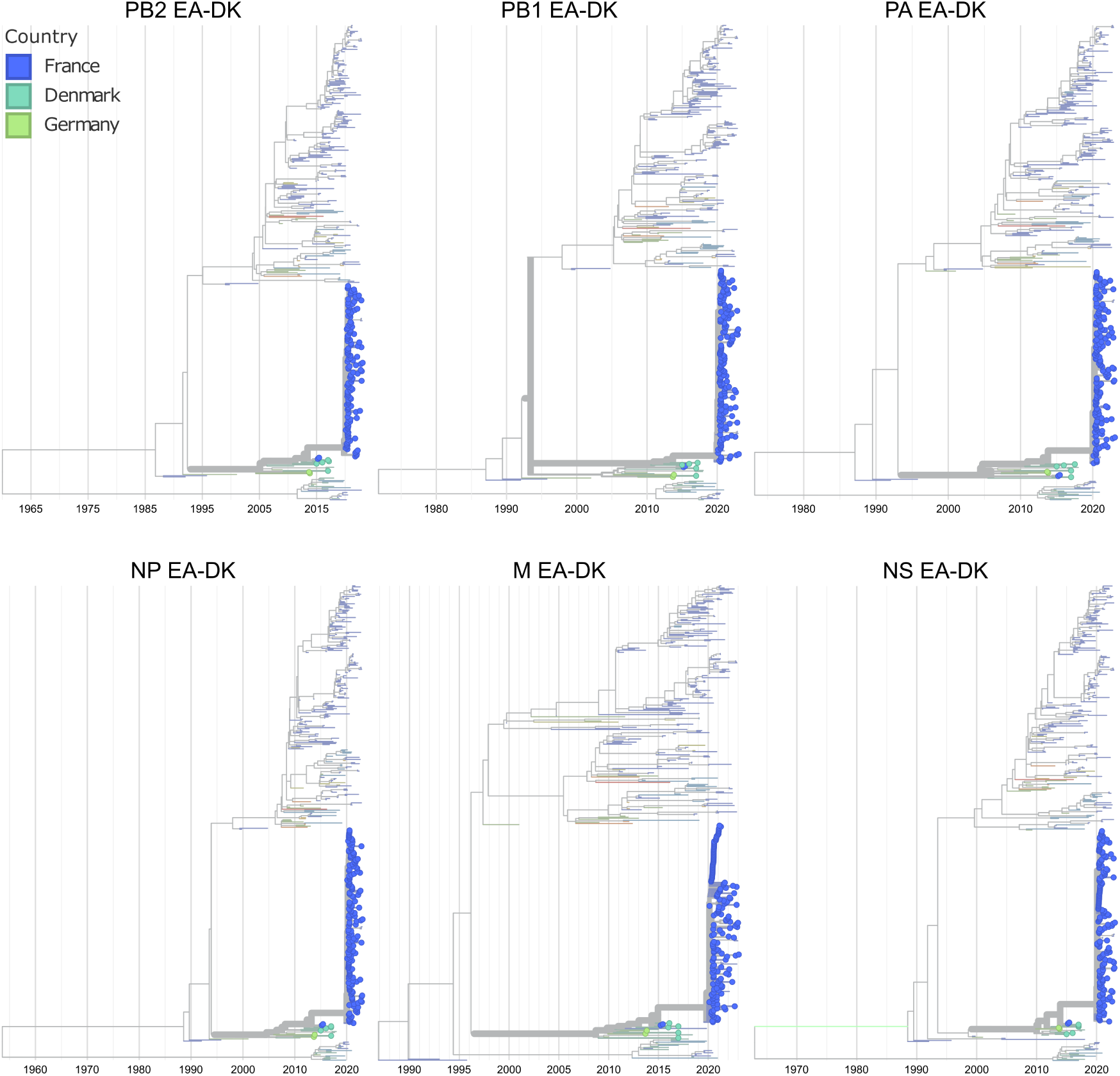
swIAV HA-1C strains EA-DK internal genes maximum-likelihood phylogenies, colored by the country of origin of the strains (France in blue, Denmark in teal and Germany in green). EA-DK strains sequences are represented by dots on the leaves and thick branches. The EA and pdm09 internal genes phylogenies of HA-1C swIAV strains are displayed as dot-less thin branches. Interactive versions of the trees can be found at https://nextstrain.org/community/gtrichard/swIAV-H1av-France-2019-2022

**Supplementary Figure 2:**
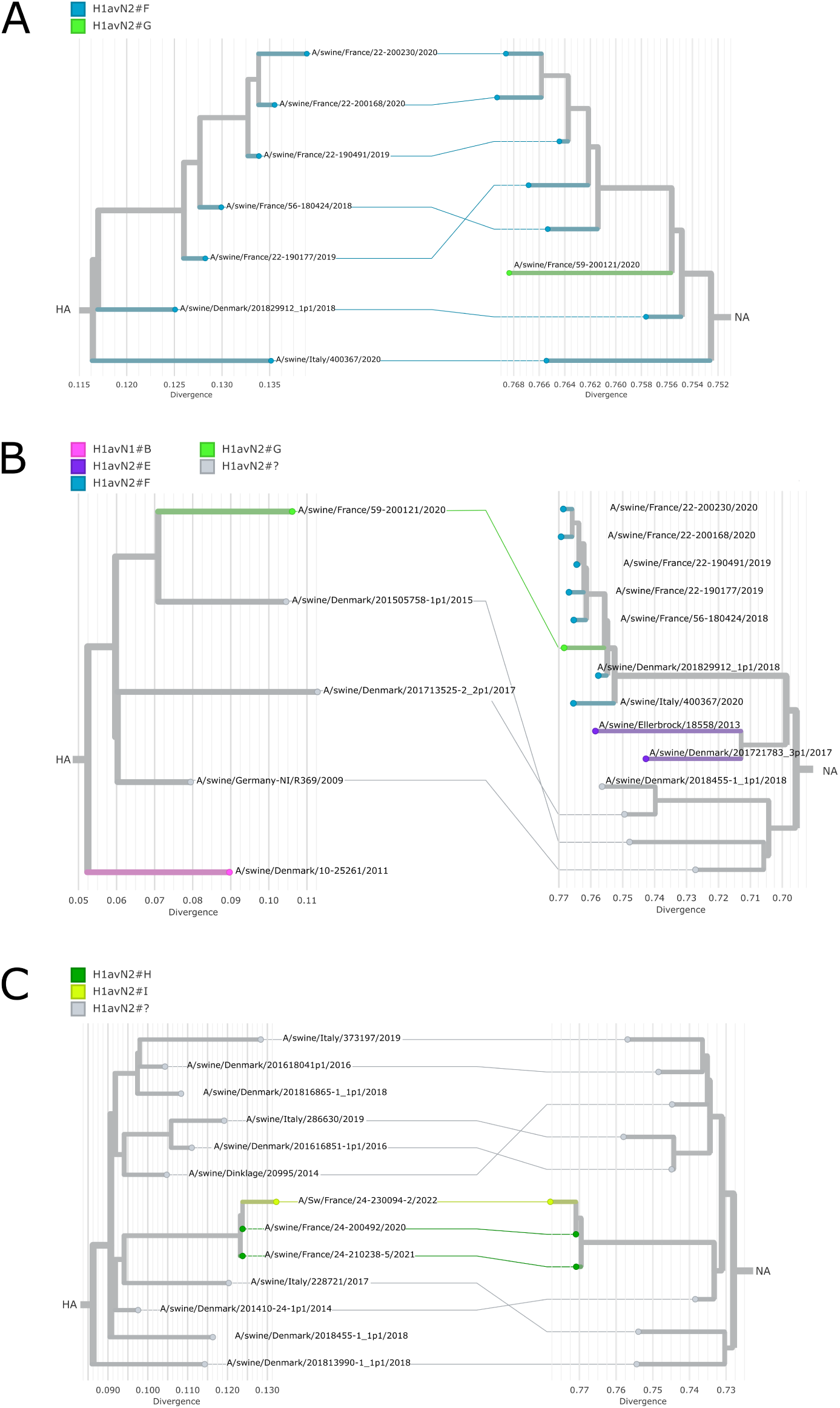
HA/NA maximum-likelihood divergence tanglegrams of H1_av_N2 swIAV genotypes found sporadically in France between 2019 and 2022, and their related publicly available European strains sequences. Left: HA sequences phylogenies. Right: NA sequences phylogenies. Middle: links between HA/NA segments each strain. **A.** H1_av_N2#F genotype tanglegram. **B.** H1_av_N2#G genotype tanglegram. **C.** H1_av_N2#H and H1_av_N2#I genotypes tanglegram. European strains with a genotype not reported in France during the 2019-2022 period or with missing segments are indicated as H1_av_N2#?. The country of origin of the sequences is indicated in the sequences names.

**Supplementary Figure 3:**
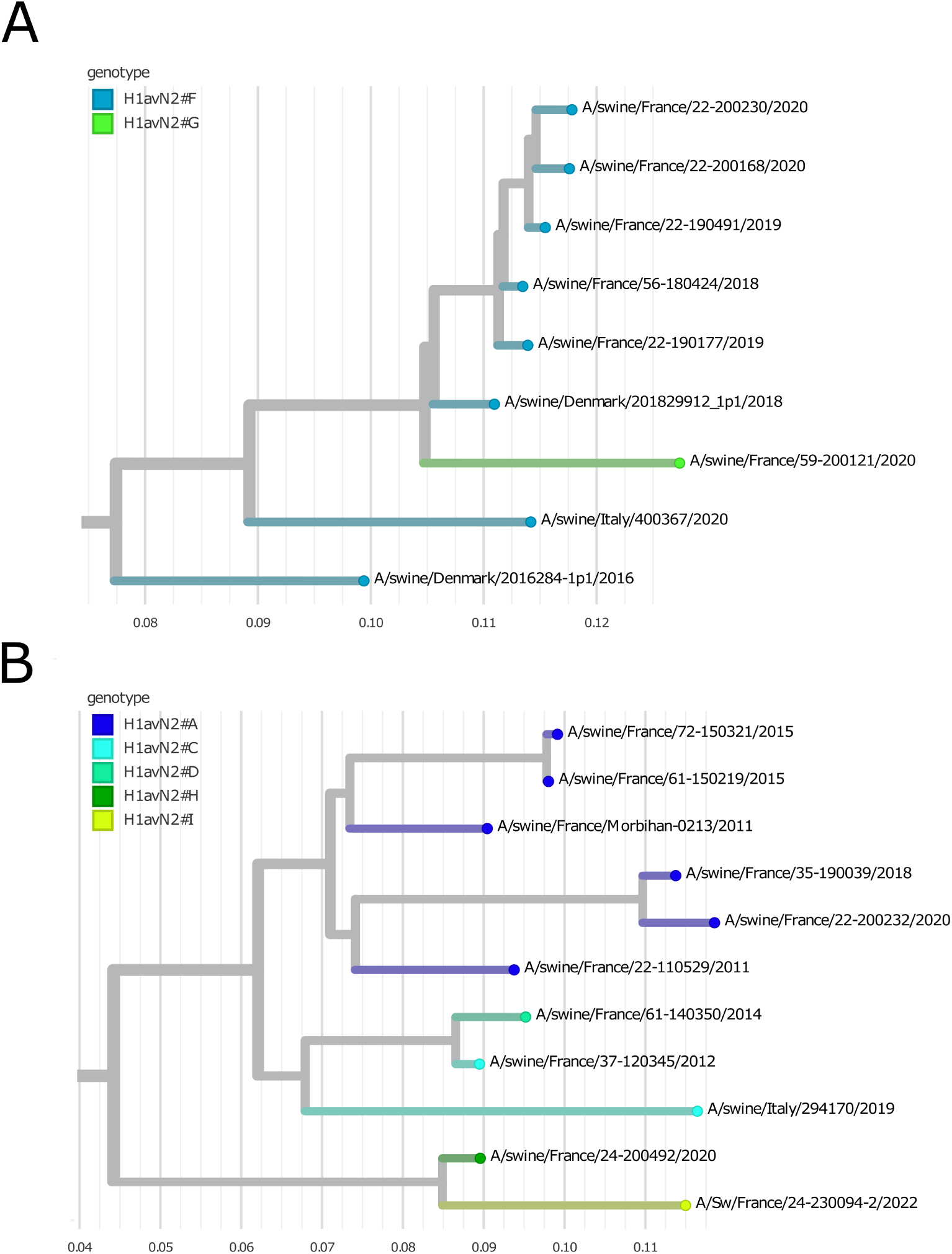
Full genome maximum-likelihood divergence phylogenies of H1_av_N2 swIAV genotypes found sporadically in France between 2019 and 2022, and their related publicly available European strains sequences. **A.** H1_av_N2#F and H1_av_N2#G genotypes phylogeny. **B.** H1_av_N2#H and H1_av_N2#I genotypes phylogeny. The country of origin of the sequences is indicated in the sequences names.

**Supplementary Figure 4:**
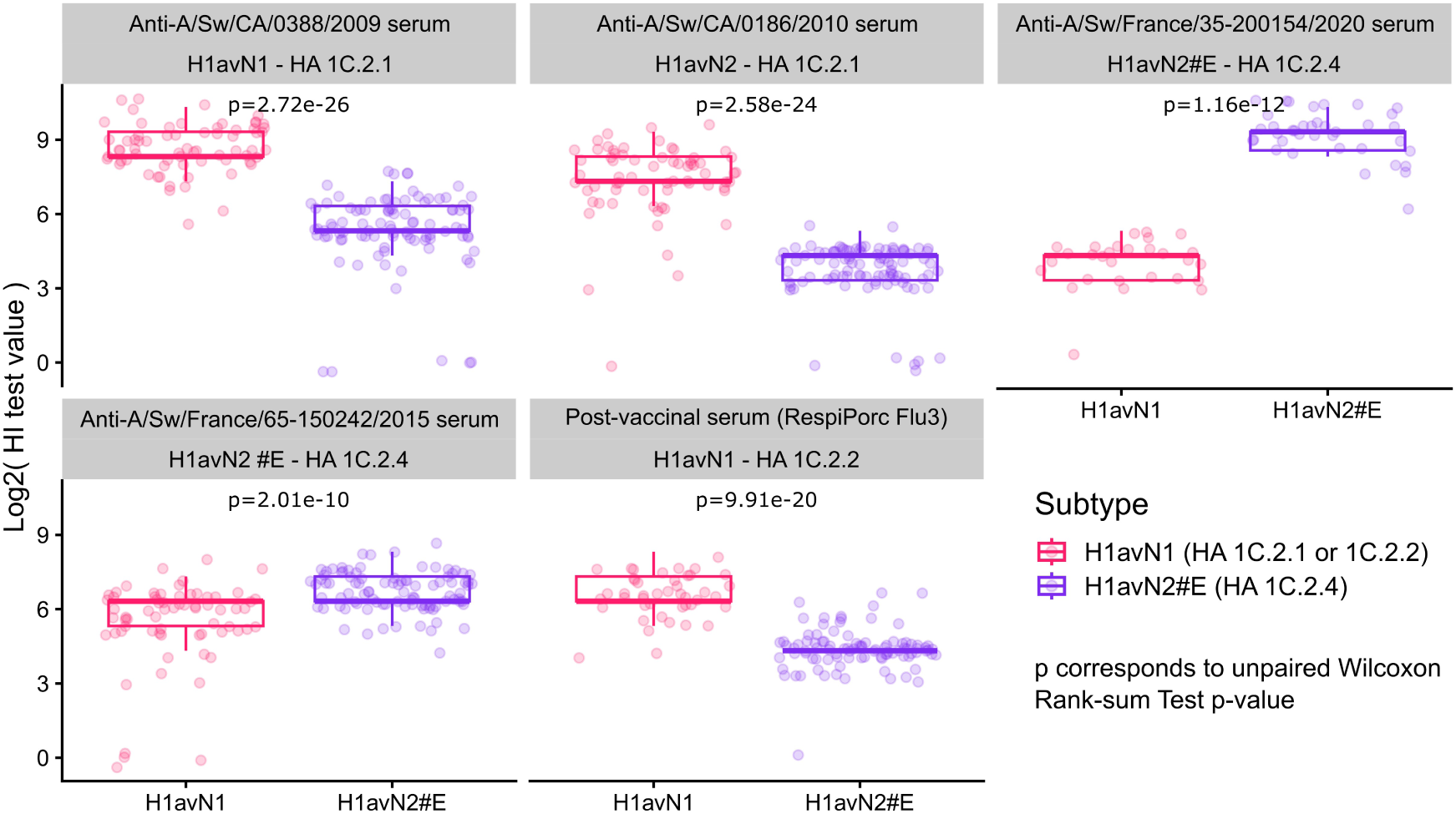
Log2 hemagglutinin inhibition (HI) test values distribution of H1_av_N1 (HA-1C.2.1 or 1C.2.2) and H1_av_N2#E (HA-1C.2.4) antigens for 5 sera whose targeted strain name, subtype and HA clade are written in gray boxes. The distributions are represented as boxplots with the median represented as a bold line, and each individual log2 HI test value is displayed as single transparent dots. Wilcoxon rank-sum tests were used to compare the H1_av_N1 and H1_av_N2#E antigens HI test values distributions for each serum.

**Supplementary Figure 5:**
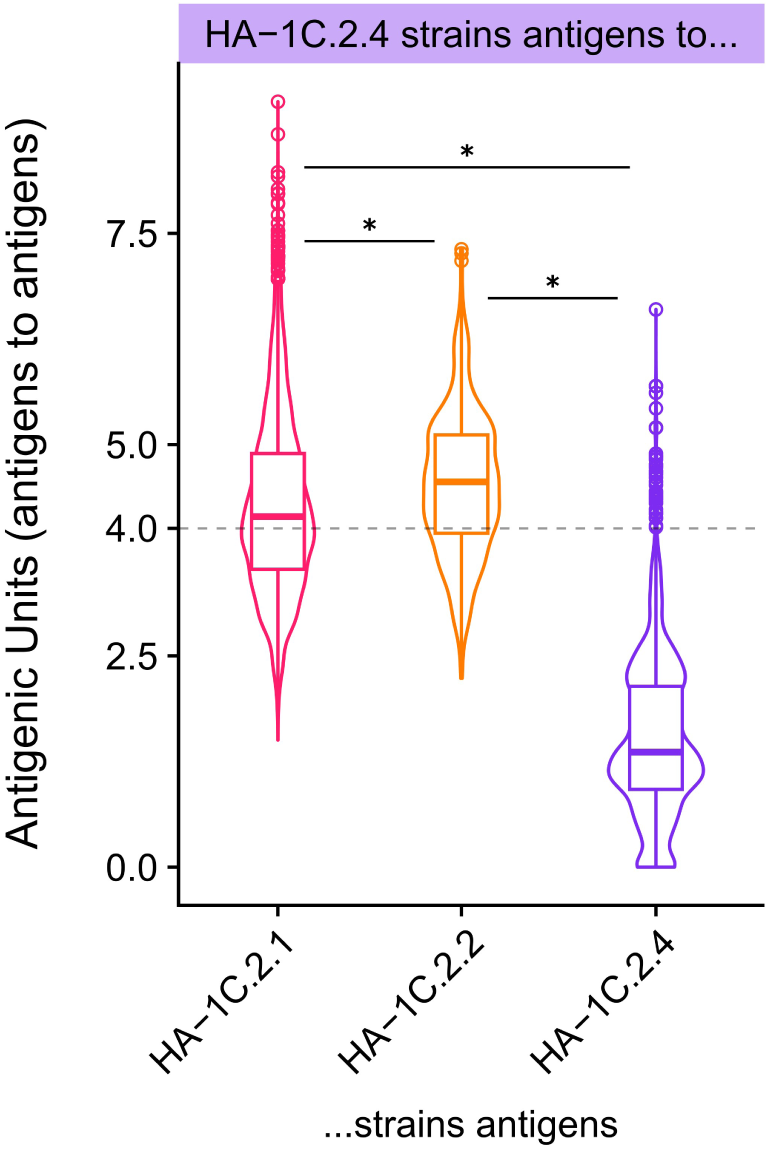
Antigenic Units distance from antigens of one HA clade to antigens of other HA clades. Each panel corresponds to antigens of one HA clade, indicated in the colored box, and their antigenic unit distances distribution to antigens of HA-clades 1C.2.1 (pink), 1C.2.2 (orange) and 1C.2.4 (purple) are represented as boxplots, with the median displayed as a bold bar. Horizontal black bars and stars represent significant distribution differences according to Wilcoxon rank-sum tests with a threshold set at p = 0.05.

**Supplementary Figure 6:**
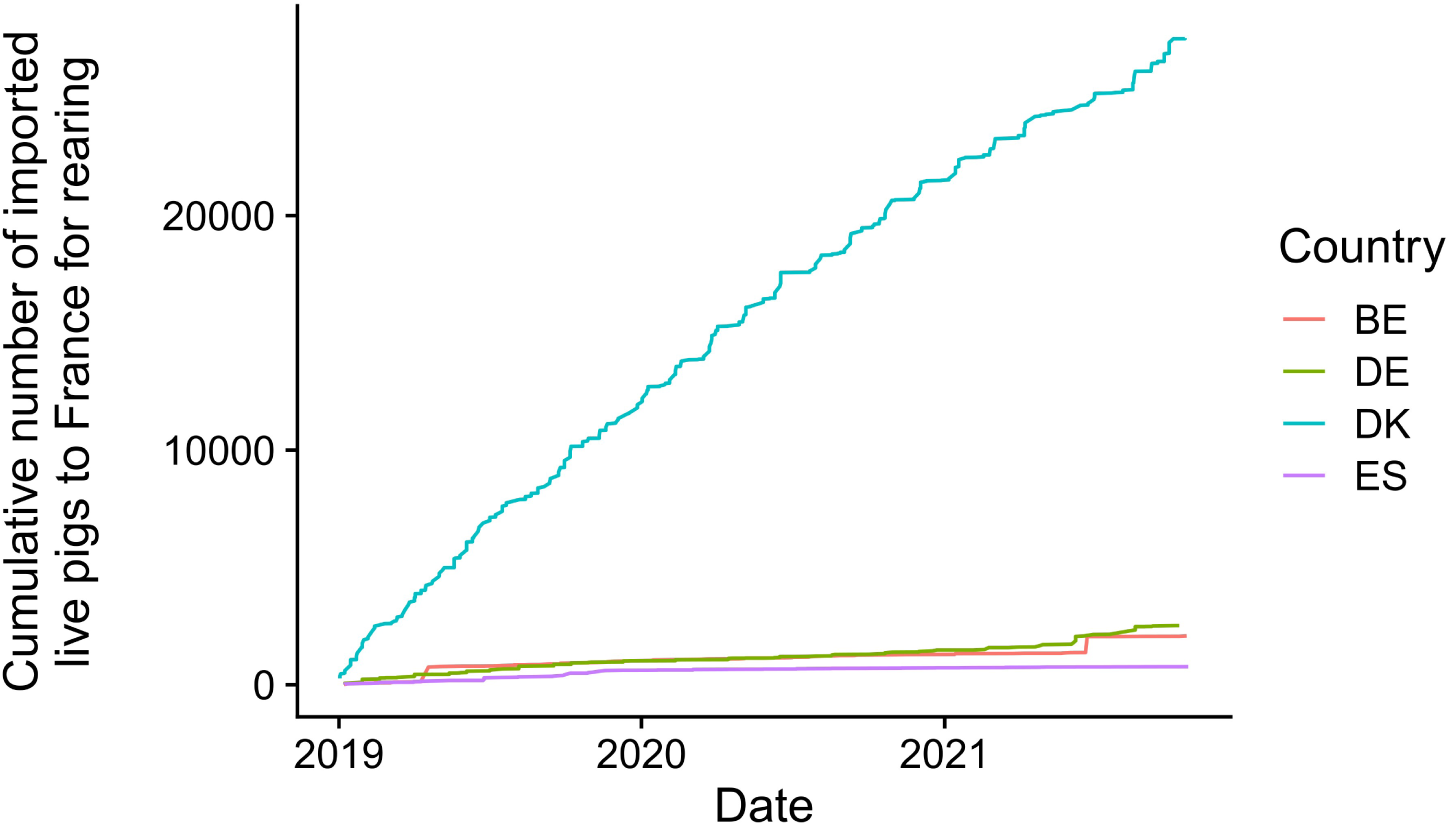
Cumulative number of live pigs imported to France for pig rearing and breeding from Belgium (BE), Germany (DE), Denmark (DK), Spain (ES). Data were collected from the TRACES database (https://food.ec.europa.eu/horizontal-topics/traces_en). Only the four major countries are displayed.

**Supplementary Figure 7:**
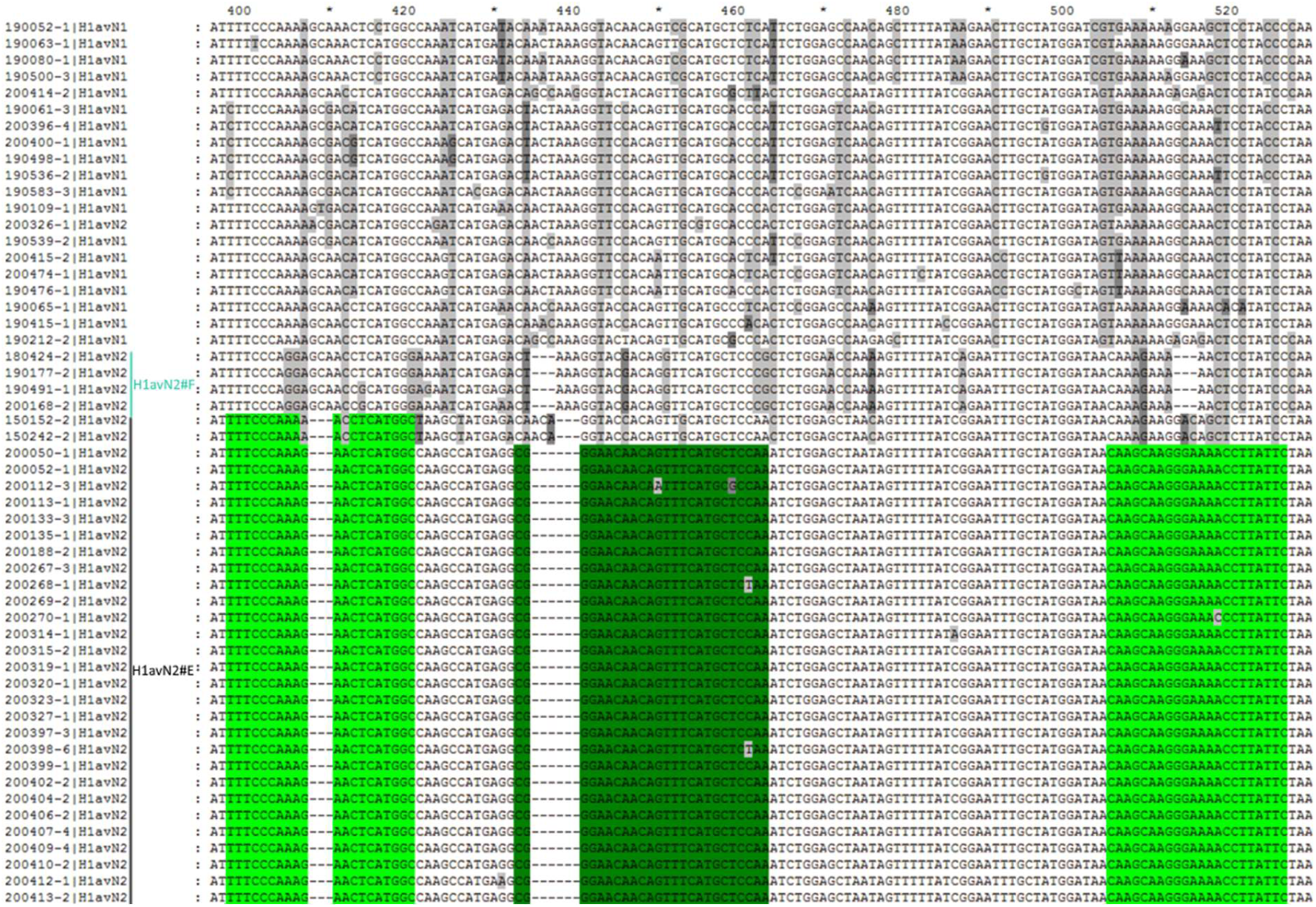
Mutations in nucleotide sequences of H1_av_N2#E strains compared to HA-1C.2.1 strains and H1_av_N2#F strains.

## References

1. Vincent A, Awada L, Brown I, Chen H, Claes F, Dauphin G, et al. Review of influenza A virus in swine worldwide: a call for increased surveillance and research. Zoonoses and public health. 2014;61(1):4–17.

2. Ma W. Swine influenza virus: Current status and challenge. Virus Research. 2020;288:198118.

3. Janke BH. Clinicopathological features of Swine influenza. Current topics in microbiology and immunology. 2013;370:69–83.

4. Short KR, Richard M, Verhagen JH, van Riel D, Schrauwen EJ, van den Brand JM, et al. One health, multiple challenges: The inter-species transmission of influenza A virus. One Health. 2015;1:1–13.

5. Vandoorn E, Leroux-Roels I, Leroux-Roels G, Parys A, Vincent A, Van Reeth K. Detection of H1 Swine Influenza A Virus Antibodies in Human Serum Samples by Age Group. Emerg Infect Dis. 2020;26(9):2118–28.

6. Henritzi D, Petric PP, Lewis NS, Graaf A, Pessia A, Starick E, et al. Surveillance of European Domestic Pig Populations Identifies an Emerging Reservoir of Potentially Zoonotic Swine Influenza A Viruses. Cell Host & Microbe. 2020.

7. Bonfante F, Fusaro A, Tassoni L, Patrono LV, Milani A, Maniero S, et al. Spillback transmission of European H1N1 avian-like swine influenza viruses to turkeys: A strain-dependent possibility? Veterinary microbiology. 2016;186:102–10.

8. Choi YK, Lee JH, Erickson G, Goyal SM, Joo HS, Webster RG, et al. H3N2 influenza virus transmission from swine to turkeys, United States. Emerg Infect Dis. 2004;10(12):2156–60.

9. Massin P, Briand FX, Cherbonnel M, Buffereau JP, Brow NIH, Eterradossi N, et al. Detection of pandemic H1N1 in turkey breeders in France. Proceedings VIIIth International Symposium on Turkey Diseases. 2010;27-29 May 2010, Berlin, Germany:212-21.

10. Starick E, Fereidouni SR, Lange E, Grund C, Vahlenkamp T, Beer M, et al. Analysis of influenza A viruses of subtype H1 from wild birds, turkeys and pigs in Germany reveals interspecies transmission events. Influenza and other respiratory viruses. 2011;5(4):276–84.

11. Zhang H, Li H, Wang W, Wang Y, Han G-Z, Chen H, et al. A unique feature of swine ANP32A provides susceptibility to avian influenza virus infection in pigs. PLOS Pathogens. 2020;16(2):e1008330.

12. Shinde V, Bridges CB, Uyeki TM, Shu B, Balish A, Xu X, et al. Triple-reassortant swine influenza A (H1) in humans in the United States, 2005-2009. The New England Journal of Medicine. 2009;360(25):2616–25.

13. Koçer ZA, Jones JC, Webster RG. Emergence of Influenza Viruses and Crossing the Species Barrier. Microbiology spectrum. 2013;1(2).

14. Nelson MI, Worobey M. Origins of the 1918 Pandemic: Revisiting the Swine "Mixing Vessel" Hypothesis. American journal of epidemiology. 2018;187(12):2498–502.

15. Anderson TK, Macken CA, Lewis NS, Scheuermann RH, Van Reeth K, Brown IH, et al. A Phylogeny-Based Global Nomenclature System and Automated Annotation Tool for H1 Hemagglutinin Genes from Swine Influenza A Viruses. mSphere. 2016;1(6):e00275–16.

16. Trebbien R, Bragstad K, Larsen LE, Nielsen J, Botner A, Heegaard PM, et al. Genetic and biological characterisation of an avian-like H1N2 swine influenza virus generated by reassortment of circulating avian-like H1N1 and H3N2 subtypes in Denmark. Virology journal. 2013;10:290.

17. Zell R, Groth M, Krumbholz A, Lange J, Philipps A, Dürrwald R. Displacement of the Gent/1999 human-like swine H1N2 influenza A virus lineage by novel H1N2 reassortants in Germany. Archives of Virology. 2020;165(1):55–67.

18. Sosa Portugal S, Cortey M, Tello M, Casanovas C, Mesonero-Escuredo S, Barrabés S, et al. Diversity of influenza A viruses retrieved from respiratory disease outbreaks and subclinically infected herds in Spain (2017–2019). Transboundary and Emerging Diseases. 2020;n/a(n/a).

19. Encinas P, Del Real G, Dutta J, Khan Z, van Bakel H, Del Burgo MÁ M, et al. Evolution of influenza A virus in intensive and free-range swine farms in Spain. Virus Evol. 2022;7(2):veab099.

20. Chiapponi C, Prosperi A, Moreno A, Baioni L, Faccini S, Manfredi R, et al. Genetic Variability among Swine Influenza Viruses in Italy: Data Analysis of the Period 2017-2020. Viruses. 2021;14(1).

21. Richard G, Byrne A, European Swine Influenza Network. European Swine Influenza Network Report on Swine Influenza A Viruses Evolution and Diversity in Europe from October 2022 to September 2023: Zenodo; 2024 2024/1/31.

22. Simon G, Larsen LE, Durrwald R, Foni E, Harder T, Van Reeth K, et al. European surveillance network for influenza in pigs: surveillance programs, diagnostic tools and Swine influenza virus subtypes identified in 14 European countries from 2010 to 2013. PloS one. 2014;9(12):e115815.

23. Sosa Portugal S, Cortey M, Tello M, Casanovas C, Mesonero-Escuredo S, Barrabés S, et al. Diversity of influenza A viruses retrieved from respiratory disease outbreaks and subclinically infected herds in Spain (2017–2019). Transboundary and Emerging Diseases. 2021;68(2):519–30.

24. Parys A, Vereecke N, Vandoorn E, Theuns S, Van Reeth K. Surveillance and Genomic Characterization of Influenza A and D Viruses in Swine, Belgium and the Netherlands, 2019-2021. Emerg Infect Dis. 2023;29(7):1459–64.

25. Chastagner A, Hervé S, Quéguiner S, Hirchaud E, Lucas P, Gorin S, et al. Genetic and Antigenic Evolution of European Swine Influenza A Viruses of HA-1C (Avian-Like) and HA-1B (Human-Like) Lineages in France from 2000 to 2018. Viruses. 2020;12(11).

26. Hervé S, Garin E, Calavas D, Lecarpentier L, Ngwa-Mbot D, Poliak S, et al. Virological and epidemiological patterns of swine influenza A virus infections in France: Cumulative data from the RESAVIP surveillance network, 2011–2018. Veterinary microbiology. 2019;239:108477.

27. Barr IG, Russell C, Besselaar TG, Cox NJ, Daniels RS, Donis R, et al. WHO recommendations for the viruses used in the 2013–2014 Northern Hemisphere influenza vaccine: Epidemiology, antigenic and genetic characteristics of influenza A(H1N1)pdm09, A(H3N2) and B influenza viruses collected from October 2012 to January 2013. Vaccine. 2014;32(37):4713–25.

28. Centers for Disease Control and Prevention. Antigenic Characterization 2022 [updated 2/12/2022. Available from: https://www.cdc.gov/flu/about/professionals/antigenic.htm.

29. Lewis NS, Russell CA, Langat P, Anderson TK, Berger K, Bielejec F, et al. The global antigenic diversity of swine influenza A viruses. Elife. 2016;5:e12217.

30. IFIP - Institut du Porc. The evolution of pig production in France 2023 [updated 20/07/2023. Available from: https://www.pig333.com/articles/how-has-pig-production-evolved-in-france_19488/.

31. Chastagner A, Bonin E, Fablet C, Quéguiner S, Hirchaud E, Lucas P, et al. Virus persistence in pig herds led to successive reassortment events between swine and human influenza A viruses, resulting in the emergence of a novel triple-reassortant swine influenza virus. Veterinary Research. 2019;50(1):77.

32. Gourreau JM, Kaiser C, Valette M, Douglas AR, Labie J, Aymard M. Isolation of two H1N2 influenza viruses from swine in France. Archives of Virology. 1994;135(3-4):365–82.

33. Brown IH, Harris PA, McCauley JW, Alexander DJ. Multiple genetic reassortment of avian and human influenza A viruses in European pigs, resulting in the emergence of an H1N2 virus of novel genotype. Journal of General Virology. 1998;79(Pt 12):2947–55.

34. Chastagner A, Hervé S, Bonin E, Quéguiner S, Hirchaud E, Henritzi D, et al. Spatio-temporal distribution and evolution of the A/H1N1 2009 pandemic virus in pigs in France from 2009 to 2017: identification of a potential swine-specific lineage. Journal of Virology. 2018.

35. te Beest DE, van Boven M, Hooiveld M, van den Dool C, Wallinga J. Driving Factors of Influenza Transmission in the Netherlands. American journal of epidemiology. 2013;178(9):1469–77.

36. Rose N, Madec F. Occurrence of respiratory disease outbreaks in fattening pigs: relation with the features of a densely and a sparsely populated pig area in France. Vet Res. 2002;33(2):179–90.

37. Schrauwen EJ, Fouchier RA. Host adaptation and transmission of influenza A viruses in mammals. Emerging Microbes & Infections 2014;3(2):e9.

38. Deblanc C, Queguiner S, Gorin S, Richard G, Moro A, Barbier N, et al. Pathogenicity and escape to pre-existing immunity of a new genotype of swine influenza H1N2 virus that emerged in France in 2020. Vet Res. 2024;55(1):65.

39. Chepkwony S, Parys A, Vandoorn E, Stadejek W, Xie J, King J, et al. Genetic and antigenic evolution of H1 swine influenza A viruses isolated in Belgium and the Netherlands from 2014 through 2019. Sci Rep. 2021;11(1):11276.

40. Sooryanarain H, Elankumaran S. Environmental Role in Influenza Virus Outbreaks. Annual Review of Animal Biosciences. 2015;3(1):347–73.

41. The European Commission. European Union Law, Commission Delegated Regulation (EU) 2020/688. 2019.

42. Pol F, Quéguiner S, Gorin S, Deblanc C, Simon G. Validation of commercial real-time RT-PCR kits for detection of influenza A viruses in porcine samples and differentiation of pandemic (H1N1) 2009 virus in pigs. J Virol Methods. 2011;171:241–7.

43. Bonin E, Quéguiner S, Woudstra C, Gorin S, Barbier N, Harder TC, et al. Molecular subtyping of European swine influenza viruses and scaling to high-throughput analysis. Virology journal. 2018;15(1):7.

44. Zhou B, Donnelly ME, Scholes DT, St George K, Hatta M, Kawaoka Y, et al. Single-reaction genomic amplification accelerates sequencing and vaccine production for classical and Swine origin human influenza a viruses. J Virol. 2009;83(19):10309–13.

45. Richard G. gtrichard/influenza_sequences_toolbox: Influenza Sequences Toolbox 0.4: Zenodo; 2024 2024/2/5.

46. Shen W, Le S, Li Y, Hu F. SeqKit: A Cross-Platform and Ultrafast Toolkit for FASTA/Q File Manipulation. PloS one. 2016;11(10):e0163962.

47. Danecek P, Bonfield JK, Liddle J, Marshall J, Ohan V, Pollard MO, et al. Twelve years of SAMtools and BCFtools. GigaScience. 2021;10(2).

48. Katoh K, Standley DM. MAFFT multiple sequence alignment software version 7: improvements in performance and usability. Mol Biol Evol. 2013;30(4):772–80.

49. Minh BQ, Schmidt HA, Chernomor O, Schrempf D, Woodhams MD, von Haeseler A, et al. IQ-TREE 2: New Models and Efficient Methods for Phylogenetic Inference in the Genomic Era. Mol Biol Evol. 2020;37(5):1530–4.

50. Sagulenko P, Puller V, Neher RA. TreeTime: Maximum-likelihood phylodynamic analysis. Virus Evol. 2018;4(1):vex042.

51. Wagih O. ggseqlogo: a versatile R package for drawing sequence logos. Bioinformatics. 2017;33(22):3645–7.

52. Cador C, Hervé S, Andraud M, Gorin S, Paboeuf F, Barbier N, et al. Maternally-derived antibodies do not prevent transmission of swine influenza A virus between pigs. Veterinary research. 2016;47(1):86.

53. OIE. Manual of Diagnostic Tests and Vaccines for Terrestrial Animals, twelfth edition 2023. World Organisation for Animal Health; 2023. p. 1–18.

54. Aragon TJ. epitools: Epidemiology Tools 2020 [Available from: https://CRAN.R-project.org/package=epitools.

