## Supplementary Table 1 for "Major change in swine influenza virus diversity in France owing to emergence and widespread dissemination of a newly introduced H1N2 1C genotype in 2020"

**Supplementary Table 1.** Analytical specificity and sensitivity results and diagnostic ability of the developed $H1_{av}$#E RT-qPCR for discrimination of HA-1C.2.4 of strains from French $H1_{av}N2$#E genotype.

### Analytical specificity of $H1_{av}$#E RT-qPCR

| Strain name | Subtype | Genotype | HA clade | HA accession number | Expected result | HA-geneRT-qPCR (Cq) |
|---|---|---|---|---|---|---|
| A/swine/France/29-200291/2020 | H1huN2 | #B | 1B.1.2.3 | MZ088801 | negative | no Cq |
| A/swine/France/29-200395/2020 | H1huN2 | #B | 1B.1.2.3 | upon request | negative | no Cq |
| A/swine/France/56-200440/2020 | H1huN2 | #B | 1B.1.2.3 | upon request | negative | no Cq |
| A/swine/France/72-200458/2020 | H1avN1 | #A | 1C.2.1 | PP895395 | negative | no Cq |
| A/swine/France/35-190039/2018 | H1avN2 | #A | 1C.2.1 | MT378990 | negative | no Cq |
| A/swine/France/29-200272/2020 | H1avN1 | #A | 1C.2.1 | MZ088793 | negative | no Cq |
| A/swine/France/22-200439/2020 | H1avN1 | #A | 1C.2.1 | PP895832 | negative | no Cq |
| A/swine/France/12-200170/2020 | H1avN1 | #A | 1C.2.1 | MZ088209 | negative | no Cq |
| A/swine/France/81-200304/2020 | H1avN1 | #A | 1C.2.1 | MZ089345 | negative | no Cq |
| A/swine/France/64-200317/2020 | H1avN1 | #A | 1C.2.1 | MZ089249 | negative | no Cq |
| A/swine/France/64-200370/2020 | H1avN1 | #A | 1C.2.1 | PP895411 | negative | no Cq |
| A/swine/France/64-200385/2020 | H1avN1 | #A | 1C.2.1 | PP896392 | negative | no Cq |
| A/swine/France/22-200290/2020 | H1avN1 | #A | 1C.2.1 | MZ088577 | negative | no Cq |
| A/swine/France/56-200313/2020 | H1avN1 | #A | 1C.2.1 | MZ089177 | negative | no Cq |
| A/swine/France/56-180424/2018 | H1avN2 | #F | 1C.2.4 | MT379174 | negative | no Cq |
| A/swine/France/64-150152/2015 | H1avN2 | #E | 1C.2.4 | MT378582 | negative | no Cq |
| A/swine/France/56-200112/2020 | H1avN2 | #E | 1C.2.4 | MZ088185 | positive | 21.58 |
| A/swine/France/22-200113/2020 | H1avN2 | #E | 1C.2.4 | MW048844 | positive | 16.01 |
| A/swine/France/29-200133/2020 | H1avN2 | #E | 1C.2.4 | MZ088721 | positive | 19.42 |
| A/swine/France/53-200134_3/2020 | H1avN2 | #E | 1C.2.4 | MZ089081 | positive | 20.23 |
| A/swine/France/28-200135/2020 | H1avN2 | #E | 1C.2.4 | MZ088625 | positive | 19.97 |
| A/swine/France/35-200154/2020 | H1avN2 | #E | 1C.2.4 | MZ088857 | positive | 21.21 |
| A/swine/France/22-200173/2020 | H1avN2 | #E | 1C.2.4 | MZ088345 | positive | 21.84 |
| A/swine/France/35-200271/2020 | H1avN2 | #E | 1C.2.4 | MZ088905 | positive | 14.55 |
| A/swine/France/35-200275/2020 | H1avN2 | #E | 1C.2.4 | MZ088913 | positive | 15.64 |
| A/swine/France/22-200286 /2020 | H1avN2 | #E | 1C.2.4 | MZ088569 | positive | 15.37 |
| A/swine/France/35-200295 /2020 | H1avN2 | #E | 1C.2.4 | MZ088929 | positive | 16.39 |
| A/swine/France/35-200298/2020 | H1avN2 | #E | 1C.2.4 | MZ088937 | positive | 13.98 |
| A/swine/France/35-200299/2020 | H1avN2 | #E | 1C.2.4 | MZ088945 | positive | 14.58 |
| A/swine/France/35-200300/2020 | H1avN2 | #E | 1C.2.4 | MZ088953 | positive | 13.39 |
| A/swine/France/72-200305/2020 | H1avN2 | #E | 1C.2.4 | MZ089313 | positive | 14.63 |
| A/swine/France/72-200318/2020 | H1avN2 | #E | 1C.2.4 | MZ089321 | positive | 14.56 |
| A/swine/France/22-200322/2020 | H1avN2 | #E | 1C.2.4 | MZ088585 | positive | 14.30 |
| A/swine/France/29-200324/2020 | H1avN2 | #E | 1C.2.4 | MZ088809 | positive | 15.06 |
| A/swine/France/22-200325/2020 | H1avN2 | #E | 1C.2.4 | MZ088593 | positive | 14.89 |
| A/swine/France/35-200332/2020 | H1avN2 | #E | 1C.2.4 | MZ088961 | positive | 14.59 |
| A/swine/France/56-200347/2020 | H1avN2 | #E | 1C.2.4 | MZ089185 | positive | 14.41 |

### Analytical sensitivity of $H1_{av}$#E RT-qPCR

| Strain name | Subtype | Genotype | HA clade | virus stock dilution | M gene RT-qPCR in house (Cq-value) | HA-gene RT-qPCR (Cq-value) |
|---|---|---|---|---|---|---|
| A/swine/France/22-200113/2020 | H1avN2 | #E | 1C.2.4 | 10-1 | 16.71 | 16.01 |
| | | | | 10-2 | 19.86 | 19.57 |
| | | | | 10-3 | 23.38 | 23.46 |
| | | | | 10-4 | 26.94 | 27.02 |
| | | | | 10-5 | 30.40 | 30.53 |
| | | | | 10-6 | 34.89 | 33.95 |
| | | | | 10-7 | no Cq | no Cq |
| | | | | 10-8 | no Cq | no Cq |

### Diagnostic ability of $H1_{av}$#E RT-qPCR

| ID | Subtype | Genotype | HA clade | HA accession number | M gene RT-qPCR* (Cq) | Expected result | HA-gene RT-qPCR (Cq) |
|---|---|---|---|---|---|---|---|
| 190052-1 | H1avN1 | #A | 1C.2.1 | upon request | **31.02** | negative | no Cq |
| 190063-1 | H1avN1 | #A | 1C.2.1 | upon request | **28.0** | negative | no Cq |
| 190080-1 | H1avN1 | #A | 1C.2.1 | MZ088233 | **19.91** | negative | no Cq |
| 190500-3 | H1avN1 | #A | 1C.2.1 | MZ088665 | **26.38** | negative | no Cq |
| 200414-2 | H1avN1 | #A | 1C.2.1 | upon request | **22** | negative | no Cq |
| 190061-3 | H1avN1 | #A | 1C.2.1 | upon request | **28.26** | negative | no Cq |
| 200396-4 | H1avN1 | #A | 1C.2.1 | MZ088633 | **18** | negative | no Cq |
| 200400-1 | H1avN1 | #A | 1C.2.1 | PP895590 | **20.73** | negative | no Cq |
| 190498-1 | H1avN1 | #A | 1C.2.1 | upon request | **30.48** | negative | no Cq |
| 190536-2 | H1avN1 | #A | 1C.2.1 | MZ088281 | **27.11** | negative | no Cq |
| 190583-3 | H1avN1 | #A | 1C.2.1 | MZ088697 | **24.71** | negative | no Cq |
| 190109-1 | H1avN1 | #A | 1C.2.1 | MZ088257 | **23.69** | negative | no Cq |
| 200326-1 | H1avN2 | #A | 1C.2.1 | upon request | **28** | negative | no Cq |
| 190539-2 | H1avN1 | #A | 1C.2.1 | MZ088681 | **25.72** | negative | no Cq |
| 200415-2 | H1avN1 | #A | 1C.2.1 | upon request | **25.26** | negative | no Cq |
| 200474-1 | H1avN1 | #A | 1C.2.1 | PP896512 | **20.34** | negative | no Cq |
| 190476-1 | H1avN1 | #A | 1C.2.1 | MZ089041 | **20.61** | negative | no Cq |
| 190065-1 | H1avN1 | #A | 1C.2.1 | upon request | **30.86** | negative | no Cq |
| 190415-1 | H1avN1 | #A | 1C.2.1 | upon request | **27.77** | negative | no Cq |
| 190212-2 | H1avN1 | #A | 1C.2.1 | MZ088825 | **25.66** | negative | no Cq |
| 180424-2 | H1avN2 | #F | 1C.2.4 | MT379174 | *27.73* | negative | no Cq |
| 190177-2 | H1avN2 | #F | 1C.2.4 | MZ088265 | *20.24* | negative | no Cq |
| 190481-3 | H1avN2 | #F | 1C.2.4 | upon request | *26.25* | negative | no Cq |
| 190491-1 | H1avN2 | #F | 1C.2.4 | MZ088153 | *24.51* | negative | no Cq |
| 200121-3 | H1avN2 | #F | 1C.2.4 | MZ089217 | *27.48* | negative | no Cq |
| 200168-1 | H1avN2 | #F | 1C.2.4 | MZ088329 | *26.19* | negative | no Cq |
| 150152-2 | H1avN2 | #E | 1C.2.4 | MT378582 | *24.89* | negative | no Cq |
| 150242-2 | H1avN2 | #E | 1C.2.4 | MT379342 | *23.78* | negative | no Cq |
| 200050-1 | H1avN2 | #E | 1C.2.4 | MZ088177 | *23.86* | positive | 24.03 |
| 200052-1 | H1avN2 | #E | 1C.2.4 | MZ088849 | *26.22* | positive | 25.68 |
| 200112-1 | H1avN2 | #E | 1C.2.4 | MZ088185 | *23.73* | positive | 24.76 |
| 200113-2 | H1avN2 | #E | 1C.2.4 | MW048844 | *24.86* | positive | 24.34 |
| 200133-3 | H1avN2 | #E | 1C.2.4 | MZ088721 | *24.98* | positive | 25.32 |
| 200135-2 | H1avN2 | #E | 1C.2.4 | MZ088625 | *29.21* | positive | 34.00 |
| 200188-1 | H1avN2 | #E | 1C.2.4 | MZ088737 | *24.01* | positive | 24.51 |
| 200267-3 | H1avN2 | #E | 1C.2.4 | upon request | **35.56** | positive | 31.45 |
| 200268-1 | H1avN2 | #E | 1C.2.4 | upon request | **34.92** | positive | 30.82 |
| 200269-2 | H1avN2 | #E | 1C.2.4 | MZ088537 | **31.18** | positive | 25.16 |
| 200270-1 | H1avN2 | #E | 1C.2.4 | MZ088897 | **30.77** | positive | 25.38 |
| 200314-1 | H1avN2 | #E | 1C.2.4 | upon request | **33.74** | positive | no Cq |
| 200315-2 | H1avN2 | #E | 1C.2.4 | upon request | **23.72** | positive | 27.29 |
| 200319-1 | H1avN2 | #E | 1C.2.4 | upon request | **34.29** | positive | 30.58 |
| 200320-1 | H1avN2 | #E | 1C.2.4 | upon request | **34.37** | positive | 30.44 |
| 200321-1 | H1avN2 | #E | 1C.2.4 | upon request | **34.13** | positive | 29.69 |
| 200323-1 | H1avN2 | #E | 1C.2.4 | upon request | **28.07** | positive | 28.86 |
| 200327-1 | H1avN2 | #E | 1C.2.4 | upon request | **25.06** | positive | 20.98 |
| 200397-3 | H1avN2 | #E | 1C.2.4 | upon request | **28** | positive | 21.87 |
| 200398-6 | H1avN2 | #E | 1C.2.4 | upon request | **24** | positive | 19.55 |
| 200399-1 | H1avN2 | #E | 1C.2.4 | upon request | **24** | positive | 21.73 |
| 200402-2 | H1avN2 | #E | 1C.2.4 | upon request | **27.52** | positive | 25.32 |
| 200404-2 | H1avN2 | #E | 1C.2.4 | upon request | **32.97** | positive | 27.04 |
| 200406-2 | H1avN2 | #E | 1C.2.4 | upon request | **33.74** | positive | 29.15 |
| 200407-4 | H1avN2 | #E | 1C.2.4 | upon request | **22** | positive | 22.04 |
| 200409-4 | H1avN2 | #E | 1C.2.4 | upon request | **33.71** | positive | 26.80 |
| 200410-2 | H1avN2 | #E | 1C.2.4 | upon request | **24.19** | positive | 20.68 |
| 200412-1 | H1avN2 | #E | 1C.2.4 | upon request | **31** | positive | 26.11 |
| 200413-2 | H1avN2 | #E | 1C.2.4 | MZ088601 | **32** | positive | 26.21 |

* M gene RT-qPCR in house in italic, M gene RT-qPCR commercial kits in bold
