## Supplementary Table 2 for "Major change in swine influenza virus diversity in France owing to emergence and widespread dissemination of a newly introduced H1N2 1C genotype in 2020"

**Supplementary Table 2.** Genbank accession numbers for swIAV sequences isolated in France and used in this study

| Strain name | Collection date | Subtype | Genotype | HA clade | PB2 | PB1 | PA | HA | NP | NA | M | NS |
|---|---|---|---|---|---|---|---|---|---|---|---|---|
| A/swine/France/22-190050/2019 | 2019-01-31 | H1avN1 | #A | 1C.2.1 | MZ088222 | MZ088223 | MZ088224 | MZ088225 | MZ088226 | MZ088227 | MZ088228 | MZ088229 |
| A/swine/France/59-190057/2019 | 2019-01-24 | H1huN2 | #C | 1B.1.2.1 | MZ089190 | MZ089191 | MZ089192 | MZ089193 | MZ089194 | MZ089195 | MZ089196 | MZ089197 |
| A/swine/France/50-190058/2019 | 2019-01-22 | H1avN1 | #A | 1C.2.1 | MZ089006 | MZ089007 | MZ089008 | MZ089009 | MZ089010 | MZ089011 | MZ089012 | MZ089013 |
| A/swine/France/56-190060/2019 | 2019-02-05 | H1avN1 | #A | 1C.2.1 | MZ088166 | MZ088167 | MZ088168 | MZ088169 | MZ088170 | MZ088171 | MZ088172 | MZ088173 |
| A/swine/France/72-190071/2019 | 2019-02-11 | H1N1pdm | #A | 1A.3.3.2 | MZ089254 | MZ089255 | MZ089256 | MZ089257 | MZ089258 | MZ089259 | MZ089260 | MZ089261 |
| A/swine/France/17-190072/2019 | 2019-02-06 | H1N1pdm | #A | 1A.3.3.2 | MZ088214 | MZ088215 | MZ088216 | MZ088217 | MZ088218 | MZ088219 | MZ088220 | MZ088221 |
| A/swine/France/22-190080/2019 | 2019-02-20 | H1avN1 | #A | 1C.2.1 | MZ088230 | MZ088231 | MZ088232 | MZ088233 | MZ088234 | MZ088235 | MZ088236 | MZ088237 |
| A/swine/France/64-190081/2019 | 2019-02-19 | H1N1pdm | #A | 1A.3.3.2 | MZ089238 | MZ089239 | MZ089240 | MZ089241 | MZ089242 | MZ089243 | MZ089244 | MZ089245 |
| A/swine/France/40-190086/2019 | 2019-03-01 | H1N1pdm | #A | 1A.3.3.2 | MZ088966 | MZ088967 | MZ088968 | MZ088969 | MZ088970 | MZ088971 | MZ088972 | MZ088973 |
| A/swine/France/22-190096/2019 | 2019-03-12 | H1N1pdm | #A | 1A.3.3.2 | MZ088238 | MZ088239 | MZ088240 | MZ088241 | MZ088242 | MZ088243 | MZ088244 | MZ088245 |
| A/swine/France/22-190099/2019 | 2019-03-12 | H1avN1 | #A | 1C.2.1 | MZ088246 | MZ088247 | MZ088248 | MZ088249 | MZ088250 | MZ088251 | MZ088252 | MZ088253 |
| A/swine/France/35-190107/2019 | 2019-03-21 | H1avN1 | #A | 1C.2.1 | MZ088814 | MZ088815 | MZ088816 | MZ088817 | MZ088818 | MZ088819 | MZ088820 | MZ088821 |
| A/swine/France/22-190109/2019 | 2019-03-21 | H1avN1 | #A | 1C.2.1 | MZ088254 | MZ088255 | MZ088256 | MZ088257 | MZ088258 | MZ088259 | MZ088260 | MZ088261 |
| A/swine/France/50-190161/2019 | 2019-04-12 | H1avN1 | #A | 1C.2.1 | MZ089014 | MZ089015 | MZ089016 | MZ089017 | MZ089018 | MZ089019 | MZ089020 | MZ089021 |
| A/swine/France/59-190174/2019 | 2019-05-02 | H1huN2 | #C | 1B.1.2.1 | MZ089198 | MZ089199 | MZ089200 | MZ089201 | MZ089202 | MZ089203 | MZ089204 | MZ089205 |
| A/swine/France/22-190177/2019 | 2019-04-26 | H1avN2 | #F | 1C.2.4 | MZ088262 | MZ088263 | MZ088264 | MZ088265 | MZ088266 | MZ088267 | MZ088268 | MZ088269 |
| A/swine/France/35-190212/2019 | 2019-05-14 | H1avN1 | #A | 1C.2.1 | MZ088822 | MZ088823 | MZ088824 | MZ088825 | MZ088826 | MZ088827 | MZ088828 | MZ088829 |
| A/swine/France/35-190213/2019 | 2019-05-13 | H1huN2 | #B | 1B.1.2.3 (Δ146-147) | MZ088830 | MZ088831 | MZ088832 | MZ088833 | MZ088834 | MZ088835 | MZ088836 | MZ088837 |
| A/swine/France/56-190288/2019 | 2019-05-06 | H1huN2 | #A | 1B.1.2.3 | MZ089094 | MZ089095 | MZ089096 | MZ089097 | MZ089098 | MZ089099 | MZ089100 | MZ089101 |
| A/swine/France/72-190321/2019 | 2019-05-28 | H1avN1 | #A | 1C.2.1 | MZ089262 | MZ089263 | MZ089264 | MZ089265 | MZ089266 | MZ089267 | MZ089268 | MZ089269 |
| A/swine/France/59-190415/2019 | 2019-07-03 | H1avN1 | #A | 1C.2.1 | upon request | | | | | | | |
| A/swine/France/29-190437/2019 | 2019-07-31 | H1avN1 | #A | 1C.2.1 | MZ088638 | MZ088639 | MZ088640 | MZ088641 | MZ088642 | MZ088643 | MZ088644 | MZ088645 |
| A/swine/France/56-190444/2019 | 2019-08-05 | H1avN1 | #A | 1C.2.1 | MZ089102 | MZ089103 | MZ089104 | MZ089105 | MZ089106 | MZ089107 | MZ089108 | MZ089109 |
| A/swine/France/50-190453/2019 | 2019-08-28 | H1avN1 | #A | 1C.2.1 | MZ089022 | MZ089023 | MZ089024 | MZ089025 | MZ089026 | MZ089027 | MZ089028 | MZ089029 |
| A/swine/France/72-190455/2019 | 2019-08-28 | H1avN1 | #A | 1C.2.1 | MZ089270 | MZ089271 | MZ089272 | MZ089273 | MZ089274 | MZ089275 | MZ089276 | MZ089277 |
| A/swine/France/50-190466/2019 | 2019-09-04 | H1avN1 | #A | 1C.2.1 | MZ089030 | MZ089031 | MZ089032 | MZ089033 | MZ089034 | MZ089035 | MZ089036 | MZ089037 |
| A/swine/France/22-190475/2019 | 2019-09-19 | H1avN1 | #A | 1C.2.1 | MZ088270 | MZ088271 | MZ088272 | MZ088273 | MZ088274 | MZ088275 | MZ088276 | MZ088277 |
| A/swine/France/50-190476/2019 | 2019-09-09 | H1avN1 | #A | 1C.2.1 | MZ089038 | MZ089039 | MZ089040 | MZ089041 | MZ089042 | MZ089043 | MZ089044 | MZ089045 |
| A/swine/France/29-190487/2019 | 2019-06-04 | H1avN1 | #A | 1C.2.1 | MZ088646 | MZ088647 | MZ088648 | MZ088649 | MZ088650 | MZ088651 | MZ088652 | MZ088653 |
| A/swine/France/53-190488/2019 | 2019-06-17 | H1avN1 | #A | 1C.2.1 | MZ089054 | MZ089055 | MZ089056 | MZ089057 | MZ089058 | MZ089059 | MZ089060 | MZ089061 |
| A/swine/France/22-190491/2019 | 2019-08-21 | H1avN2 | #F | 1C.2.4 | MZ088150 | MZ088151 | MZ088152 | MZ088153 | MZ088154 | MZ088155 | MZ088156 | MZ088157 |

| | | | | | | | | | | | | |
|---|---|---|---|---|---|---|---|---|---|---|---|---|
| A/swine/France/29-190493/2019 | 2019-09-19 | H1avN1 | #A | 1C.2.1 | MZ088654 | MZ088655 | MZ088656 | MZ088657 | MZ088658 | MZ088659 | MZ088660 | MZ088661 |
| A/swine/France/22-190500/2019 | 2019-10-08 | H1avN1 | #A | 1C.2.1 | MZ088662 | MZ088663 | MZ088664 | MZ088665 | MZ088666 | MZ088667 | MZ088668 | MZ088669 |
| A/swine/France/29-190501/2019 | 2019-10-15 | H1avN1 | #A | 1C.2.1 | MZ088670 | MZ088671 | MZ088672 | MZ088673 | MZ088674 | MZ088675 | MZ088676 | MZ088677 |
| A/swine/France/22-190536/2019 | 2019-10-24 | H1avN1 | #A | 1C.2.1 | MZ088278 | MZ088279 | MZ088280 | MZ088281 | MZ088282 | MZ088283 | MZ088284 | MZ088285 |
| A/swine/France/29-190539/2019 | 2019-10-31 | H1avN1 | #A | 1C.2.1 | MZ088678 | MZ088679 | MZ088680 | MZ088681 | MZ088682 | MZ088683 | MZ088684 | MZ088685 |
| A/swine/France/53-190540/2019 | 2019-10-29 | H1avN1 | #A | 1C.2.1 | MZ089062 | MZ089063 | MZ089064 | MZ089065 | MZ089066 | MZ089067 | MZ089068 | MZ089069 |
| A/swine/France/50-190548/2019 | 2019-10-24 | H1avN1 | #A | 1C.2.1 | MZ089046 | MZ089047 | MZ089048 | MZ089049 | MZ089050 | MZ089051 | MZ089052 | MZ089053 |
| A/swine/France/85-190552/2019 | 2019-10-30 | H1avN1 | #A | 1C.2.1 | upon request | | | | | | | |
| A/swine/France/59-190568/2019 | 2019-11-12 | H1avN1 | #A | 1C.2.1 | MZ089206 | MZ089207 | MZ089208 | MZ089209 | MZ089210 | MZ089211 | MZ089212 | MZ089213 |
| A/swine/France/22-190570/2019 | 2019-11-19 | H1huN2 | #A | 1B.1.2.3 | MZ088286 | MZ088287 | MZ088288 | MZ088289 | MZ088290 | MZ088291 | MZ088292 | MZ088293 |
| A/swine/France/72-190574/2019 | 2019-11-19 | H1avN1 | #A | 1C.2.1 | MZ089278 | MZ089279 | MZ089280 | MZ089281 | MZ089282 | MZ089283 | MZ089284 | MZ089285 |
| A/swine/France/29-190580/2019 | 2019-11-26 | H1N1pdm | #B | 1A.3.3.2 | MZ088686 | MZ088687 | MZ088688 | MZ088689 | MZ088690 | MZ088691 | MZ088692 | MZ088693 |
| A/swine/France/59-190581/2019 | 2019-11-26 | H1avN1 | #B | 1C.2.2 | MZ082141 | MZ082142 | MZ082143 | MZ082144 | MZ082145 | MZ082146 | MZ082147 | MZ082148 |
| A/swine/France/29-190583/2019 | 2019-12-10 | H1avN1 | #A | 1C.2.1 | MZ088694 | MZ088695 | MZ088696 | MZ088697 | MZ088698 | MZ088699 | MZ088700 | MZ088701 |
| A/swine/France/56-190592/2019 | 2019-08-13 | H1N1pdm | #B | 1A.3.3.2 | MZ089110 | MZ089111 | MZ089112 | MZ089113 | MZ089114 | MZ089115 | MZ089116 | MZ089117 |
| A/swine/France/22-190593/2019 | 2019-10-17 | H1avN2 | #F | 1C.2.4 | upon request | | | | | | | |
| A/swine/France/35-200008/2019 | 2019-12-16 | H1avN1 | #A | 1C.2.1 | MZ088838 | MZ088839 | MZ088840 | MZ088841 | MZ088842 | MZ088843 | MZ088844 | MZ088845 |
| A/swine/France/22-200009_1/2019 | 2019-12-30 | H1avN2 | #A | 1C.2.1 | upon request | | | | | | | |
| A/swine/France/22-200009_2/2019 | 2019-12-30 | H1avN1 | #A | 1C.2.1 | MZ082149 | MZ082150 | MZ082151 | MZ082152 | MZ082153 | MZ082154 | MZ082155 | MZ082156 |
| A/swine/France/44-200018/2020 | 2020-01-21 | H1avN1 | #A | 1C.2.1 | MZ088974 | MZ088975 | MZ088976 | MZ088977 | MZ088978 | MZ088979 | MZ088980 | MZ088981 |
| A/swine/France/01-200024/2020 | 2020-01-17 | H1avN1 | #A | 1C.2.1 | MZ088190 | MZ088191 | MZ088192 | MZ088193 | MZ088194 | MZ088195 | MZ088196 | MZ088197 |
| A/swine/France/56-200025/2020 | 2020-01-29 | H1avN1 | #A | 1C.2.1 | MZ089118 | MZ089119 | MZ089120 | MZ089121 | MZ089122 | MZ089123 | MZ089124 | MZ089125 |
| A/swine/France/29-200045/2020 | 2020-02-12 | H1N1pdm | #B | 1A.3.3.2 | MZ088702 | MZ088703 | MZ088704 | MZ088705 | MZ088706 | MZ088707 | MZ088708 | MZ088709 |
| A/swine/France/59-200047/2020 | 2020-01-31 | H1avN1 | #A | 1C.2.1 | upon request | | | | | | | |
| A/swine/France/56-200050/2020 | 2020-02-13 | H1avN2 | #E | 1C.2.4 | MZ088174 | MZ088175 | MZ088176 | MZ088177 | MZ088178 | MZ088179 | MZ088180 | MZ088181 |
| A/swine/France/35-200052/2020 | 2020-02-19 | H1avN2 | #E | 1C.2.4 | MZ088846 | MZ088847 | MZ088848 | MZ088849 | MZ088850 | MZ088851 | MZ088852 | MZ088853 |
| A/swine/France/05-200058/2020 | 2020-02-21 | H1avN1 | #A | 1C.2.1 | MZ088198 | MZ088199 | MZ088200 | MZ088201 | MZ088202 | MZ088203 | MZ088204 | MZ088205 |
| A/swine/France/22-200062/2020 | 2020-01-30 | H1avN1 | #A | 1C.2.1 | MZ088294 | MZ088295 | MZ088296 | MZ088297 | MZ088298 | MZ088299 | MZ088300 | MZ088301 |
| A/swine/France/24-200063/2020 | 2020-01-14 | H1avN1 | #A | 1C.2.1 | MZ088606 | MZ088607 | MZ088608 | MZ088609 | MZ088610 | MZ088611 | MZ088612 | MZ088613 |
| A/swine/France/82-200069/2020 | 2020-02-28 | H1avN1 | #A | 1C.2.1 | MZ089350 | MZ089351 | MZ089352 | MZ089353 | MZ089354 | MZ089355 | MZ089356 | MZ089357 |
| A/swine/France/80-200094/2020 | 2020-03-09 | H1N1pdm | #A | 1A.3.3.2 | MZ089334 | MZ089335 | MZ089336 | MZ089337 | MZ089338 | MZ089339 | MZ089340 | MZ089341 |
| A/swine/France/44-200095/2020 | 2020-03-05 | H1N1pdm | #A | 1A.3.3.2 | MZ088982 | MZ088983 | MZ088984 | MZ088985 | MZ088986 | MZ088987 | MZ088988 | MZ088989 |
| A/swine/France/22-200096/2020 | 2020-04-15 | H1avN2 | #E | 1C.2.4 | upon request | | | | | | | |
| A/swine/France/35-200099/2020 | 2020-04-20 | H1avN2 | #E | 1C.2.4 | upon request | | | | | | | |

| | | | | | | | | | | | | |
|---|---|---|---|---|---|---|---|---|---|---|---|---|
| A/swine/France/56-200103/2020 | 2020-03-27 | H1avN2 | #E | 1C.2.4 | upon request | | | | | | | |
| A/swine/France/29-200104/2020 | 2020-03-18 | H1avN1 | #A | 1C.2.1 | MZ088710 | MZ088711 | MZ088712 | MZ088713 | MZ088714 | MZ088715 | MZ088716 | MZ088717 |
| A/swine/France/22-200108/2020 | 2020-03-12 | H1N1pdm | #A | 1A.3.3.2 | MZ088302 | MZ088303 | MZ088304 | MZ088305 | MZ088306 | MZ088307 | MZ088308 | MZ088309 |
| A/swine/France/56-200112/2020 | 2020-05-05 | H1avN2 | #E | 1C.2.4 | MZ088182 | MZ088183 | MZ088184 | MZ088185 | MZ088186 | MZ088187 | MZ088188 | MZ088189 |
| A/swine/France/22-200113/2020 | 2020-05-05 | H1avN2 | #E | 1C.2.4 | MW048841 | MW048842 | MW048843 | MW048844 | MW048845 | MW048846 | MW048847 | MW048848 |
| A/swine/France/01-200114/2020 | 2020-02-05 | H1avN1 | #A | 1C.2.1 | upon request | | | | | | | |
| A/swine/France/22-200115/2020 | 2020-02-25 | H1N1pdm | #B | 1A.3.3.2 | MZ088318 | MZ088319 | MZ088320 | MZ088321 | MZ088322 | MZ088323 | MZ088324 | MZ088325 |
| A/swine/France/74-200117/2020 | 2020-03-03 | H1avN1 | #A | 1C.2.1 | MZ089326 | MZ089327 | MZ089328 | MZ089329 | MZ089330 | MZ089331 | MZ089332 | MZ089333 |
| A/swine/France/59-200121/2020 | 2020-03-24 | H1avN2 | #F | 1C.2.4 | MZ089214 | MZ089215 | MZ089216 | MZ089217 | MZ089218 | MZ089219 | MZ089220 | MZ089221 |
| A/swine/France/29-200133/2020 | 2020-05-18 | H1avN2 | #E | 1C.2.4 | MZ088718 | MZ088719 | MZ088720 | MZ088721 | MZ088722 | MZ088723 | MZ088724 | MZ088725 |
| A/swine/France/53-200134_1/2020 | 2020-05-18 | H1avN1 | #A | 1C.2.1 | MZ089070 | MZ089071 | MZ089072 | MZ089073 | MZ089074 | MZ089075 | MZ089076 | MZ089077 |
| A/swine/France/53-200134_3/2020 | 2020-05-18 | H1avN2 | #E | 1C.2.4 | MZ089078 | MZ089079 | MZ089080 | MZ089081 | MZ089082 | MZ089083 | MZ089084 | MZ089085 |
| A/swine/France/28-200135/2020 | 2020-05-19 | H1avN2 | #E | 1C.2.4 | MZ088622 | MZ088623 | MZ088624 | MZ088625 | MZ088626 | MZ088627 | MZ088628 | MZ088629 |
| A/swine/France/44-200139/2020 | 2020-04-20 | H1avN2 | #E | 1C.2.4 | MZ088990 | MZ088991 | MZ088992 | MZ088993 | MZ088994 | MZ088995 | MZ088996 | MZ088997 |
| A/swine/France/35-200154/2020 | 2020-06-03 | H1avN2 | #E | 1C.2.4 | MZ088854 | MZ088855 | MZ088856 | MZ088857 | MZ088858 | MZ088859 | MZ088860 | MZ088861 |
| A/swine/France/53-200155/2020 | 2020-05-26 | H1avN2 | #E | 1C.2.4 | MZ089086 | MZ089087 | MZ089088 | MZ089089 | MZ089090 | MZ089091 | MZ089092 | MZ089093 |
| A/swine/France/22-200168/2020 | 2020-05-04 | H1avN2 | #F | 1C.2.4 | MZ088326 | MZ088327 | MZ088328 | MZ088329 | MZ088330 | MZ088331 | MZ088332 | MZ088333 |
| A/swine/France/56-200169/2020 | 2020-05-04 | H1avN2 | #E | 1C.2.4 | MZ089126 | MZ089127 | MZ089128 | MZ089129 | MZ089130 | MZ089131 | MZ089132 | MZ089133 |
| A/swine/France/12-200170/2020 | 2020-04-29 | H1avN1 | #A | 1C.2.1 | MZ088206 | MZ088207 | MZ088208 | MZ088209 | MZ088210 | MZ088211 | MZ088212 | MZ088213 |
| A/swine/France/22-200171/2020 | 2020-04-15 | H1avN2 | #E | 1C.2.4 | MZ088334 | MZ088335 | MZ088336 | MZ088337 | MZ088338 | MZ088339 | MZ088340 | MZ088341 |
| A/swine/France/22-200173/2020 | 2020-06-09 | H1avN2 | #E | 1C.2.4 | MZ088342 | MZ088343 | MZ088344 | MZ088345 | MZ088346 | MZ088347 | MZ088348 | MZ088349 |
| A/swine/France/22-200175/2020 | 2020-06-09 | H1avN2 | #E | 1C.2.4 | MZ088158 | MZ088159 | MZ088160 | MZ088161 | MZ088162 | MZ088163 | MZ088164 | MZ088165 |
| A/swine/France/22-200176/2020 | 2020-06-08 | H1avN2 | #E | 1C.2.4 | MZ088350 | MZ088351 | MZ088352 | MZ088353 | MZ088354 | MZ088355 | MZ088356 | MZ088357 |
| A/swine/France/22-200177/2020 | 2020-06-11 | H1avN2 | #E | 1C.2.4 | MZ088358 | MZ088359 | MZ088360 | MZ088361 | MZ088362 | MZ088363 | MZ088364 | MZ088365 |
| A/swine/France/29-200185/2020 | 2020-06-12 | H1avN2 | #E | 1C.2.4 | MZ088726 | MZ088727 | MZ088728 | MZ088729 | MZ088730 | MZ088731 | MZ088732 | MZ088733 |
| A/swine/France/29-200188/2020 | 2020-06-16 | H1avN2 | #E | 1C.2.4 | MZ088734 | MZ088735 | MZ088736 | MZ088737 | MZ088738 | MZ088739 | MZ088740 | MZ088741 |
| A/swine/France/72-200189/2020 | 2020-06-15 | H1avN2 | #E | 1C.2.4 | upon request | | | | | | | |
| A/swine/France/22-200193/2020 | 2020-06-19 | H1N1pdm | #B | 1A.3.3.2 | MZ088366 | MZ088367 | MZ088368 | MZ088369 | MZ088370 | MZ088371 | MZ088372 | MZ088373 |
| A/swine/France/56-200201/2020 | 2020-04-23 | H1avN2 | #E | 1C.2.4 | MZ089134 | MZ089135 | MZ089136 | MZ089137 | MZ089138 | MZ089139 | MZ089140 | MZ089141 |
| A/swine/France/35-200203/2020 | 2020-05-26 | H1avN1 | #A | 1C.2.1 | MZ088862 | MZ088863 | MZ088864 | MZ088865 | MZ088866 | MZ088867 | MZ088868 | MZ088869 |
| A/swine/France/72-200205/2020 | 2020-06-05 | H1avN2 | #E | 1C.2.4 | MZ089286 | MZ089287 | MZ089288 | MZ089289 | MZ089290 | MZ089291 | MZ089292 | MZ089293 |
| A/swine/France/35-200206/2020 | 2020-06-09 | H1avN2 | #E | 1C.2.4 | MZ088870 | MZ088871 | MZ088872 | MZ088873 | MZ088874 | MZ088875 | MZ088876 | MZ088877 |
| A/swine/France/22-200207/2020 | 2020-06-12 | H1avN2 | #E | 1C.2.4 | MZ088374 | MZ088375 | MZ088376 | MZ088377 | MZ088378 | MZ088379 | MZ088380 | MZ088381 |
| A/swine/France/22-200208/2020 | 2020-06-01 | H1avN2 | #F | 1C.2.4 | upon request | | | | | | | |

| | | | | | | | | | | | | |
|---|---|---|---|---|---|---|---|---|---|---|---|---|
| A/swine/France/22-200210/2020 | 2020-07-06 | H1avN2 | #E | 1C.2.4 | MZ088382 | MZ088383 | MZ088384 | MZ088385 | MZ088386 | MZ088387 | MZ088388 | MZ088389 |
| A/swine/France/22-200212/2020 | 2020-07-08 | H1avN2 | #E | 1C.2.4 | MZ088390 | MZ088391 | MZ088392 | MZ088393 | MZ088394 | MZ088395 | MZ088396 | MZ088397 |
| A/swine/France/22-200213/2020 | 2020-07-10 | H1avN2 | #E | 1C.2.4 | MZ088398 | MZ088399 | MZ088400 | MZ088401 | MZ088402 | MZ088403 | MZ088404 | MZ088405 |
| A/swine/France/29-200214/2020 | 2020-07-13 | H1avN2 | #E | 1C.2.4 | MZ088742 | MZ088743 | MZ088744 | MZ088745 | MZ088746 | MZ088747 | MZ088748 | MZ088749 |
| A/swine/France/22-200215/2020 | 2020-07-13 | H1avN2 | #E | 1C.2.4 | MZ088406 | MZ088407 | MZ088408 | MZ088409 | MZ088410 | MZ088411 | MZ088412 | MZ088413 |
| A/swine/France/49-200216/2020 | 2020-07-13 | H1avN2 | #E | 1C.2.4 | MZ088998 | MZ088999 | MZ089000 | MZ089001 | MZ089002 | MZ089003 | MZ089004 | MZ089005 |
| A/swine/France/85-200222/2020 | 2020-07-13 | H1avN1 | #A | 1C.2.1 | MZ089358 | MZ089359 | MZ089360 | MZ089361 | MZ089362 | MZ089363 | MZ089364 | MZ089365 |
| A/swine/France/22-200227/2020 | 2020-06-09 | H1avN2 | #E | 1C.2.4 | MZ088414 | MZ088415 | MZ088416 | MZ088417 | MZ088418 | MZ088419 | MZ088420 | MZ088421 |
| A/swine/France/22-200228/2020 | 2020-06-11 | H1avN2 | #E | 1C.2.4 | MZ088422 | MZ088423 | MZ088424 | MZ088425 | MZ088426 | MZ088427 | MZ088428 | MZ088429 |
| A/swine/France/22-200229/2020 | 2020-06-25 | H1avN2 | #E | 1C.2.4 | MZ088430 | MZ088431 | MZ088432 | MZ088433 | MZ088434 | MZ088435 | MZ088436 | MZ088437 |
| A/swine/France/22-200230/2020 | 2020-06-17 | H1avN2 | #F | 1C.2.4 | MZ088438 | MZ088439 | MZ088440 | MZ088441 | MZ088442 | MZ088443 | MZ088444 | MZ088445 |
| A/swine/France/56-200231/2020 | 2020-06-22 | H1avN2 | #E | 1C.2.4 | MZ089142 | MZ089143 | MZ089144 | MZ089145 | MZ089146 | MZ089147 | MZ089148 | MZ089149 |
| A/swine/France/22-200232/2020 | 2020-06-15 | H1avN2 | #A | 1C.2.1 | MZ088446 | MZ088447 | MZ088448 | MZ088449 | MZ088450 | MZ088451 | MZ088452 | MZ088453 |
| A/swine/France/22-200235/2020 | 2020-07-17 | H1avN2 | #E | 1C.2.4 | MZ088454 | MZ088455 | MZ088456 | MZ088457 | MZ088458 | MZ088459 | MZ088460 | MZ088461 |
| A/swine/France/22-200236/2020 | 2020-07-17 | H1avN2 | #E | 1C.2.4 | MZ088462 | MZ088463 | MZ088464 | MZ088465 | MZ088466 | MZ088467 | MZ088468 | MZ088469 |
| A/swine/France/22-200237/2020 | 2020-07-20 | H1avN2 | #E | 1C.2.4 | MZ088470 | MZ088471 | MZ088472 | MZ088473 | MZ088474 | MZ088475 | MZ088476 | MZ088477 |
| A/swine/France/35-200238/2020 | 2020-07-20 | H1avN2 | #E | 1C.2.4 | MZ088878 | MZ088879 | MZ088880 | MZ088881 | MZ088882 | MZ088883 | MZ088884 | MZ088885 |
| A/swine/France/22-200239/2020 | 2020-07-21 | H1avN2 | #E | 1C.2.4 | MZ088478 | MZ088479 | MZ088480 | MZ088481 | MZ088482 | MZ088483 | MZ088484 | MZ088485 |
| A/swine/France/29-200240/2020 | 2020-07-21 | H1huN2 | #A | 1B.1.2.3 | MZ088750 | MZ088751 | MZ088752 | MZ088753 | MZ088754 | MZ088755 | MZ088756 | MZ088757 |
| A/swine/France/22-200241/2020 | 2020-07-23 | H1avN2 | #E | 1C.2.4 | MZ088486 | MZ088487 | MZ088488 | MZ088489 | MZ088490 | MZ088491 | MZ088492 | MZ088493 |
| A/swine/France/22-200242/2020 | 2020-07-29 | H1avN2 | #E | 1C.2.4 | MZ088494 | MZ088495 | MZ088496 | MZ088497 | MZ088498 | MZ088499 | MZ088500 | MZ088501 |
| A/swine/France/72-200244/2020 | 2020-07-22 | H1avN2 | #E | 1C.2.4 | MZ089294 | MZ089295 | MZ089296 | MZ089297 | MZ089298 | MZ089299 | MZ089300 | MZ089301 |
| A/swine/France/22-200249/2020 | 2020-08-06 | H1avN2 | #E | 1C.2.4 | MZ088502 | MZ088503 | MZ088504 | MZ088505 | MZ088506 | MZ088507 | MZ088508 | MZ088509 |
| A/swine/France/59-200250/2020 | 2020-08-05 | H1pdmN2 | #A | 1.A.3.3.2 | MZ089222 | MZ089223 | MZ089224 | MZ089225 | MZ089226 | MZ089227 | MZ089228 | MZ089229 |
| A/swine/France/35-200251/2020 | 2020-08-10 | H1avN2 | #E | 1C.2.4 | MZ088886 | MZ088887 | MZ088888 | MZ088889 | MZ088890 | MZ088891 | MZ088892 | MZ088893 |
| A/swine/France/29-200252/2020 | 2020-08-07 | H1avN2 | #E | 1C.2.4 | MZ088758 | MZ088759 | MZ088760 | MZ088761 | MZ088762 | MZ088763 | MZ088764 | MZ088765 |
| A/swine/France/29-200254/2020 | 2020-08-13 | H1avN2 | #E | 1C.2.4 | MZ088766 | MZ088767 | MZ088768 | MZ088769 | MZ088770 | MZ088771 | MZ088772 | MZ088773 |
| A/swine/France/22-200255/2020 | 2020-08-12 | H1avN2 | #E | 1C.2.4 | MZ088510 | MZ088511 | MZ088512 | MZ088513 | MZ088514 | MZ088515 | MZ088516 | MZ088517 |
| A/swine/France/56-200256/2020 | 2020-08-17 | H1avN2 | #E | 1C.2.4 | MZ089150 | MZ089151 | MZ089152 | MZ089153 | MZ089154 | MZ089155 | MZ089156 | MZ089157 |
| A/swine/France/56-200257/2020 | 2020-08-17 | H1avN2 | #E | 1C.2.4 | MZ089158 | MZ089159 | MZ089160 | MZ089161 | MZ089162 | MZ089163 | MZ089164 | MZ089165 |
| A/swine/France/22-200258/2020 | 2020-08-19 | H1avN2 | #E | 1C.2.4 | MZ088518 | MZ088519 | MZ088520 | MZ088521 | MZ088522 | MZ088523 | MZ088524 | MZ088525 |
| A/swine/France/56-200259/2020 | 2020-08-20 | H1avN2 | #E | 1C.2.4 | MZ089166 | MZ089167 | MZ089168 | MZ089169 | MZ089170 | MZ089171 | MZ089172 | MZ089173 |
| A/swine/France/22-200260/2020 | 2020-08-20 | H1avN2 | #E | 1C.2.4 | MZ088526 | MZ088527 | MZ088528 | MZ088529 | MZ088530 | MZ088531 | MZ088532 | MZ088533 |
| A/swine/France/29-200263/2020 | 2020-06-24 | H1avN2 | #E | 1C.2.4 | MZ088774 | MZ088775 | MZ088776 | MZ088777 | MZ088778 | MZ088779 | MZ088780 | MZ088781 |

| | | | | | | | | | | | | |
|---|---|---|---|---|---|---|---|---|---|---|---|---|
| A/swine/France/29-200264/2020 | 2020-08-26 | H1avN2 | #E | 1C.2.4 | MZ088782 | MZ088783 | MZ088784 | MZ088785 | MZ088786 | MZ088787 | MZ088788 | MZ088789 |
| A/swine/France/56-200265/2020 | 2020-08-24 | H1avN2 | #E | 1C.2.4 | upon request | | | | | | | |
| A/swine/France/56-200268/2020 | 2020-08-28 | H1avN2 | #E | 1C.2.4 | upon request | | | | | | | |
| A/swine/France/22-200269/2020 | 2020-08-31 | H1avN2 | #E | 1C.2.4 | MZ088534 | MZ088535 | MZ088536 | MZ088537 | MZ088538 | MZ088539 | MZ088540 | MZ088541 |
| A/swine/France/35-200270/2020 | 2020-08-31 | H1avN2 | #E | 1C.2.4 | MZ088894 | MZ088895 | MZ088896 | MZ088897 | MZ088898 | MZ088899 | MZ088900 | MZ088901 |
| A/swine/France/35-200271/2020 | 2020-09-01 | H1avN2 | #E | 1C.2.4 | MZ088902 | MZ088903 | MZ088904 | MZ088905 | MZ088906 | MZ088907 | MZ088908 | MZ088909 |
| A/swine/France/29-200272/2020 | 2020-08-31 | H1avN1 | #A | 1C.2.1 | MZ088790 | MZ088791 | MZ088792 | MZ088793 | MZ088794 | MZ088795 | MZ088796 | MZ088797 |
| A/swine/France/35-200275/2020 | 2020-09-03 | H1avN2 | #E | 1C.2.4 | MZ088910 | MZ088911 | MZ088912 | MZ088913 | MZ088914 | MZ088915 | MZ088916 | MZ088917 |
| A/swine/France/72-200276/2020 | 2020-06-29 | H1avN2 | #E | 1C.2.4 | upon request | | | | | | | |
| A/swine/France/72-200277/2020 | 2020-06-30 | H1avN2 | #E | 1C.2.4 | MZ089302 | MZ089303 | MZ089304 | MZ089305 | MZ089306 | MZ089307 | MZ089308 | MZ089309 |
| A/swine/France/22-200278/2020 | 2020-06-29 | H1avN2 | #E | 1C.2.4 | MZ088542 | MZ088543 | MZ088544 | MZ088545 | MZ088546 | MZ088547 | MZ088548 | MZ088549 |
| A/swine/France/35-200279/2020 | 2020-06-29 | H1avN1 | #A | 1C.2.1 | MZ088918 | MZ088919 | MZ088920 | MZ088921 | MZ088922 | MZ088923 | MZ088924 | MZ088925 |
| A/swine/France/22-200281/2020 | 2020-07-12 | H1avN2 | #E | 1C.2.4 | MZ088550 | MZ088551 | MZ088552 | MZ088553 | MZ088554 | MZ088555 | MZ088556 | MZ088557 |
| A/swine/France/22-200283/2020 | 2020-07-15 | H1avN2 | #E | 1C.2.4 | MZ088558 | MZ088559 | MZ088560 | MZ088561 | MZ088562 | MZ088563 | MZ088564 | MZ088565 |
| A/swine/France/59-200284/2020 | 2020-06-30 | H1pdmN2 | #A | 1.A.3.3.2 | MZ089230 | MZ089231 | MZ089232 | MZ089233 | MZ089234 | MZ089235 | MZ089236 | MZ089237 |
| A/swine/France/22-200286 /2020 | 2020-07-02 | H1avN2 | #E | 1C.2.4 | MZ088566 | MZ088567 | MZ088568 | MZ088569 | MZ088570 | MZ088571 | MZ088572 | MZ088573 |
| A/swine/France/35-200287/2020 | 2020-06-23 | H1avN2 | #E | 1C.2.4 | upon request | | | | | | | |
| A/swine/France/22-200290/2020 | 2020-09-08 | H1avN1 | #A | 1C.2.1 | MZ088574 | MZ088575 | MZ088576 | MZ088577 | MZ088578 | MZ088579 | MZ088580 | MZ088581 |
| A/swine/France/29-200291/2020 | 2020-09-09 | H1huN2 | #A | 1B.1.2.3 | MZ088798 | MZ088799 | MZ088800 | MZ088801 | MZ088802 | MZ088803 | MZ088804 | MZ088805 |
| A/swine/France/35-200295 /2020 | 2020-09-14 | H1avN2 | #E | 1C.2.4 | MZ088926 | MZ088927 | MZ088928 | MZ088929 | MZ088930 | MZ088931 | MZ088932 | MZ088933 |
| A/swine/France/35-200298/2020 | 2020-09-10 | H1avN2 | #E | 1C.2.4 | MZ088934 | MZ088935 | MZ088936 | MZ088937 | MZ088938 | MZ088939 | MZ088940 | MZ088941 |
| A/swine/France/35-200299/2020 | 2020-09-14 | H1avN2 | #E | 1C.2.4 | MZ088942 | MZ088943 | MZ088944 | MZ088945 | MZ088946 | MZ088947 | MZ088948 | MZ088949 |
| A/swine/France/35-200300/2020 | 2020-09-17 | H1avN2 | #E | 1C.2.4 | MZ088950 | MZ088951 | MZ088952 | MZ088953 | MZ088954 | MZ088955 | MZ088956 | MZ088957 |
| A/swine/France/81-200304/2020 | 2020-09-10 | H1avN1 | #A | 1C.2.1 | MZ089342 | MZ089343 | MZ089344 | MZ089345 | MZ089346 | MZ089347 | MZ089348 | MZ089349 |
| A/swine/France/72-200305/2020 | 2020-09-16 | H1avN2 | #E | 1C.2.4 | MZ089310 | MZ089311 | MZ089312 | MZ089313 | MZ089314 | MZ089315 | MZ089316 | MZ089317 |
| A/swine/France/56-200313/2020 | 2020-09-22 | H1avN1 | #A | 1C.2.1 | MZ089174 | MZ089175 | MZ089176 | MZ089177 | MZ089178 | MZ089179 | MZ089180 | MZ089181 |
| A/swine/France/22-200315/2020 | 2020-09-18 | H1avN2 | #E | 1C.2.4 | upon request | | | | | | | |
| A/swine/France/64-200317/2020 | 2020-09-18 | H1avN1 | #A | 1C.2.1 | MZ089246 | MZ089247 | MZ089248 | MZ089249 | MZ089250 | MZ089251 | MZ089252 | MZ089253 |
| A/swine/France/72-200318/2020 | 2020-09-28 | H1avN2 | #E | 1C.2.4 | MZ089318 | MZ089319 | MZ089320 | MZ089321 | MZ089322 | MZ089323 | MZ089324 | MZ089325 |
| A/swine/France/22-200319/2020 | 2020-07-10 | H1avN2 | #E | 1C.2.4 | upon request | | | | | | | |
| A/swine/France/22-200320/2020 | 2020-07-12 | H1avN2 | #E | 1C.2.4 | upon request | | | | | | | |
| A/swine/France/22-200321/2020 | 2020-07-02 | H1avN2 | #E | 1C.2.4 | upon request | | | | | | | |
| A/swine/France/22-200322/2020 | 2020-07-22 | H1avN2 | #E | 1C.2.4 | MZ088582 | MZ088583 | MZ088584 | MZ088585 | MZ088586 | MZ088587 | MZ088588 | MZ088589 |
| A/swine/France/29-200323/2020 | 2020-08-02 | H1avN2 | #E | 1C.2.4 | upon request | | | | | | | |

| | | | | | | | | | | | | |
|---|---|---|---|---|---|---|---|---|---|---|---|---|
| A/swine/France/29-200324/2020 | 2020-08-13 | H1avN2 | #E | 1C.2.4 | MZ088806 | MZ088807 | MZ088808 | MZ088809 | MZ088810 | MZ088811 | MZ088812 | MZ088813 |
| A/swine/France/22-200325/2020 | 2020-08-18 | H1avN2 | #E | 1C.2.4 | MZ088590 | MZ088591 | MZ088592 | MZ088593 | MZ088594 | MZ088595 | MZ088596 | MZ088597 |
| A/swine/France/22-200326/2020 | 2020-06-30 | H1avN2 | #A | 1C.2.1 | upon request | | | | | | | |
| A/swine/France/53-200327/2020 | 2020-08-18 | H1avN2 | #E | 1C.2.4 | upon request | | | | | | | |
| A/swine/France/35-200332/2020 | 2020-09-28 | H1avN2 | #E | 1C.2.4 | MZ088958 | MZ088959 | MZ088960 | MZ088961 | MZ088962 | MZ088963 | MZ088964 | MZ088965 |
| A/swine/France/56-200347/2020 | 2020-10-06 | H1avN2 | #E | 1C.2.4 | MZ089182 | MZ089183 | MZ089184 | MZ089185 | MZ089186 | MZ089187 | MZ089188 | MZ089189 |
| A/swine/France/28-200396/2020 | 2020-08-07 | H1avN1 | #A | 1C.2.1 | MZ088630 | MZ088631 | MZ088632 | MZ088633 | MZ088634 | MZ088635 | MZ088636 | MZ088637 |
| A/swine/France/72-200397/2020 | 2020-07-23 | H1avN2 | #E | 1C.2.4 | upon request | | | | | | | |
| A/swine/France/22-200398/2020 | 2020-08-14 | H1avN2 | #E | 1C.2.4 | upon request | | | | | | | |
| A/swine/France/56-200399/2020 | 2020-06-06 | H1avN2 | #E | 1C.2.4 | upon request | | | | | | | |
| A/swine/France/72-200402/2020 | 2020-09-16 | H1avN2 | #E | 1C.2.4 | upon request | | | | | | | |
| A/swine/France/22-200404/2020 | 2020-09-18 | H1avN2 | #E | 1C.2.4 | upon request | | | | | | | |
| A/swine/France/22-200406/2020 | 2020-09-18 | H1avN2 | #E | 1C.2.4 | upon request | | | | | | | |
| A/swine/France/56-200407/2020 | 2020-09-16 | H1avN2 | #E | 1C.2.4 | upon request | | | | | | | |
| A/swine/France/22-200409/2020 | 2020-09-23 | H1avN2 | #E | 1C.2.4 | upon request | | | | | | | |
| A/swine/France/22-200410/2020 | 2020-09-29 | H1avN2 | #E | 1C.2.4 | upon request | | | | | | | |
| A/swine/France/22-200412/2020 | 2020-09-11 | H1avN2 | #E | 1C.2.4 | upon request | | | | | | | |
| A/swine/France/22-200413/2020 | 2020-09-17 | H1avN2 | #E | 1C.2.4 | MZ088598 | MZ088599 | MZ088600 | MZ088601 | MZ088602 | MZ088603 | MZ088604 | MZ088605 |
| A/swine/France/56-200414/2020 | 2020-09-24 | H1avN1 | #A | 1C.2.1 | upon request | | | | | | | |
| A/swine/France/50-200415/2020 | 2020-09-29 | H1avN1 | #A | 1C.2.1 | upon request | | | | | | | |
| A/swine/France/24-200492/2020 | 2020-12-01 | H1avN2 | #E | 1C.2.4 | MZ088614 | MZ088615 | MZ088616 | MZ088617 | MZ088618 | MZ088619 | MZ088620 | MZ088621 |
| A/swine/France/86-210314-2/2021 | 2021-03-28 | H1avN1 | #A | 1C.2.1 | PP895364 | upon request | PP895365 | PP895366 | PP895367 | PP895368 | PP895370 | PP895369 |
| A/swine/France/79-210235-4/2021 | 2021-01-15 | H1avN2 | #E | 1C.2.4 | upon request | PP895371 | PP895372 | PP895373 | PP895374 | PP895375 | PP895377 | PP895376 |
| A/swine/France/72-210233-2/2021 | 2021-01-19 | H1avN2 | #E | 1C.2.4 | upon request | PP895378 | PP895379 | PP895380 | PP895381 | PP895382 | PP895384 | PP895383 |
| A/swine/France/72-200462-2/2020 | 2020-07-02 | H1avN2 | #E | 1C.2.4 | upon request | PP895385 | PP895386 | PP895387 | PP895388 | PP895389 | PP895391 | PP895390 |
| A/swine/France/72-200458-1/2020 | 2020-11-20 | H1avN1 | #A | 1C.2.1 | PP895392 | PP895393 | PP895394 | PP895395 | PP895396 | PP895397 | PP895399 | PP895398 |
| A/swine/France/64-210493-3/2021 | 2021-09-29 | H1avN1 | #A | 1C.2.1 | PP895400 | PP895401 | PP895402 | PP895403 | PP895404 | PP895405 | PP895407 | PP895406 |
| A/swine/France/64-200370-3/2020 | 2020-07-17 | H1avN1 | #A | 1C.2.1 | PP895408 | PP895409 | PP895410 | PP895411 | PP895412 | upon request | PP895414 | PP895413 |
| A/swine/France/59-210424-2/2021 | 2021-07-18 | H1avN1 | #A | 1C.2.1 | PP895415 | PP895416 | upon request | PP895417 | PP895418 | PP895419 | PP895421 | PP895420 |
| A/swine/France/56-220533-3/2022 | 2022-08-31 | H1avN2 | #E | 1C.2.4 | upon request | PP895422 | PP895423 | PP895424 | PP895425 | PP895426 | PP895428 | PP895427 |
| A/swine/France/53-220295-1/2022 | 2022-02-16 | H1avN1 | #D | 1C.2.1 | upon request | PP895429 | PP895430 | PP895431 | PP895432 | PP895433 | PP895435 | PP895434 |
| A/swine/France/53-210237-123/2021 | 2021-01-13 | H1avN2 | #E | 1C.2.4 | upon request | PP895436 | PP895437 | PP895438 | PP895439 | PP895440 | PP895442 | PP895441 |
| A/swine/France/50-220669-3/2022 | 2022-11-18 | H1avN1 | #D | 1C.2.1 | upon request | PP895443 | PP895444 | PP895445 | upon request | PP895446 | PP895448 | PP895447 |
| A/swine/France/50-210417-5/2021 | 2021-07-14 | H1avN2 | #E | 1C.2.4 | upon request | upon request | PP895449 | PP895450 | PP895451 | PP895452 | PP895454 | PP895453 |

| | | | | | | | | | | | | |
|---|---|---|---|---|---|---|---|---|---|---|---|---|
| A/swine/France/50-210307-1/2021 | 2021-03-12 | H1avN2 | #E | 1C.2.4 | upon request | PP895455 | PP895456 | PP895457 | PP895458 | PP895459 | PP895461 | PP895460 |
| A/swine/France/50-200471-3/2020 | 2020-06-30 | H1avN2 | #E | 1C.2.4 | upon request | PP895462 | PP895463 | PP895464 | PP895465 | PP895466 | PP895468 | PP895467 |
| A/swine/France/49-230070-3/2022 | 2022-12-29 | H1avN2 | #E | 1C.2.4 | upon request | PP895469 | PP895470 | PP895471 | PP895472 | PP895473 | PP895475 | PP895474 |
| A/swine/France/44-200512-1/2020 | 2020-07-22 | H1avN2 | #E | 1C.2.4 | upon request | PP895476 | PP895477 | PP895478 | PP895479 | PP895480 | PP895482 | PP895481 |
| A/swine/France/44-200510-2/2020 | 2020-07-23 | H1avN2 | #E | 1C.2.4 | upon request | PP895483 | PP895484 | PP895485 | PP895486 | PP895487 | PP895489 | PP895488 |
| A/swine/France/44-200423-1/2020 | 2020-07-18 | H1avN2 | #E | 1C.2.4 | upon request | PP895490 | PP895491 | PP895492 | PP895493 | PP895494 | PP895496 | PP895495 |
| A/swine/France/35-220447-3/2022 | 2022-06-17 | H1avN1 | #A | 1C.2.1 | upon request | PP895497 | PP895498 | PP895499 | PP895500 | PP895501 | PP895503 | PP895502 |
| A/swine/France/35-210519-2/2021 | 2021-09-01 | H1avN2 | #E | 1C.2.4 | upon request | PP895504 | PP895505 | PP895506 | PP895507 | PP895508 | PP895510 | PP895509 |
| A/swine/France/35-210402-2/2021 | 2021-04-10 | H1avN2 | #E | 1C.2.4 | upon request | PP895511 | PP895512 | PP895513 | PP895514 | PP895515 | PP895517 | PP895516 |
| A/swine/France/35-210129-2/2021 | 2021-01-12 | H1avN2 | #E | 1C.2.4 | upon request | PP895518 | PP895519 | PP895520 | PP895521 | PP895522 | PP895524 | PP895523 |
| A/swine/France/35-210089-3/2021 | 2021-01-18 | H1avN2 | #E | 1C.2.4 | upon request | PP895525 | PP895526 | PP895527 | PP895528 | PP895529 | PP895531 | PP895530 |
| A/swine/France/35-200469-2/2020 | 2020-07-11 | H1avN2 | #E | 1C.2.4 | upon request | PP895532 | PP895533 | PP895534 | PP895535 | PP895536 | PP895538 | PP895537 |
| A/swine/France/29-220506-7/2022 | 2022-07-29 | H1avN2 | #E | 1C.2.4 | upon request | PP895539 | upon request | PP895540 | PP895541 | PP895542 | PP895544 | PP895543 |
| A/swine/France/29-210536-7/2021 | 2021-11-23 | H1avN1 | #A | 1C.2.1 | PP895545 | PP895546 | PP895547 | PP895548 | PP895549 | PP895550 | upon request | PP895551 |
| A/swine/France/29-210451-2/2021 | 2021-06-03 | H1avN2 | #E | 1C.2.4 | upon request | PP895552 | PP895553 | PP895554 | PP895555 | PP895556 | PP895558 | PP895557 |
| A/swine/France/29-210414-4/2021 | 2021-07-03 | H1avN1 | #D | 1C.2.1 | upon request | PP895559 | upon request | PP895560 | PP895561 | PP895562 | PP895564 | PP895563 |
| A/swine/France/29-210413-3/2021 | 2021-07-02 | H1avN1 | #A | 1C.2.1 | upon request | upon request | upon request | PP895565 | PP895566 | PP895567 | PP895569 | PP895568 |
| A/swine/France/29-210403-2/2021 | 2021-04-23 | H1avN2 | #E | 1C.2.4 | upon request | PP895570 | PP895571 | PP895572 | PP895573 | PP895574 | PP895576 | PP895575 |
| A/swine/France/29-210268-6/2021 | 2021-04-13 | H1avN1 | #D | 1C.2.1 | upon request | upon request | upon request | PP895577 | upon request | PP895578 | PP895580 | PP895579 |
| A/swine/France/29-200472-1/2020 | 2020-06-19 | H1avN2 | #E | 1C.2.4 | upon request | PP895581 | PP895582 | PP895583 | PP895584 | PP895585 | PP895587 | PP895586 |
| A/swine/France/29-200400-1/2020 | 2020-09-15 | H1avN1 | #A | 1C.2.1 | upon request | PP895588 | PP895589 | PP895590 | PP895591 | PP895592 | PP895594 | PP895593 |
| A/swine/France/29-200383-3/2020 | 2020-07-19 | H1avN2 | #E | 1C.2.4 | upon request | PP895595 | PP895596 | PP895597 | PP895598 | PP895599 | PP895601 | PP895600 |
| A/swine/France/22-220664-1/2022 | 2022-09-21 | H1avN1 | #? | 1C.2.1 | PP895602 | PP895603 | PP895604 | PP895605 | PP895606 | PP895607 | upon request | PP895608 |
| A/swine/France/22-220545-2/2022 | 2022-09-13 | H1avN1 | #A | 1C.2.1 | PP895609 | PP895610 | PP895611 | PP895612 | PP895613 | PP895614 | upon request | PP895615 |
| A/swine/France/22-220524-1/2022 | 2022-08-10 | H1avN1 | #A | 1C.2.1 | upon request | PP895616 | PP895617 | PP895618 | PP895619 | PP895620 | PP895622 | PP895621 |
| A/swine/France/22-220332-1/2022 | 2022-04-27 | H1avN1 | #A | 1C.2.1 | upon request | PP895623 | PP895624 | PP895625 | PP895626 | PP895627 | PP895629 | PP895628 |
| A/swine/France/22-220174-1/2022 | 2022-01-31 | H1avN1 | #A | 1C.2.1 | upon request | PP895630 | PP895631 | PP895632 | PP895633 | PP895634 | PP895636 | PP895635 |
| A/swine/France/22-210512-5/2021 | 2021-10-22 | H1avN2 | #E | 1C.2.4 | upon request | PP895637 | PP895638 | PP895639 | PP895640 | PP895641 | PP895643 | PP895642 |
| A/swine/France/22-210505-3/2021 | 2021-10-11 | H1avN2 | #E | 1C.2.4 | upon request | PP895644 | PP895645 | PP895646 | PP895647 | PP895648 | PP895650 | PP895649 |
| A/swine/France/22-210481-2/2021 | 2021-08-27 | H1avN1 | #A | 1C.2.1 | upon request | PP895651 | PP895652 | PP895653 | upon request | PP895654 | PP895656 | PP895655 |
| A/swine/France/22-210446-1/2021 | 2021-05-29 | H1avN2 | #E | 1C.2.4 | upon request | PP895657 | PP895658 | PP895659 | PP895660 | PP895661 | PP895663 | PP895662 |
| A/swine/France/22-210427-4/2021 | 2021-08-10 | H1avN1 | #A | 1C.2.1 | upon request | PP895664 | PP895665 | PP895666 | PP895667 | PP895668 | PP895670 | PP895669 |
| A/swine/France/22-210427-3/2021 | 2021-08-10 | H1avN2 | #E | 1C.2.4 | PP895671 | PP895672 | PP895673 | PP895674 | PP895675 | PP895676 | PP895677 | upon request |
| A/swine/France/22-210404-4/2021 | 2021-04-16 | H1avN1 | #A | 1C.2.1 | upon request | upon request | PP895678 | PP895679 | PP895680 | PP895681 | PP895683 | PP895682 |

| | | | | | | | | | | | | |
|---|---|---|---|---|---|---|---|---|---|---|---|---|
| A/swine/France/22-210399-2/2021 | 2021-04-08 | H1avN2 | #E | 1C.2.4 | upon request | upon request | upon request | PP895684 | PP895685 | PP895686 | PP895688 | PP895687 |
| A/swine/France/22-210348-2/2021 | 2021-05-11 | H1avN2 | #E | 1C.2.4 | upon request | PP895689 | PP895690 | PP895691 | PP895692 | PP895693 | PP895695 | PP895694 |
| A/swine/France/22-210323-1/2021 | 2021-05-07 | H1avN1 | #A | 1C.2.1 | upon request | PP895696 | PP895697 | PP895698 | upon request | PP895699 | PP895701 | PP895700 |
| A/swine/France/22-210317-2/2021 | 2021-03-21 | H1avN2 | #E | 1C.2.4 | upon request | PP895702 | PP895703 | PP895704 | PP895705 | PP895706 | PP895708 | PP895707 |
| A/swine/France/22-210309-2/2021 | 2021-02-25 | H1avN1 | #A | 1C.2.1 | upon request | PP895709 | PP895710 | PP895711 | PP895712 | PP895713 | PP895715 | PP895714 |
| A/swine/France/22-210305-1/2021 | 2021-03-17 | H1avN2 | #E | 1C.2.4 | upon request | PP895716 | PP895717 | PP895718 | PP895719 | PP895720 | PP895722 | PP895721 |
| A/swine/France/22-210301-6/2021 | 2021-02-17 | H1avN2 | #E | 1C.2.4 | upon request | upon request | upon request | PP895723 | PP895724 | PP895725 | PP895727 | PP895726 |
| A/swine/France/22-210232-2/2021 | 2021-01-18 | H1avN2 | #E | 1C.2.4 | upon request | PP895728 | PP895729 | PP895730 | PP895731 | PP895732 | PP895734 | PP895733 |
| A/swine/France/22-210061-5/2021 | 2021-01-06 | H1avN1 | #B | 1C.2.1 | upon request | upon request | PP895735 | PP895736 | PP895737 | PP895738 | PP895740 | PP895739 |
| A/swine/France/22-200486-1/2020 | 2020-07-19 | H1avN2 | #E | 1C.2.4 | upon request | upon request | upon request | PP895741 | PP895742 | PP895743 | PP895745 | PP895744 |
| A/swine/France/22-200456-2/2020 | 2020-07-07 | H1avN2 | #E | 1C.2.4 | upon request | PP895746 | PP895747 | PP895748 | PP895749 | PP895750 | PP895752 | PP895751 |
| A/swine/France/22-200454-1/2020 | 2020-07-18 | H1avN2 | #E | 1C.2.4 | upon request | PP895753 | PP895754 | PP895755 | PP895756 | PP895757 | PP895759 | PP895758 |
| A/swine/France/01-220342-1/2022 | 2022-04-22 | H1avN1 | #? | 1C.2.1 | upon request | upon request | upon request | PP895760 | PP895761 | PP895762 | PP895764 | PP895763 |
| A/swine/France/01-220092-2/2022 | 2022-02-16 | H1avN1 | #C | 1C.2.2 | PP895765 | PP895766 | PP895767 | PP895768 | PP895769 | PP895770 | PP895772 | PP895771 |
| A/swine/France/01-220413-3/2022 | 2022-05-29 | H1avN1 | #C | 1C.2.2 | PP895773 | PP895774 | PP895775 | PP895776 | PP895777 | PP895778 | PP895780 | PP895779 |
| A/swine/France/05-220082-3/2022 | 2022-02-11 | H1avN2 | #E | 1C.2.4 | PP895781 | PP895782 | PP895783 | PP895784 | PP895785 | PP895786 | PP895788 | PP895787 |
| A/swine/France/15-220077-2/2022 | 2022-01-26 | H1avN1 | #A | 1C.2.1 | PP895789 | PP895790 | PP895791 | PP895792 | PP895793 | PP895794 | PP895796 | PP895795 |
| A/swine/France/22-200009-2/2019 | 2019-12-30 | H1avN1 | #A | 1C.2.1 | PP895797 | PP895798 | PP895799 | PP895800 | PP895801 | PP895802 | PP895804 | PP895803 |
| A/swine/France/22-200382-2/2020 | 2020-10-15 | H1avN2 | #E | 1C.2.4 | PP895805 | PP895806 | PP895807 | PP895808 | PP895809 | PP895810 | PP895812 | PP895811 |
| A/swine/France/22-200435-1/2020 | 2020-11-09 | H1avN2 | #E | 1C.2.4 | PP895813 | PP895814 | PP895815 | PP895816 | PP895817 | PP895818 | PP895820 | PP895819 |
| A/swine/France/22-200436-2/2020 | 2020-11-09 | H1avN2 | #E | 1C.2.4 | PP895821 | PP895822 | PP895823 | PP895824 | PP895825 | PP895826 | PP895828 | PP895827 |
| A/swine/France/22-200439-3/2020 | 2020-11-10 | H1avN1 | #A | 1C.2.1 | PP895829 | PP895830 | PP895831 | PP895832 | PP895833 | PP895834 | PP895836 | PP895835 |
| A/swine/France/22-200505-2/2020 | 2020-12-10 | H1avN2 | #E | 1C.2.4 | PP895837 | PP895838 | PP895839 | PP895840 | PP895841 | PP895842 | PP895844 | PP895843 |
| A/swine/France/22-210030-2/2020 | 2020-12-29 | H1avN2 | #E | 1C.2.4 | PP895845 | PP895846 | PP895847 | PP895848 | PP895849 | PP895850 | PP895852 | PP895851 |
| A/swine/France/22-210039-1/2020 | 2020-12-02 | H1avN2 | #E | 1C.2.4 | PP895853 | PP895854 | PP895855 | PP895856 | PP895857 | PP895858 | PP895860 | PP895859 |
| A/swine/France/22-210042-1/2020 | 2020-12-09 | H1avN2 | #E | 1C.2.4 | PP895861 | PP895862 | PP895863 | PP895864 | PP895865 | PP895866 | PP895868 | PP895867 |
| A/swine/France/22-210061-2/2021 | 2021-01-06 | H1avN1 | #B | 1C.2.1 | PP895869 | PP895870 | PP895871 | PP895872 | PP895873 | PP895874 | PP895876 | PP895875 |
| A/swine/France/22-210119-1/2020 | 2020-12-14 | H1avN2 | #E | 1C.2.4 | PP895877 | PP895878 | PP895879 | PP895880 | PP895881 | PP895882 | PP895884 | PP895883 |
| A/swine/France/22-210139-3/2021 | 2021-02-10 | H1avN1 | #A | 1C.2.1 | PP895885 | PP895886 | PP895887 | PP895888 | PP895889 | PP895890 | PP895892 | PP895891 |
| A/swine/France/22-210234-2/2021 | 2021-01-20 | H1avN2 | #E | 1C.2.4 | PP895893 | PP895894 | PP895895 | PP895896 | PP895897 | PP895898 | PP895900 | PP895899 |
| A/swine/France/22-210252-1/2021 | 2021-01-27 | H1avN2 | #E | 1C.2.4 | PP895901 | PP895902 | PP895903 | PP895904 | PP895905 | PP895906 | PP895908 | PP895907 |
| A/swine/France/22-210267-4/2021 | 2021-04-10 | H1avN2 | #E | 1C.2.4 | PP895909 | PP895910 | PP895911 | PP895912 | PP895913 | PP895914 | PP895916 | PP895915 |
| A/swine/France/22-210295-4/2021 | 2021-04-28 | H1avN2 | #E | 1C.2.4 | PP895917 | PP895918 | PP895919 | PP895920 | PP895921 | PP895922 | PP895924 | PP895923 |
| A/swine/France/22-210311-1/2021 | 2021-03-20 | H1avN2 | #E | 1C.2.4 | PP895925 | PP895926 | PP895927 | PP895928 | PP895929 | PP895930 | PP895932 | PP895931 |

| | | | | | | | | | | | | |
|---|---|---|---|---|---|---|---|---|---|---|---|---|
| A/swine/France/22-210437-1/2021 | 2021-08-24 | H1avN2 | #E | 1C.2.4 | PP895933 | PP895934 | PP895935 | PP895936 | PP895937 | PP895938 | PP895940 | PP895939 |
| A/swine/France/22-210477-2/2021 | 2021-07-23 | H1avN2 | #E | 1C.2.4 | PP895941 | PP895942 | PP895943 | PP895944 | PP895945 | PP895946 | PP895948 | PP895947 |
| A/swine/France/22-210541-19/2021 | 2021-11-30 | H1avN2 | #E | 1C.2.4 | PP895949 | PP895950 | PP895951 | PP895952 | PP895953 | PP895954 | PP895956 | PP895955 |
| A/swine/France/22-220035-1/2021 | 2021-11-24 | H1avN2 | #E | 1C.2.4 | PP895957 | PP895958 | PP895959 | PP895960 | PP895961 | PP895962 | PP895964 | PP895963 |
| A/swine/France/22-220119-1/2022 | 2022-01-17 | H1avN2 | #E | 1C.2.4 | PP895965 | PP895966 | PP895967 | PP895968 | PP895969 | PP895970 | PP895972 | PP895971 |
| A/swine/France/22-220451-1/2022 | 2022-06-01 | H1avN2 | #E | 1C.2.4 | PP895973 | PP895974 | PP895975 | PP895976 | PP895977 | PP895978 | PP895980 | PP895979 |
| A/swine/France/22-220525-2/2022 | 2022-08-10 | H1avN2 | #E | 1C.2.4 | PP895981 | PP895982 | PP895983 | PP895984 | PP895985 | PP895986 | PP895988 | PP895987 |
| A/swine/France/22-220666-1/2022 | 2022-10-03 | H1avN2 | #E | 1C.2.4 | PP895989 | PP895990 | PP895991 | PP895992 | PP895993 | PP895994 | PP895996 | PP895995 |
| A/swine/France/24-210238-5/2021 | 2021-01-13 | H1avN2 | #H | 1C.2.4 | PP895997 | PP895998 | PP895999 | PP896000 | PP896001 | PP896002 | PP896004 | PP896003 |
| A/swine/France/24-230094-2/2022 | 2022-11-09 | H1avN2 | #I | 1C.2.4 | PP896005 | PP896006 | PP896007 | PP896008 | PP896009 | PP896010 | PP896012 | PP896011 |
| A/swine/France/27-220375-5/2022 | 2022-03-12 | H1avN2 | #E | 1C.2.4 | PP896013 | PP896014 | PP896015 | PP896016 | PP896017 | PP896018 | PP896020 | PP896019 |
| A/swine/France/28-210405-2/2021 | 2021-04-21 | H1avN2 | #E | 1C.2.4 | PP896021 | PP896022 | PP896023 | PP896024 | PP896025 | PP896026 | PP896028 | PP896027 |
| A/swine/France/29-200507-2/2020 | 2020-12-15 | H1avN2 | #E | 1C.2.4 | PP896029 | PP896030 | PP896031 | PP896032 | PP896033 | PP896034 | PP896036 | PP896035 |
| A/swine/France/29-210041-1/2020 | 2020-12-02 | H1avN2 | #E | 1C.2.4 | PP896037 | PP896038 | PP896039 | PP896040 | PP896041 | PP896042 | PP896044 | PP896043 |
| A/swine/France/29-210515-1/2021 | 2021-10-29 | H1avN2 | #E | 1C.2.4 | PP896045 | PP896046 | PP896047 | PP896048 | PP896049 | PP896050 | PP896052 | PP896051 |
| A/swine/France/29-220033-8/2022 | 2022-01-18 | H1avN2 | #E | 1C.2.4 | PP896053 | PP896054 | PP896055 | PP896056 | PP896057 | PP896058 | PP896060 | PP896059 |
| A/swine/France/29-220173-1/2022 | 2022-01-18 | H1avN2 | #E | 1C.2.4 | PP896061 | PP896062 | PP896063 | PP896064 | PP896065 | PP896066 | PP896068 | PP896067 |
| A/swine/France/29-220615-2/2022 | 2022-09-23 | H1avN2 | #E | 1C.2.4 | PP896069 | PP896070 | PP896071 | PP896072 | PP896073 | PP896074 | PP896076 | PP896075 |
| A/swine/France/29-220685-3/2022 | 2022-12-01 | H1avN2 | #E | 1C.2.4 | PP896077 | PP896078 | PP896079 | PP896080 | PP896081 | PP896082 | PP896084 | PP896083 |
| A/swine/France/29-220698-2/2022 | 2022-11-08 | H1avN2 | #E | 1C.2.4 | PP896085 | PP896086 | PP896087 | PP896088 | PP896089 | PP896090 | PP896092 | PP896091 |
| A/swine/France/33-220648-3/2022 | 2022-10-03 | H1avN1 | #A | 1C.2.1 | PP896093 | PP896094 | PP896095 | PP896096 | PP896097 | PP896098 | PP896100 | PP896099 |
| A/swine/France/33-220699-2/2022 | 2022-11-03 | H1avN1 | #A | 1C.2.1 | PP896101 | PP896102 | PP896103 | PP896104 | PP896105 | PP896106 | PP896108 | PP896107 |
| A/swine/France/35-200425-2/2020 | 2020-11-02 | H1avN2 | #E | 1C.2.4 | PP896109 | PP896110 | PP896111 | PP896112 | PP896113 | PP896114 | PP896116 | PP896115 |
| A/swine/France/35-200437-1/2020 | 2020-11-10 | H1avN2 | #E | 1C.2.4 | PP896117 | PP896118 | PP896119 | PP896120 | PP896121 | PP896122 | PP896124 | PP896123 |
| A/swine/France/35-200438-2/2020 | 2020-11-10 | H1avN2 | #E | 1C.2.4 | PP896125 | PP896126 | PP896127 | PP896128 | PP896129 | PP896130 | PP896132 | PP896131 |
| A/swine/France/35-200470-2/2020 | 2020-11-25 | H1avN2 | #E | 1C.2.4 | PP896133 | PP896134 | PP896135 | PP896136 | PP896137 | PP896138 | PP896140 | PP896139 |
| A/swine/France/35-200508-2/2020 | 2020-12-10 | H1avN2 | #E | 1C.2.4 | PP896141 | PP896142 | PP896143 | PP896144 | PP896145 | PP896146 | PP896148 | PP896147 |
| A/swine/France/35-210040-1/2020 | 2020-11-06 | H1avN2 | #E | 1C.2.4 | PP896149 | PP896150 | PP896151 | PP896152 | PP896153 | PP896154 | PP896156 | PP896155 |
| A/swine/France/35-210062-6/2021 | 2021-01-11 | H1avN2 | #E | 1C.2.4 | PP896157 | PP896158 | PP896159 | PP896160 | PP896161 | PP896162 | PP896164 | PP896163 |
| A/swine/France/35-210140-1/2021 | 2021-02-14 | H1avN2 | #E | 1C.2.4 | PP896165 | PP896166 | PP896167 | PP896168 | PP896169 | PP896170 | PP896172 | PP896171 |
| A/swine/France/35-210165-4/2021 | 2021-02-21 | H1avN2 | #E | 1C.2.4 | PP896173 | PP896174 | PP896175 | PP896176 | PP896177 | PP896178 | PP896180 | PP896179 |
| A/swine/France/35-210254-2/2021 | 2021-01-27 | H1avN2 | #E | 1C.2.4 | PP896181 | PP896182 | PP896183 | PP896184 | PP896185 | PP896186 | PP896188 | PP896187 |
| A/swine/France/35-210370-1/2021 | 2021-05-21 | H1avN2 | #E | 1C.2.4 | PP896189 | PP896190 | PP896191 | PP896192 | PP896193 | PP896194 | PP896196 | PP896195 |
| A/swine/France/35-210500-1/2021 | 2021-08-19 | H1avN1 | #A | 1C.2.1 | PP896197 | PP896198 | PP896199 | PP896200 | PP896201 | PP896202 | PP896204 | PP896203 |

| | | | | | | | | | | | | |
|---|---|---|---|---|---|---|---|---|---|---|---|---|
| A/swine/France/35-210501-6/2021 | 2021-10-04 | H1avN2 | #E | 1C.2.4 | PP896205 | PP896206 | PP896207 | PP896208 | PP896209 | PP896210 | PP896212 | PP896211 |
| A/swine/France/35-210518-1/2021 | 2021-09-07 | H1avN1 | #A | 1C.2.1 | PP896213 | PP896214 | PP896215 | PP896216 | PP896217 | PP896218 | PP896220 | PP896219 |
| A/swine/France/35-220448-3/2022 | 2022-06-28 | H1avN2 | #E | 1C.2.4 | PP896221 | PP896222 | PP896223 | PP896224 | PP896225 | PP896226 | PP896228 | PP896227 |
| A/swine/France/37-220188-1/2022 | 2022-03-15 | H1avN2 | #E | 1C.2.4 | PP896229 | PP896230 | PP896231 | PP896232 | PP896233 | PP896234 | PP896236 | PP896235 |
| A/swine/France/44-200384-2/2020 | 2020-10-19 | H1avN2 | #E | 1C.2.4 | PP896237 | PP896238 | PP896239 | PP896240 | PP896241 | PP896242 | PP896244 | PP896243 |
| A/swine/France/44-200460-3/2020 | 2020-11-16 | H1avN2 | #E | 1C.2.4 | PP896245 | PP896246 | PP896247 | PP896248 | PP896249 | PP896250 | PP896252 | PP896251 |
| A/swine/France/44-220189-1/2022 | 2022-03-23 | H1avN2 | #E | 1C.2.4 | PP896253 | PP896254 | PP896255 | PP896256 | PP896257 | PP896258 | PP896260 | PP896259 |
| A/swine/France/49-200353-3/2020 | 2020-10-06 | H1avN2 | #E | 1C.2.4 | PP896261 | PP896262 | PP896263 | PP896264 | PP896265 | PP896266 | PP896268 | PP896267 |
| A/swine/France/49-200490-3/2020 | 2020-11-27 | H1avN2 | #E | 1C.2.4 | PP896269 | PP896270 | PP896271 | PP896272 | PP896273 | PP896274 | PP896276 | PP896275 |
| A/swine/France/49-210490-1/2021 | 2021-09-21 | H1avN2 | #E | 1C.2.4 | PP896277 | PP896278 | PP896279 | PP896280 | PP896281 | PP896282 | PP896284 | PP896283 |
| A/swine/France/49-220101-1/2022 | 2022-02-22 | H1avN2 | #E | 1C.2.4 | PP896285 | PP896286 | PP896287 | PP896288 | PP896289 | PP896290 | PP896292 | PP896291 |
| A/swine/France/50-220084-2/2022 | 2022-02-07 | H1avN1 | #A | 1C.2.1 | PP896293 | PP896294 | PP896295 | PP896296 | PP896297 | PP896298 | PP896300 | PP896299 |
| A/swine/France/53-210090-4/2021 | 2021-01-20 | H1avN2 | #E | 1C.2.4 | PP896301 | PP896302 | PP896303 | PP896304 | PP896305 | PP896306 | PP896308 | PP896307 |
| A/swine/France/53-210091-2/2021 | 2021-01-21 | H1avN2 | #E | 1C.2.4 | PP896309 | PP896310 | PP896311 | PP896312 | PP896313 | PP896314 | PP896316 | PP896315 |
| A/swine/France/53-210426-3/2021 | 2021-07-30 | H1avN2 | #E | 1C.2.4 | PP896317 | PP896318 | PP896319 | PP896320 | PP896321 | PP896322 | PP896324 | PP896323 |
| A/swine/France/53-230071-5/2022 | 2022-11-22 | H1avN2 | #E | 1C.2.4 | PP896325 | PP896326 | PP896327 | PP896328 | PP896329 | PP896330 | PP896332 | PP896331 |
| A/swine/France/56-200441-3/2020 | 2020-11-12 | H1avN2 | #E | 1C.2.4 | PP896333 | PP896334 | PP896335 | PP896336 | PP896337 | PP896338 | PP896340 | PP896339 |
| A/swine/France/56-210105-1/2021 | 2021-01-15 | H1avN1 | #A | 1C.2.1 | PP896341 | PP896342 | PP896343 | PP896344 | PP896345 | PP896346 | PP896348 | PP896347 |
| A/swine/France/56-210124-1/2021 | 2021-01-04 | H1avN1 | #A | 1C.2.1 | PP896349 | PP896350 | PP896351 | PP896352 | PP896353 | PP896354 | PP896356 | PP896355 |
| A/swine/France/56-210255-1/2021 | 2021-02-01 | H1avN2 | #E | 1C.2.4 | PP896357 | PP896358 | PP896359 | PP896360 | PP896361 | PP896362 | PP896364 | PP896363 |
| A/swine/France/56-220651-2/2022 | 2022-11-07 | H1avN2 | #E | 1C.2.4 | PP896365 | PP896366 | PP896367 | PP896368 | PP896369 | PP896370 | PP896372 | PP896371 |
| A/swine/France/59-220437-3/2022 | 2022-06-10 | H1avN1 | #C | 1C.2.2 | PP896373 | PP896374 | PP896375 | PP896376 | PP896377 | PP896378 | PP896380 | PP896379 |
| A/swine/France/59-220536-3/2022 | 2022-06-01 | H1avN1 | #C | 1C.2.2 | PP896381 | PP896382 | PP896383 | PP896384 | PP896385 | PP896386 | PP896388 | PP896387 |
| A/swine/France/64-200385-2/2020 | 2020-10-21 | H1avN1 | #A | 1C.2.1 | PP896389 | PP896390 | PP896391 | PP896392 | PP896393 | PP896394 | PP896396 | PP896395 |
| A/swine/France/64-220661-4/2022 | 2022-11-21 | H1avN1 | #A | 1C.2.1 | PP896397 | PP896398 | PP896399 | PP896400 | PP896401 | PP896402 | PP896404 | PP896403 |
| A/swine/France/72-200461-1/2020 | 2020-11-09 | H1avN2 | #E | 1C.2.4 | PP896405 | PP896406 | PP896407 | PP896408 | PP896409 | PP896410 | PP896412 | PP896411 |
| A/swine/France/72-210053-6/2021 | 2021-01-04 | H1avN2 | #E | 1C.2.4 | PP896413 | PP896414 | PP896415 | PP896416 | PP896417 | PP896418 | PP896420 | PP896419 |
| A/swine/France/72-210055-1/2021 | 2021-01-11 | H1avN2 | #E | 1C.2.4 | PP896421 | PP896422 | PP896423 | PP896424 | PP896425 | PP896426 | PP896428 | PP896427 |
| A/swine/France/72-210508-1/2021 | 2021-10-12 | H1avN2 | #E | 1C.2.4 | PP896429 | PP896430 | PP896431 | PP896432 | PP896433 | PP896434 | PP896436 | PP896435 |
| A/swine/France/72-210509-4/2021 | 2021-10-19 | H1avN2 | #E | 1C.2.4 | PP896437 | PP896438 | PP896439 | PP896440 | PP896441 | PP896442 | PP896444 | PP896443 |
| A/swine/France/72-220016-1/2021 | 2021-12-27 | H1avN2 | #E | 1C.2.4 | PP896445 | PP896446 | PP896447 | PP896448 | PP896449 | PP896450 | PP896452 | PP896451 |
| A/swine/France/72-220017-3/2022 | 2022-01-03 | H1avN2 | #E | 1C.2.4 | PP896453 | PP896454 | PP896455 | PP896456 | PP896457 | PP896458 | PP896460 | PP896459 |
| A/swine/France/72-220656-1/2022 | 2022-11-08 | H1avN1 | #B | 1C.2.1 | PP896461 | PP896462 | PP896463 | PP896464 | PP896465 | PP896466 | PP896468 | PP896467 |
| A/swine/France/72-220656-3/2022 | 2022-11-08 | H1avN1 | #B | 1C.2.1 | PP896469 | PP896470 | PP896471 | PP896472 | PP896473 | PP896474 | PP896476 | PP896475 |

| | | | | | | | | | | | | |
|---|---|---|---|---|---|---|---|---|---|---|---|---|
| A/swine/France/72-230004-2/2022 | 2022-12-20 | H1avN1 | #B | 1C.2.1 | PP896477 | PP896478 | PP896479 | PP896480 | PP896481 | PP896482 | PP896484 | PP896483 |
| A/swine/France/76-210447-1/2021 | 2021-05-29 | H1avN2 | #E | 1C.2.4 | PP896485 | PP896486 | PP896487 | PP896488 | PP896489 | PP896490 | PP896492 | PP896491 |
| A/swine/France/79-210184-5/2021 | 2021-03-03 | H1avN1 | #A | 1C.2.1 | PP896493 | PP896494 | PP896495 | PP896496 | PP896497 | PP896498 | PP896500 | PP896499 |
| A/swine/France/80-220527-2/2022 | 2022-08-12 | H1avN1 | #C | 1C.2.2 | PP896501 | PP896502 | PP896503 | PP896504 | PP896505 | PP896506 | PP896508 | PP896507 |
| A/swine/France/81-200474-1/2020 | 2020-10-15 | H1avN1 | #A | 1C.2.1 | PP896509 | PP896510 | PP896511 | PP896512 | PP896513 | PP896514 | PP896516 | PP896515 |
| A/swine/France/81-210264-4/2021 | 2021-03-03 | H1avN2 | #E | 1C.2.4 | PP896517 | PP896518 | PP896519 | PP896520 | PP896521 | PP896522 | PP896524 | PP896523 |
| A/swine/France/81-220129-1/2022 | 2022-03-03 | H1avN1 | #A | 1C.2.1 | PP896525 | PP896526 | PP896527 | PP896528 | PP896529 | PP896530 | PP896532 | PP896531 |
| A/swine/France/85-210258-1/2021 | 2021-03-24 | H1avN1 | #A | 1C.2.1 | PP896533 | PP896534 | PP896535 | PP896536 | PP896537 | PP896538 | PP896540 | PP896539 |
| A/swine/France/86-210313-1/2021 | 2021-03-17 | H1avN1 | #A | 1C.2.1 | PP896541 | PP896542 | PP896543 | PP896544 | PP896545 | PP896546 | PP896548 | PP896547 |
